# Learning stochastic reaction-diffusion models from limited data using spatiotemporal features

**DOI:** 10.1101/2024.10.02.616367

**Authors:** Bedri Abubaker-Sharif, Tatsat Banerjee, Peter N. Devreotes, Pablo A. Iglesias

## Abstract

Pattern-forming stochastic systems arise throughout biology, with dynamic molecular waves observed in biochemical networks regulating critical cellular processes. Modeling these reaction-diffusion systems using handcrafted stochastic partial differential equations (PDEs) requires extensive trial-and-error tuning. Data-driven approaches for improved modeling are needed but have been hindered by data scarcity and noise. Here, we present a solution to the inverse problem of learning stochastic reaction-diffusion models from limited data by optimizing two spatiotemporal features: (1) stochastic dynamics and (2) spatiotemporal patterns. Combined with sparsity enforcement, this method identifies novel activator-inhibitor models with interpretable structure. We demonstrate robust learning from simulations of excitable systems with varying data scarcity, as well as noisy live-cell imaging data with low temporal resolution and a single observed biomolecule. This generalizable approach to learning governing stochastic PDEs enhances our ability to model and understand complex spatiotemporal systems from limited, real-world data.

**Teaser:** This machine learning approach estimates stochastic PDE models using noisy, scarce data from simulations and live-cell imaging.

## Introduction

Stochastic reaction-diffusion systems, in which locally interacting species diffuse in space, have provided a framework for understanding a range of complex phenomena, from molecular scale processes in biology (*1–4*) and physics (*5–7*), to geographical scale events in ecology (*8, 9*) and epidemiology (*10, 11*). With dynamics that are stochastic and spatially-distributed, the system can spontaneously generate a wide array of patterns, such as traveling waves and oscillations, that govern system behavior. In cell biology, for example, the formation of dynamic molecular patterns plays a crucial role in regulating physiological processes and behavior in various cell types. One of the most salient examples of reaction-diffusion systems with emergent properties involves excitable systems, a subset of dynamical systems that display common fundamental characteristics such as nonlinearity, all-or-none responses, and refractory periods. Since Hodgkin and Huxley’s groundbreaking discovery of the mechanisms underlying nerve cell activity, the theory of excitability has been applied across various scientific disciplines (*12–22*). Mathematical models based on stochastic reaction-diffusion equations (SRDEs), a class of stochastic partial differential equations (PDEs), have greatly aided the understanding of these complex systems. However, the mathematical structures are manually set based on knowledge and require extremely difficult parameter tuning. A significant challenge remains in developing data-driven approaches capable of learning new and more accurate models describing spatially-distributed stochastic systems, especially from noisy and scarce real-world datasets. Such algorithms would provide powerful tools for modeling and predicting complex spatiotemporal processes in biology and other scientific disciplines.

Modeling the biochemical network in cells has generally focused on approximating the patterns observed in experiments. In many cell types, extensive imaging data indicates that biomolecules can react and diffuse as a coordinated excitable system, self-organizing into dynamic domains in the cell cortex that propagate as waves (*23–32*). For example, in migrating cells, “front” network components, such as Ras and PIP3, generally form the leading edge of the wave, while “refractory” components, such as PKBA potentially, accumulate to varying degrees behind the wave front; other “back” components appear in the inactive regions of the cell cortex both ahead of and behind waves (*33–38*). Interestingly, these zones approximate the general patterns observed in activator-inhibitor models such as the FitzHugh-Nagumo (FHN) model with a single positive feedback loop and a single delayed negative feedback loop (*22*). Previous studies employing activator-inhibitor models have been used extensively to model a wide array of wave dynamics (*39–47*). These conceptual models are simulated as stochastic partial differential equations (PDEs), with spatial Langevin-type dynamics. Model structure and parameters are adjusted manually based on experience and observation. Nonetheless, this iterative process is often time-intensive and does not facilitate the automated discovery of new models with potentially more insightful structures that better capture the dynamics and patterns observed experimentally in cells.

To address the inverse problem of learning nonlinear models from data, several approaches have been developed in the machine learning literature (*48,49*). However, many of these techniques focus on deterministic PDE models (*50–56*) or spatially independent stochastic differential equation (SDE) models (*57, 58*). In recent years, there has also been an advancement of Fokker-Planck (FP)-based approaches for learning the structure and parameters of spatially-independent SDEs (*58–63*). In cell migration research, some studies have adapted these FP-based approaches to estimate models of cell motion in chemical gradients (*64*). However, to our knowledge, none of these approaches have explored the types of models and datasets relevant to biological stochastic reaction-diffusion systems. Microscopy data for these cellular systems have unique limitations, such as being stochastic in time and space, highly dynamic on the order of seconds, and often limited in observation to one or two model species. Generally, current approaches overlook stochastic PDEs, rely on large datasets with spatiotemporal resolution exceeding system dynamics by orders of magnitude, and require observations being available for all modeled variables.

In this study, we present a novel proof-of-concept solution to learn stochastic PDE models from limited experimental data. Our approach involves extracting two distinct but complementary spatiotemporal features from data: 1) microscale stochastic dynamics in spatial systems, estimated efficiently using a novel method by considering a Laplacian-extended state space, and 2) macroscale spatiotemporal patterns (wave features), estimated using probability distributions and autocorrelations. We then apply various machine learning approaches and show the effectiveness of optimizing for these features to learn stochastic reaction-diffusion equation (SRDE) models from complex datasets over a range of data resolutions.

To optimize stochastic dynamics, we develop Kramers-Moyal optimization by extending previous FP-based algorithms (*58, 59*) to spatially-distributed systems. In this algorithm, we learn stochastic models by fitting the deterministic (“drift”) and stochastic (“diffusivity”) terms of stochastic PDEs to corresponding data-derived conditional expectations, also known as Kramers-Moyal (KM) averages. To learn model structure together with model parameters, we enforce sparsity over a library of functional terms using a reverse greedy search algorithm (*58, 59*), and estimate optimal sparsity using Bayesian information criterion (BIC) instead of the cross-validation techniques used in previous work. Additionally, unlike past studies which did not consider spatially-distributed excitable models, we adapt these algorithms to learn SRDE models with polynomial and rational reaction terms.

To optimize wave features, we implement a novel Monte Carlo strategy to tune model parameters. For initialization, we use initial parameters obtained from either KM regression when available, or from previous knowledge. At each iteration, we generate a reconstruction dataset consisting of multiple simulations, and then compute various spatiotemporal metrics such as wave activity, number of firings, and autocorrelation functions. We then optimize a custom objective function based on the relative error in spatiotemporal patterns between the training dataset and the reconstruction dataset in order to estimate optimal reaction-diffusion model parameters. To simultaneously optimize both stochastic dynamics and wave features for particularly complex datasets, we incorporate Kramers-Moyal metrics in the custom objective function.

Our dual strategy here provides an efficient data-driven approach to learning governing equations for spatially-distributed, stochastic systems. We apply the approach on training datasets of excitable systems obtained from both *in silico* simulations and experimental live-cell imaging. For simulations, we focus on two-component, activator-inhibitor based SRDE models derived from the FitzHugh-Nagumo (FHN) model and a biochemically adapted FHN variant (FR model). These models mimic the type of waves observed in spatiotemporal data acquired from confocal and total internal reflection fluorescence (TIRF) microscopy of cells. To validate our method in the context of some of the unique challenges of learning from experimental data, we test a range of simulation data contexts involving 1) 1D and 2D spatially-distributed, stochastic systems, 2) highly nonlinear dynamic processes with waves lasting approximately 10 to 60 seconds, 3) known or unknown model structure, 4) temporal subsampling every 1 or 4 seconds, and 5) observing only one model species. We ultimately demonstrate the effectiveness of our approach on cell microscopy data by learning a stochastic PDE model from a single replicate of experimental training data with low temporal resolution of 7 seconds, and only a single observed species, PIP3. Overall, this study provides a robust, generalizable machine learning approach for estimating stochastic PDE models of complex systems directly from limited spatiotemporal data.

## Results

### Stochastic reaction-diffusion models

The overall strategy employed for learning stochastic reaction-diffusion models from data is illustrated in Figure 1. Stochastic simulations were first generated based on two distinct models of excitable systems: the classical FitzHugh-Nagumo (FHN) model (*65, 66*), and a modified version with “front” (F) and “refractory” (R) species for biochemical processes, which we refer to as the FR model (*39, 67*). These activator-inhibitor models (activator *x_A_*, inhibitor *x_l_*) were implemented with spatial Langevin dynamics as stochastic partial differential equations (PDEs) (Fig. 1a). In Itô differential form, we have the following coupled equations:

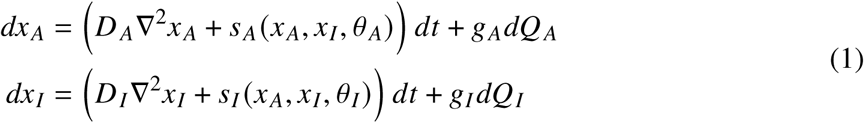

**Figure 1:**
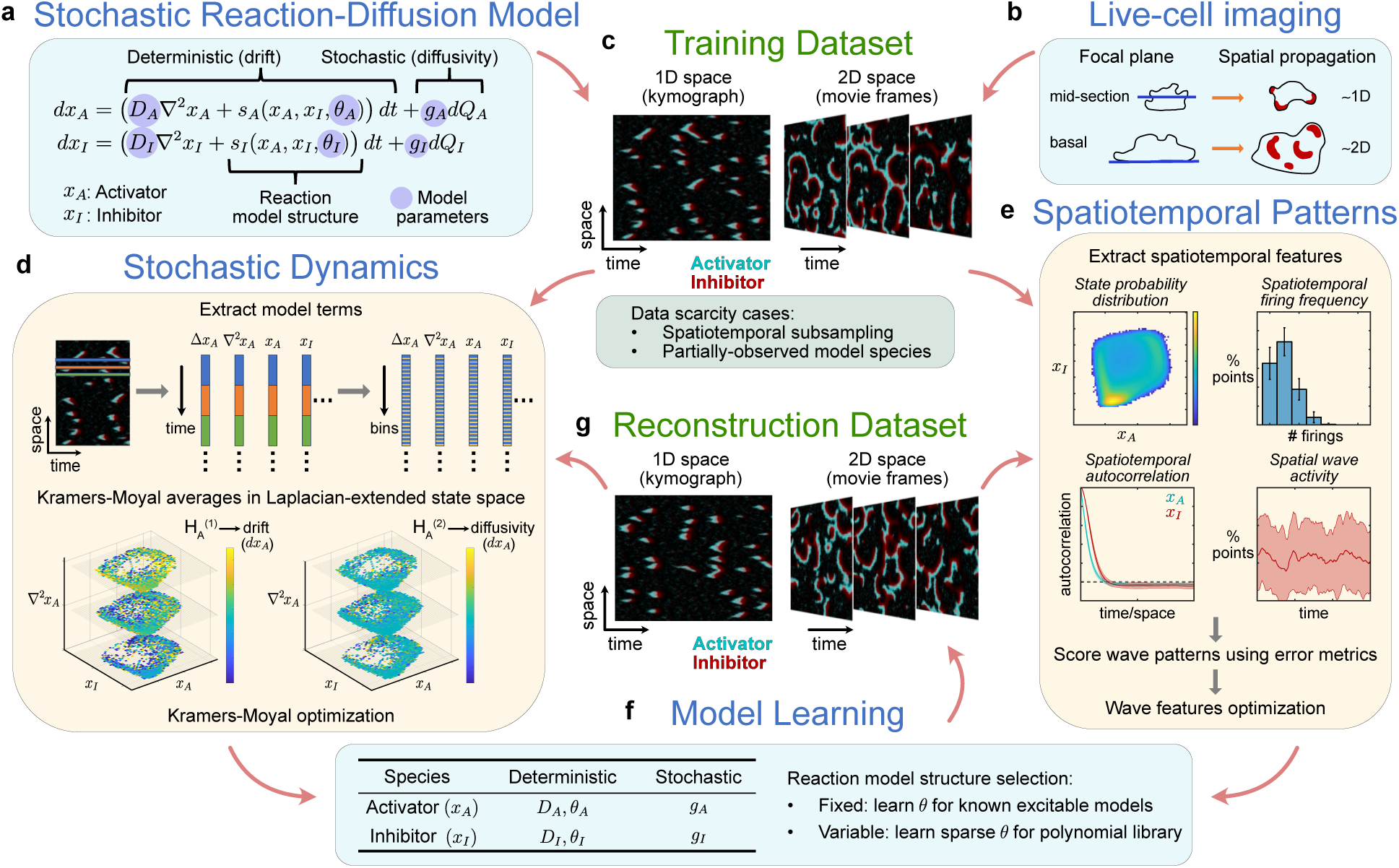
General approach for learning stochastic reaction-diffusion models from data. **a.** A stochastic reaction-diffusion system with two species: an activator (*x_A_*) and an inhibitor (*x_l_*). The system is modeled using coupled stochastic PDEs with drift (deterministic) and diffusivity (stochastic) components. A set of constant reaction parameter vectors (*0 _A_*, *0_l_*), spatial diffusion parameters (*D_A_*, *D_l_*), and noise parameters (*g_A_*, *g_l_*) specify the model. **b**. Live-cell microscopy data provides time-series of 1D or 2D spatial signals, depending on the focal plane during image acquisition. **c.** Examples of spatiotemporal data based on simulations of the equations in **a** or acquired from microscopy experiments in **b**. Multiple runs are performed to generate a training dataset for simulations. Data limitations are selected to mimic cases observed in live-cell imaging data. **d.** Data-driven learning using Kramers-Moyal (KM) averages estimating stochastic dynamics for spatially-distributed systems. Regression is used to fit model functions based on the drift and diffusivity to KM averages computed for each simulation in the dataset. **e.** Data-driven learning using spatiotemporal patterns. Sample features shown are extracted from datasets and scored to assess the similarity of the training vs. reconstruction datasets. The metrics are also used to estimate parameters that optimize wave features. **f.** Learning model structure and parameters in cases of fixed or variable model structure. **g.** Example spatiotemporal simulations obtained using optimal parameters from **f**. Multiple runs are performed to generate a reconstruction dataset.

Note that the deterministic components include state-dependent reaction terms (*s_A_*, *s_l_*), with a set of constant reaction parameters (*0 _A_*, *0_l_*), as well as spatial Laplacian terms (∇^2^*x_A_*, ∇^2^*x_l_*), with constant spatial diffusion parameters (*D_A_*, *D_l_*). The stochastic component is modeled in the activator dynamics by *dQ_A_*, an increment of a zero mean Gaussian white noise process with variance ∝ *dt*, and *g_A_* as proportionality constant. For all training data, no state noise was modeled in the inhibitor equation (*g_l_* = 0). The full set of model parameters are given by 0*_A_* = {*D_A_, 0 _A_, g_A_*} and 0*_l_* = {*D_l_, 0_l_, g_l_* }.

For clarity, we denote the activator and inhibitor species as *u* and *v* in the FHN model and as *f* and *r* in the FR model. As in previous studies (*22, 68*), the model structures for the FHN and FR models are distinguished by the reaction terms such that,

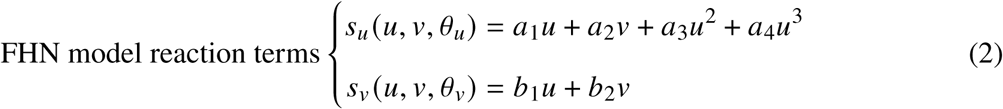

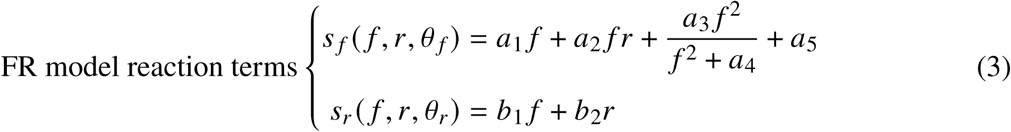

where reaction parameters *0 _A_*= {*a_i_*} and *0_l_* = {*b_i_*}. Note that while both models are nonlinear in the state variables, the FHN model is linear in the parameters, while the FR model is nonlinear in the parameters because of the more biochemically relevant Michaelis-Menten-like term. Additionally, while the FHN model variables are based on model membrane potential and take on negative values, the FR model variables describe the concentrations of biochemical species and generally remain non-negative during simulation.

### Visualization of spatiotemporal data from excitable systems

Figure 2 presents visualizations of the data observed in 1D and 2D simulations of the FHN and FR models. Three different visualizations are shown: the spatiotemporal trajectories, the temporal dynamics at a single spatial grid point, and phase portraits of system trajectories at a single grid point. The spatiotemporal trajectories of 1D simulations were visually represented fully in kymographs, where the x-axis is time, and the y-axis represents spatial points, while for 2D simulations, selected frame sequences of a movie were shown.

**Figure 2:**
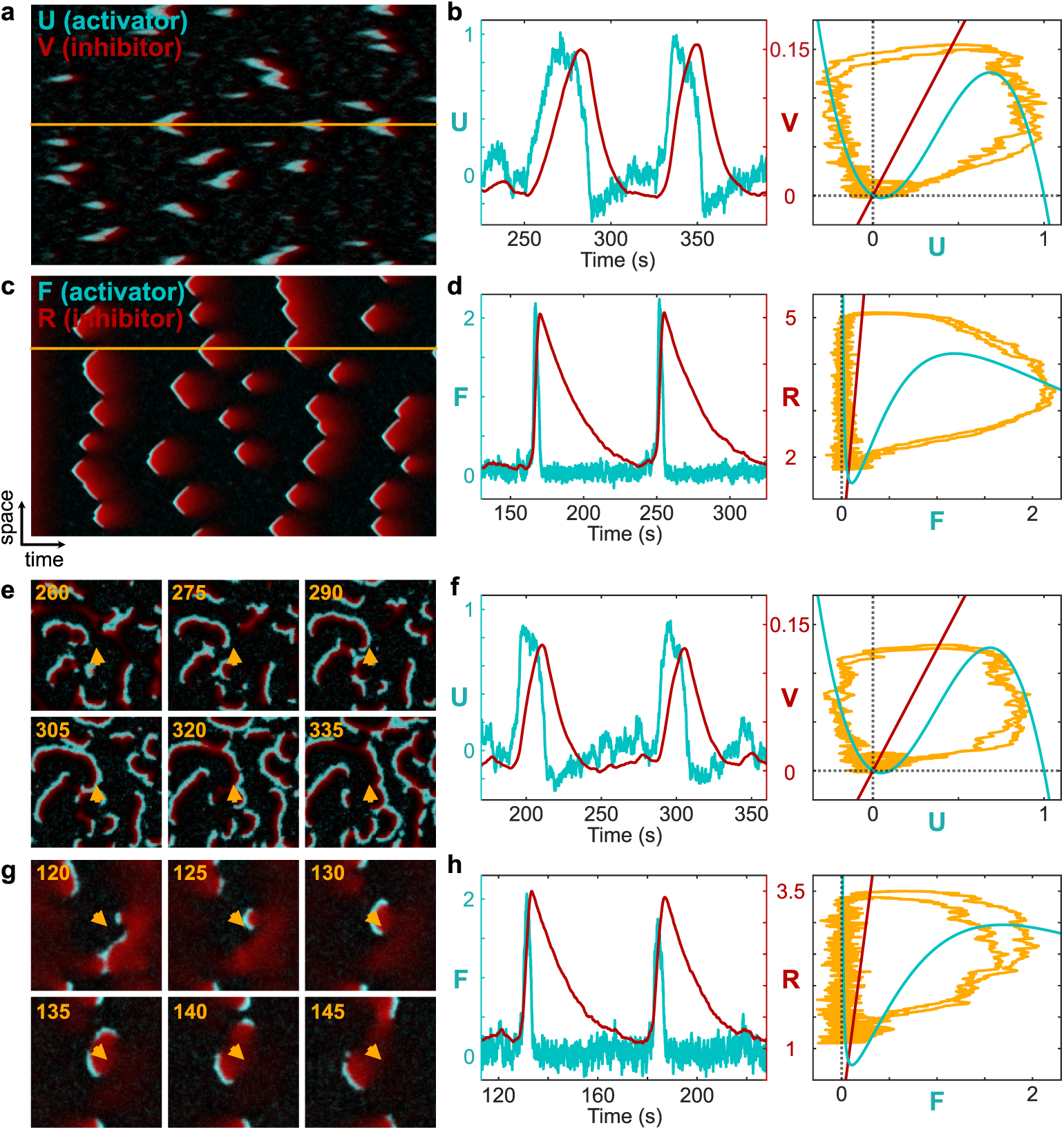
Visualization of training simulations of excitable reaction-diffusion models simulated in 1D and 2D space. **a.** Kymograph of a 400 s stochastic reaction-diffusion simulation based on the classical FitzHughNagumo (FHN) model over a 1D spatial domain (FHN-1D). Activator (U) and inhibitor (V) signals are shown in cyan and red, respectively. **b.** Signal values of U and V plotted together over time (left), and trajectories of U and V plotted in the phase plane (right). Both correspond to signals at the grid point indicated in **a** by the yellow line. **c-d.** Similar visualizations of the FR-1D model. F and R denote the activator and inhibitor, respectively. **e-h.** Frames from FHN (**e**) and FR (**g**) models simulated over a 2D spatial domain for 400 s. Time stamps indicate simulation time (seconds). Yellow arrows indicate spatial points at which trajectories were plotted over time (**f,h**, left) and in the phase plane (**f,h**, right).

Dynamic features spontaneously emerged in both the FHN-1D and FR-1D kymographs (Fig. 2a,c) with the formation of 1D waves originating at random initiation points and expanding in V-shaped patches in space-time with characteristic length scales in both time and space. The activator and the inhibitor in these two models demonstrated similar spatial profiles as visible by the significant overlap in the vertical propagation distance of both the activator and the inhibitor for any given wave (Fig. 2a,c). Temporally, FHN-1D wave patterns for the activator and the inhibitor showed behavior characteristic of excitability and exhibited similar duration as indicated by the strong horizontal overlap in the patches. The temporal signal at a single grid point in the left plot in Figure 2b shows this more clearly, with activator and inhibitor spikes lasting about 40 and 50 seconds, respectively, and separated by a delay of ∼20 seconds. In contrast, in the FR-1D model, the wave patterns for the activator lasted significantly shorter (∼20 s) compared to the inhibitor (∼60 s), and with shorter delay (∼5 s) between the two species (Fig. 2d, left).

Examining trajectories in phase space, the FHN-1D and FR-1D models demonstrated similar excursions during a firing when the system crossed an activation threshold as a result of noise or spatial diffusion (Fig. 2b,d). The nullclines of the dynamics of the reaction terms (without spatial diffusion) are displayed together with the trajectories at a single grid point as a wave passed through. The behaviors in the phase plane are indicative of a system with a stable equilibrium point near a Hopf bifurcation point. As two waves passed through the single grid point shown in orange in Figure 2a and were plotted in time (Fig. 2b, left), the FHN-1D activator and inhibitor traced a stereotypical trajectory in the phase plane (Fig. 2b, right). The excursion began from the equilibrium point (intersection of the nullclines) and moved to the right as the activator increased while the inhibitor remained low, then moved upward as the delayed inhibitor accumulated, then moved left as the activator decreased by the negative feedback from the inhibitor, before returning to the equilibrium. In the FR-1D simulations, for two waves passing through a grid point shown in Figure 2c, we displayed the time series and corresponding trajectories in the phase plane (Fig. 2d). Compared to the FHN model, the FR model parameters yielded faster activator dynamics and slower inhibitor decay.

Excitability in 2D spatial simulations formed traveling waves with rich features. FHN-2D showed circular waves that spontaneously initiated and propagated radially (Fig. 2e,Movie S1). Over time, as a result of annihilations when inhibitor signals from two waves merged, large waves were split into patches propagating through the spatial domain (Fig. 2e). FR-2D simulations showed similar traveling waves but with limited propagation range (Fig. 2g, Movie S2). The inhibitor R appeared to accumulate rapidly and with long duration, hindering propagation and extinguishing the waves (Fig. 2g). The temporal signals at a single grid point in Figure 2f,h show this distinction more clearly between the FHN-2D and FR-2D models. In FHN-2D, the activator and inhibitor lasted ∼30–50 seconds and were separated by a slight delay of ∼10 seconds (Fig. 2f, left). In contrast, in the FR-2D model, the wave patterns for the activator were significantly shorter (∼5–10 s) compared to the inhibitor (∼40 s), with a time delay under 5 seconds (Fig. 2h, left). The trajectories in the phase plane at the same grid point plotted in Figure 2e,g for both FHN-2D and FR-2D, were remarkably similar to those found in 1D spatial simulations (Fig. 2f,h, right). The FR-2D simulation showed some slight differences compared to FR-1D in phase plane analysis with the trajectories at a single grid point not settling to the equilibrium point of the spatially-homogeneous reaction equation nullclines (Fig. 2h, right).

### Estimating FHN reaction-diffusion model parameters from stochastic dynamics

Kramers-Moyal regression, a parameter estimation algorithm based on stochastic dynamics, was initially applied to FHN-1D and FHN-2D simulations, assuming a known reaction-diffusion model structure for each. A training dataset of 50 1D and 2D simulations recorded every *τ*_0_ = 0.1 s, was used for analysis (Methods). From the kymographs or movies, Kramers-Moyal (KM) averages were computed over a binned Laplacian-extended state space (Supplementary Methods). The first KM average, 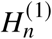, relates to the deterministic component of the FHN stochastic reaction-diffusion equation (SRDE), as a function of the binned values of *u*, *v*, and the Laplacian of the corresponding *n*-th species. For visualization, projected KM averages were displayed as a heatmap over the activator-inhibitor phase plane. As seen in Fig. S1, the heatmap for 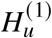 and 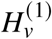 matched the stochastic dynamics observed in the time series at single grid points (Fig. 2b). The 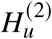 heatmap corresponding to the constant stochastic term in the activator SRDE was relatively constant, while the 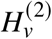 heatmap for the inhibitor was approximately zero, owing to the absence of stochastic noise in the inhibitor dynamics.

Linear regression over a library of basis functions, corresponding precisely to the terms in the FHN model, was performed on these KM averages to identify an optimal set of parameters that minimized the mean squared error (MSE) between empirical KM averages and the corresponding deterministic/stochastic model terms (theoretical KM averages). The parameters, computed separately for each simulation, demonstrated remarkable accuracy and precision when compared to the ground truth values used in simulation generation, even across a wide range of magnitudes (Table 1). To validate these parameters further, a set of 50 FHN-1D and FHN-2D simulations were generated based on the estimated parameters, forming a reconstruction dataset (Fig. 3). Comparisons between representative kymographs from the training and reconstruction datasets (Fig. 3a,e) revealed the inherent stochastic nature of the system, with no two simulations being identical. Feature analysis was employed for a more nuanced comparison of the two datasets. Characterization of these spatiotemporal features involved computing metrics for temporal firing frequency, spatial wave activity, and spatiotemporal autocorrelations (Fig. 3b–d,f–h).

**Figure 3:**
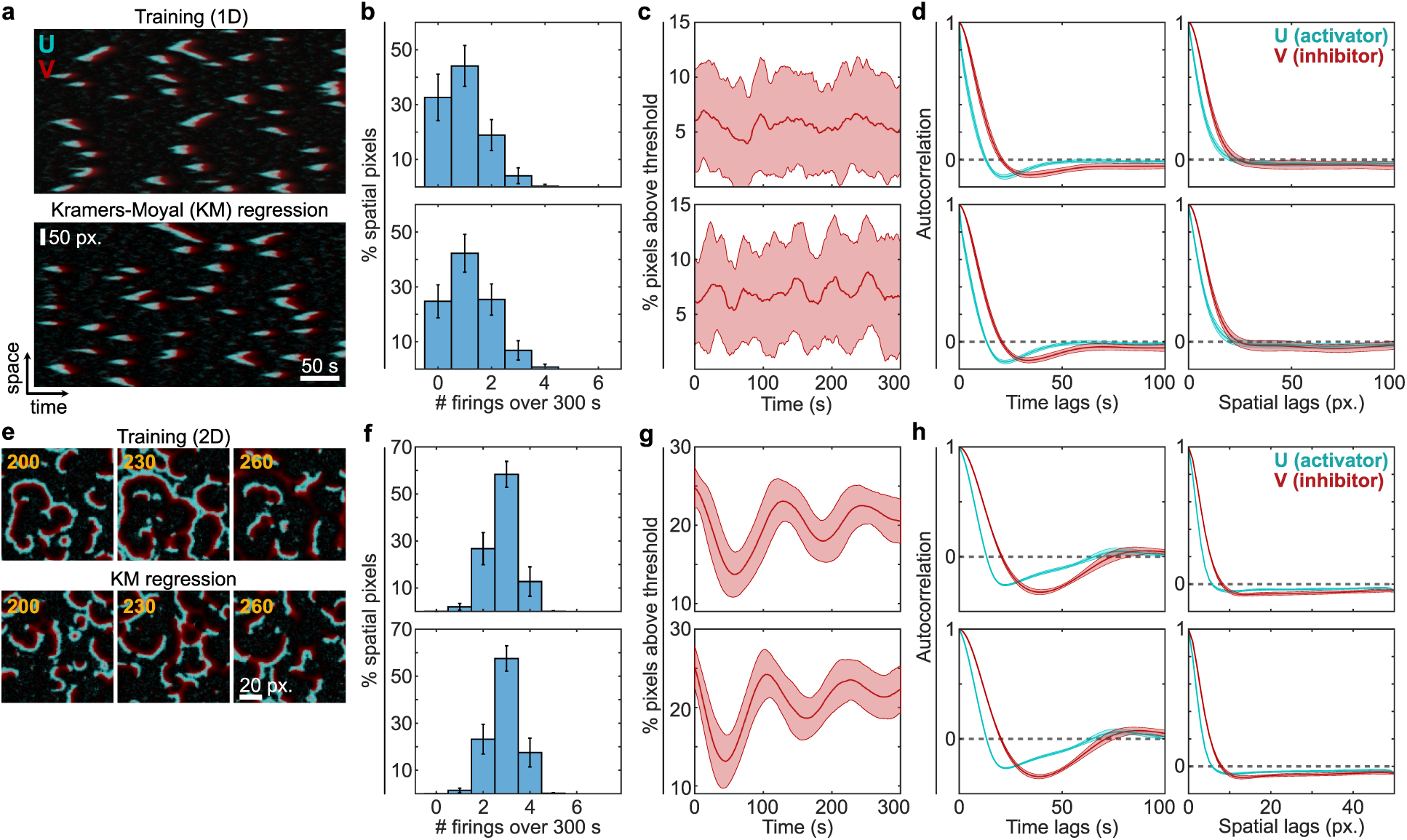
Reconstruction and validation with Kramers-Moyal (KM) based estimates of FHN reaction-diffusion model parameters. **a.** Representative kymographs from a 400 s simulation in the FHN-1D training dataset (top) and from a reconstruction dataset generated by FHN reaction-diffusion model parameters estimated using KM regression (bottom). (N=50 each; 50 spatial pixels (px.) and 50 s fiducial markers apply to both kymographs). **b.** Temporal firing frequency, computed as the distribution of the number of firings in a 300 s interval at spatial points over all simulations in the FHN-1D training data (top) and KM-based reconstructions (bottom). (N=50 each; error bars indicate SD). **c.** Spatial wave activity, computed as the percentage of spatial points (pixels) above threshold in a 300 s interval over all simulations in the FHN-1D training data (top) and KM-based reconstructions (bottom). (N=50 each; mean ± SD is plotted). **d.** Spatiotemporal patterns as quantified on each training kymograph (top row) and KM-based reconstruction (bottom row) using a temporal autocorrelation function averaged over spatial grid points (left column) and a spatial autocorrelation function averaged over time points (right column). (N=50 each; mean ± SD is plotted). **e.** Frames from representative training and KM-based reconstruction simulations for the FHN-2D reaction-diffusion model. (N=50 each). **f–h.** Similar to **b–d**, for FHN-2D reaction-diffusion simulations.

**Table 1:** True and estimated stochastic reaction-diffusion equation (SRDE) parameters for FHN-1D and FHN-2D models. Estimates using Kramers-Moyal regression on training simulations (N=50).

| | $D_u$ | $a_1$ | $a_2$ | $a_3$ | $a_4$ | $g_u$ |
| --- | --- | --- | --- | --- | --- | --- |
| **FHN-1D** |  |  |  |  |  |  |
| Ground truth | 1.4 | -0.2 | -2 | 2.2 | -2 | 0.1 |
| Mean | 1.382 | -0.193 | -1.984 | 2.163 | -1.969 | 0.103 |
| SD | 0.035 | 0.025 | 0.070 | 0.089 | 0.091 | 0.001 |
| **FHN-2D** |  |  |  |  |  |  |
| Ground truth | 0.13 | -0.2 | -2 | 2.2 | -2 | 0.0725 |
| Mean | 0.129 | -0.189 | -1.986 | 2.140 | -1.942 | 0.0735 |
| SD | 0.003 | 0.014 | 0.031 | 0.048 | 0.048 | 0.001 |
| | $D_v$ | $b_1$ | $b_2$ | $g_v$ | | |
| **FHN-1D** |  |  |  |  |  |  |
| Ground truth | 1.2 | 0.02 | -0.06 | 0 |  |  |
| Mean | 1.194 | 0.0200 | -0.0599 | 0.0006 |  |  |
| SD | 0.005 | $2.2 \times 10^{-5}$ | $1.3 \times 10^{-4}$ | $2.5 \times 10^{-5}$ | | |
| **FHN-2D** |  |  |  |  |  |  |
| Ground truth | 0.00625 | 0.0150 | -0.0450 | 0 |  |  |
| Mean | 0.00626 | 0.0150 | -0.0449 | 0.0006 |  |  |
| SD | $1.00 \times 10^{-4}$ | $9.16 \times 10^{-6}$ | $6.08 \times 10^{-5}$ | $9.20 \times 10^{-6}$ | | |

For both 1D and 2D simulations, the FHN training and reconstruction data displayed similar distributions for the number of firing events at a grid point, with the peak centered around 1 or 3 firings for 1D or 2D simulations, respectively, and a slight rightward skew for 1D data due to a small proportion of grid points with greater than 2 firings (Fig. 3b,f). The spatial wave activity, as indicated by the proportion of spatial points above the threshold at any given time, was remarkably consistent, with variability centered around 5% for 1D models and 20% for 2D models (Fig. 3c,g). Wave activity was particularly consistent in the 2D model in which oscillations continued through the last 300 s of training and reconstruction simulations. Temporal and spatial autocorrelations were also strikingly similar, indicating comparable pattern length scales in time (1D: ∼40-50 s; 2D: ∼50-70 s) and space (1D/2D: ∼5-10 px.) for both the training and reconstruction FHN datasets (Fig. 3d,h).

### Estimating FR reaction-diffusion model parameters from stochastic dynamics

The process of identifying model parameters for the FR model, under the assumption of a known reaction-diffusion model structure, involved the application of the KM regression algorithm with modifications. Given the faster dynamics, the FR training dataset was analyzed using simulations recorded at *τ*_0_ = 0.05 s. KM averages were computed from the kymographs or movies, and non-linear regression (using the Levenberg-Marquardt algorithm) was subsequently performed to determine the optimal set of parameters minimizing the MSE between the empirical KM averages and the corresponding deterministic/stochastic terms in the FR stochastic reaction-diffusion equation (Methods). The comparison of estimated parameters to the ground truth values of training data revealed their remarkable accuracy and precision across both 1D and 2D spatial FR simulations (Table 2). The estimated parameters for the activator in the FR model displayed larger variability compared to those for the inhibitor, which lacked a stochastic term in its equation. Additionally, the variability of parameters estimated for FR was slightly less than that observed for FHN, likely due to the smaller recording time step used for FR data analysis.

**Table 2:** True and estimated stochastic reaction-diffusion equation (SRDE) parameters for FR-1D and FR-2D models. Estimates using Kramers-Moyal regression on training simulations (N=50).

| | $D_f$ | $a_1$ | $a_2$ | $a_3$ | $a_4$ | $a_5$ | $g_f$ |
| --- | --- | --- | --- | --- | --- | --- | --- |
| **FR-1D** |  |  |  |  |  |  |  |
| Ground truth | 2.6 | −0.00835 | −0.8335 | 8.35 | 1.44 | 0.0635 | 0.15 |
| Mean | 2.583 | 0.00643 | −0.8305 | 8.280 | 1.439 | 0.0620 | 0.161 |
| SD | 0.025 | 0.0484 | 0.0078 | 0.126 | 0.016 | 0.0038 | 0.001 |
| **FR-2D** |  |  |  |  |  |  |  |
| Ground truth | 1.14 | −0.0120 | −1.200 | 12.024 | 2.88 | 0.0482 | 0.31 |
| Mean | 1.135 | 0.0177 | −1.199 | 11.898 | 2.873 | 0.0435 | 0.324 |
| SD | 0.013 | 0.0546 | 0.014 | 0.157 | 0.058 | 0.0054 | 0.002 |
| | $D_r$ | $b_1$ | $b_2$ | $g_r$ | | | |
| **FR-1D** |  |  |  |  |  |  |  |
| Ground truth | 5.2 | 0.5875 | −0.025 | 0 |  |  |  |
| Mean | 5.197 | 0.5872 | −0.0250 | 0.036 |  |  |  |
| SD | 0.004 | 0.0002 | $3.80 \times 10^{-5}$ | 0.0007 | | | |
| **FR-2D** |  |  |  |  |  |  |  |
| Ground truth | 1 | 0.5229 | −0.045 | 0 |  |  |  |
| Mean | 0.999 | 0.5226 | −0.0449 | 0.0384 |  |  |  |
| SD | 0.001 | 0.0003 | 0.0001 | 0.0007 |  |  |  |

To assess the robustness of these parameters, an additional 50 1D and 2D FR simulations were generated based on the estimated parameters, forming reconstruction datasets. A comparative analysis of representative kymographs and movies from the 1D or 2D reconstruction dataset (Fig. 4b,d) and the corresponding training dataset (Fig. 4a,e) underscored the inherent stochastic nature of the system, highlighting the unique nature of each simulation. As above, wave feature analysis was subsequently employed for a nuanced comparison of the two datasets.

**Figure 4:**
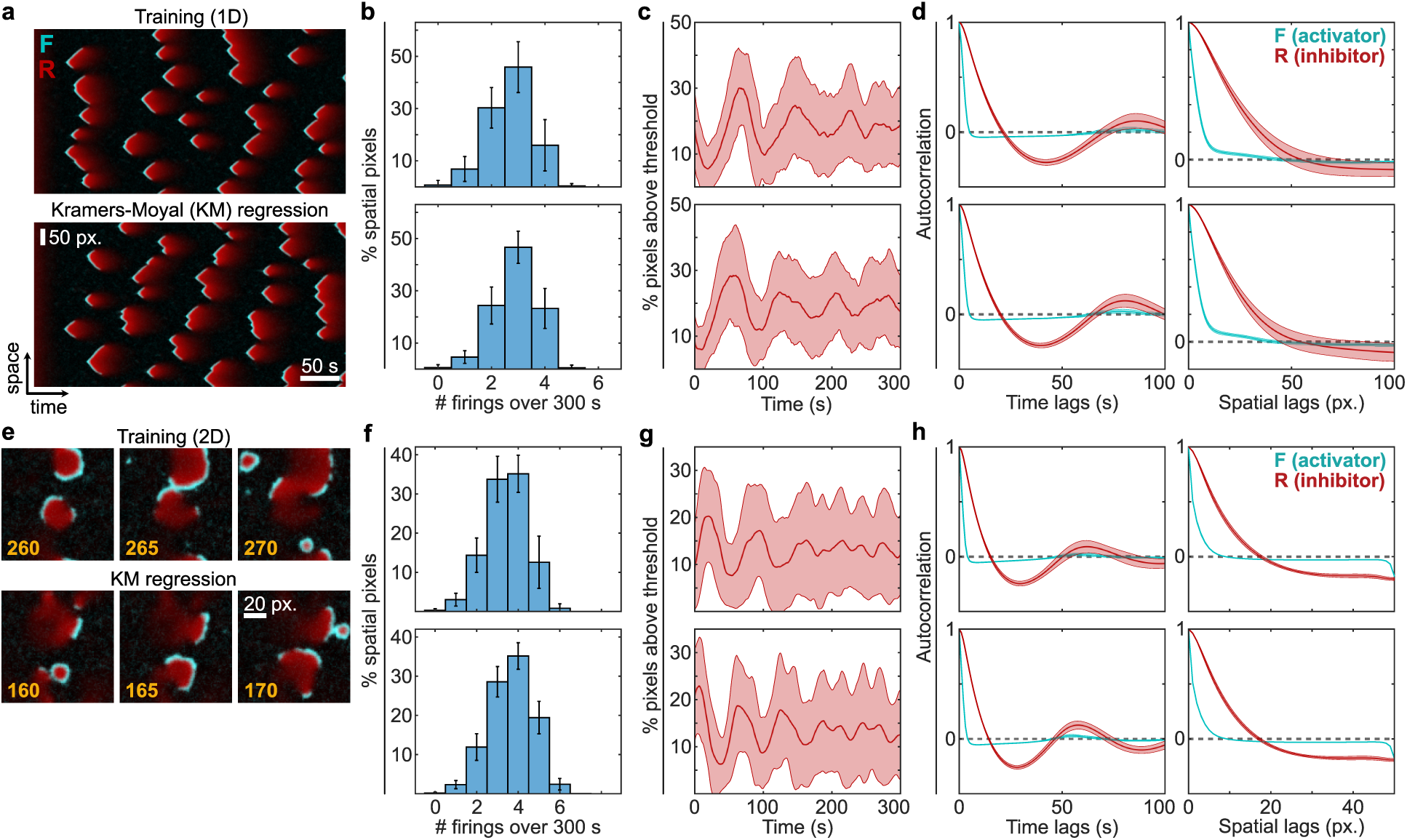
Reconstruction and validation with Kramers-Moyal (KM) based estimates of FR reaction-diffusion model parameters. **a.**Representative kymographs from a 400 s simulation in the FR-1D training dataset (top) and from a reconstruction dataset generated by FR reaction-diffusion model parameters estimated using KM regression (bottom). (N=50 each; 50 spatial pixels (px.) and 50 s fiducial markers apply to both kymographs). **b.** Temporal firing frequency, computed as the distribution of the number of firings in a 300 s interval at spatial points over all simulations in the FR-1D training data (top) and KM-based reconstructions (bottom). (N=50 each; error bars indicate SD). **c.** Spatial wave activity, computed as the percentage of spatial points (pixels) above threshold in a 300 s interval over all simulations in the FR-1D training data (top) and KM-based reconstructions (bottom). (N=50 each; mean ± SD is plotted). **d.** Spatiotemporal patterns as quantified on each training kymograph (top row) and KM-based reconstruction (bottom row) using a temporal autocorrelation function averaged over spatial grid points (left column) and a spatial autocorrelation function averaged over time points (right column). (N=50 each; mean ± SD is plotted). **e.** Frames from representative training and KM-based reconstruction simulations for the FR-2D reaction-diffusion model. (N=50 each). **f–h.** Similar to **b–d**, for FR-2D reaction-diffusion simulations.

A high degree of similarity was observed between training and reconstruction datasets. The number of firings at any grid point exhibited similar distributions in both FR training and reconstruction simulations (Fig. 4b,f), with a peak centered around 3 firings for 1D simulations, and around 4 firings for 2D simulations. The spatial wave activity, quantified by the proportion of grid points above threshold at any given time, exhibited high similarity between training and reconstruction simulations, with variability centered around 10% for 1D simulations and 20% for 2D simulations (Fig. 4c,g). Strikingly, even the slight oscillations in spatial wave activity over the last 300 s of training for 1D and 2D simulations were replicated in the reconstructions. The temporal and spatial autocorrelations also demonstrated a high degree of similarity between training and reconstruction data (Fig. 4d,h), indicating comparable wavelength scales in time (1D: ∼5–20 s; 2D: ∼5-15 s) and space (1D: ∼25-50 px.; 2D: ∼10-50 px.).

### Learning sparse polynomial reaction-diffusion models from FHN dynamics and patterns

Building on the above work estimating parameters for models with a fixed structure for the deterministic and stochastic components of the stochastic reaction-diffusion equation (SRDE), we next addressed the challenge of learning model structure and parameters together from FHN-1D training simulations. Specifically, we considered a library of polynomial reaction-diffusion models based on equations 1, 2, and E23. Note that the library varies only in the polynomial reaction terms in the deterministic components of the SRDE, and that the forms of the spatial diffusion and noise terms were held fixed for both the activator and inhibitor equations. Importantly, the true FHN reaction-diffusion model structure is a subset of this library.

To learn sparse models initially from stochastic dynamics, we used sparse KM regression (Methods). This involved a series of linear regressions to minimize the MSE between the empirical and parameterized theoretical KM averages for the deterministic components of the activator and inhibitor SRDE separately. Starting with 11 deterministic parameters (one spatial diffusion and ten polynomial reaction model terms) for the initial regression, we sequentially eliminated the smallest reaction parameter in absolute value at each step. The stochastic components of the activator and inhibitor SRDE were estimated assuming constant noise coefficients. Figure 5a illustrates the specific model terms eliminated at each iteration. Sparse KM regression was applied separately to each of the 50 training simulations, and the proportion of simulations in which a specific model term in the polynomial reaction-diffusion library remained selected was plotted for each column (iteration). Owing to the inherent stochasticity in the training dataset, some variability in the eliminated terms was observed resulting in shaded regions, especially in early steps where few model terms had been eliminated (higher model complexity).

**Figure 5:**
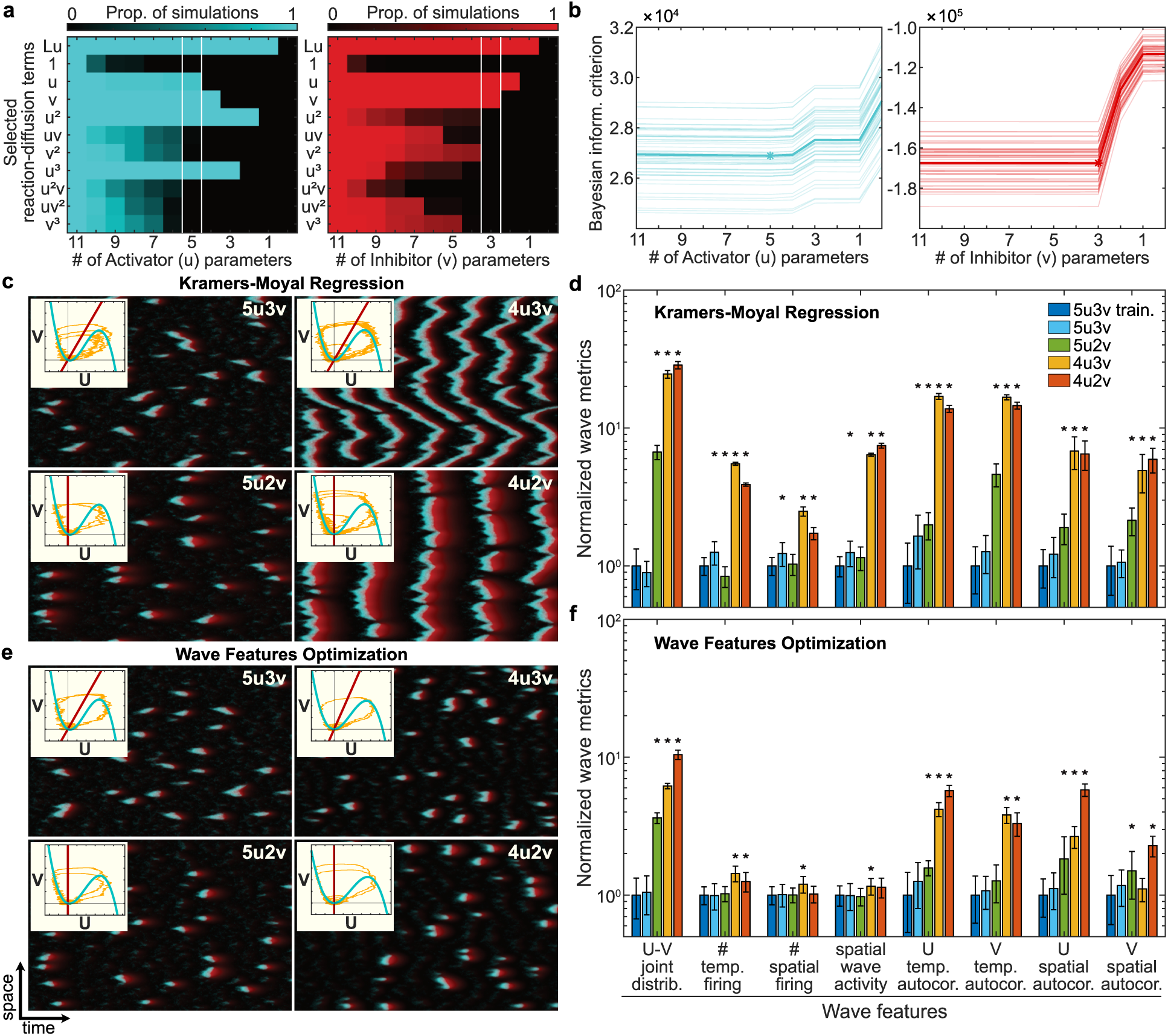
Learning sparse models from FHN-1D data by optimizing Kramers-Moyal (KM) averages and wave features. **a.**Reverse-greedy search approach estimating parameters using KM-based regression at each iteration, and sequentially eliminating terms from a library of polynomial reaction-diffusion models. This sparse KM regression was performed separately for the activator equation (left) and inhibitor equation (right), and repeated for each kymograph in the FHN-1D training dataset (N=50). Colors indicate the proportion of simulations for which each model term remained selected as model complexity decreased from 11 to zero parameters over the range of iterations. White lines indicate the optimal sparsity as estimated from the next panel. **b.** Bayesian information criterion (BIC) estimated at each iteration of sparse KM regression for the activator and inhibitor equations and plotted as separate lines for each simulation (average is in bold). The star indicates the most common minimum of the BIC (optimal sparsity) over all simulations. **c.** Reconstructions using KM-based parameters for four sparse models labeled by the number of terms in the activator (U) and inhibitor (V) equations: 5u3v (optimal sparsity; five terms for U equation, three terms for V equation), 5u2v, 4u3v, and 4u2v. Insets show representative nullclines and phase-plane trajectories. **d.** Spatiotemporal wave metrics computed for FHN-1D training data (N=50) and reconstruction datasets of the four sparse model structures (N=20 each). Metrics were normalized relative to training data and plotted on a logarithmic scale. Stars indicate statistically significant differences relative to training data for each wave feature (Wilcoxon rank-sum test, *p <* 0.01). **e.** Similar to **c**, for reconstructions based on parameters estimated for the same sparse model structures and optimizing for wave features. **f.** Similar to **d**, a quantitative comparison of each normalized wave metric in the reconstruction dataset for each sparse model structure (N=20) to the normalized wave metrics of the original FHN-1D training data.

To determine the optimal number of parameters, we employed the Bayesian Information Criterion (BIC), which balances accuracy and sparsity (Methods, Fig. 5b). Analysis of the mode of the distribution of optimal sparsity (BIC minimum) from all simulations (Fig. S2a) indicated that the optimal number of reaction-diffusion parameters was five for the activator and three for the inhibitor (denoted “5u3v”). The reaction-diffusion terms corresponding to these parameters were identified from the model terms at the iteration highlighted in white in Figure 5a, by selecting the top five activator terms and the top three inhibitor terms remaining over 50 training simulations (Fig. 5a). Consequently, this method successfully recovered the original FHN reaction-diffusion model structure.

Figure 5c presents representative reconstructions using parameters from sparse KM regression for the optimal model structure, 5u3v, alongside reconstructions using three sparser models: 5u2v, 4u3v, and 4u2v (Table S1). The 5u2v model displayed similar wave patterns to those of the 5u3v model, albeit with reduced activity, while the 4u3v and 4u2v models predominantly exhibited oscillatory behavior, differing in period and synchronization across the spatial dimension. The insets reveal that these differences arise because the inhibitor nullclines intersect the activator nullcline at the local minimum, positioning the system at a Hopf bifurcation.

To quantify the differences in the wave patterns clearly observed among the models tested, we considered several normalized wave metrics (Fig. 5d). These included the following averaged spatiotemporal features: the joint probability distribution of U and V, spatiotemporal wave frequency, spatial wave activity, and spatiotemporal patterns analyzed using autocorrelation functions. Analysis of these metrics revealed that reconstructions of the first two models (5u3v and 5u2v) showed reasonably close alignment with the training data, while the last two models (4u3v and 4u2v) exhibited significant discrepancies.

Motivated by these findings, we explored whether optimizing based on these wave features could identify model parameters that more closely matched the training data. Using a Nelder-Mead simplex search, we optimized a custom objective function involving the sum of squared relative errors in wave features of a reconstruction dataset relative to that of training FHN-1D data (Methods, Fig. S2b). Figure 5e shows kymographs for the optimal parameters based on wave features optimization for the four model structures considered earlier. The 5u3v and 5u2v models displayed similar behavior, whereas the 4u3v simulation showed no oscillations, and the 4u2v model exhibited broken patches with some oscillatory behavior. Quantification of individual raw wave features (Fig. S3, Fig. S4), normalized wave metrics (Fig. 5d), and overall wave metrics error (Fig. 5f) all confirmed the improved characteristics of simulations after wave features optimization.

### Learning sparse polynomial reaction-diffusion models from FR dynamics and patterns

Extending this approach to real systems requires fitting data-derived quantities to a library of generalized models that may not include the exact system. To explore the feasibility of this, we used training data generated by the FR-1D model which includes a rational term in the activator dynamics. As before, we began with initially fitting KM quantities to a polynomial reaction-diffusion library, and then fine-tuned parameters using wave features optimization. Unlike the FHN dataset, however, the true FR reaction-diffusion model structure is a subset of the library for the inhibitor (R) dynamics, but not for the activator (F).

In Figure 6a, we started with 11 deterministic parameters for both the activator and inhibitor and sequentially eliminated terms using sparse KM regression, as previously. In contrast to FHN data, here there was decreased variability (fewer shaded regions) in the selected reaction diffusion terms for both the activator and the inhibitor even at early iterations with higher model complexity, likely due to the smaller recording time step *τ*_0_ = 0.05 s used for analysis. Once again, we computed the BIC over all iterations for each simulation (Fig. 6b); then we used the mode of the distribution of optimal sparsity (BIC minimum) from all simulations to obtain the final estimate of optimal sparsity (Fig. S5a). The optimal model structure corresponded to a model denoted “9f3r” with 9 parameters for the activator (one spatial diffusion and eight reaction) and three parameters for the inhibitor (one spatial diffusion and two reaction). The model terms at the corresponding iterations highlighted in Figure 6a matched the exact structure of the deterministic component of the inhibitor equation; however, for the activator, the estimated 9-term polynomial reaction-diffusion model was quite distinct from the training FR model. Additionally, despite 9 parameters being optimal, we observed minimal differences in BIC from models with 9 parameters to 5 parameters, as seen in Fig. 6b.

**Figure 6:**
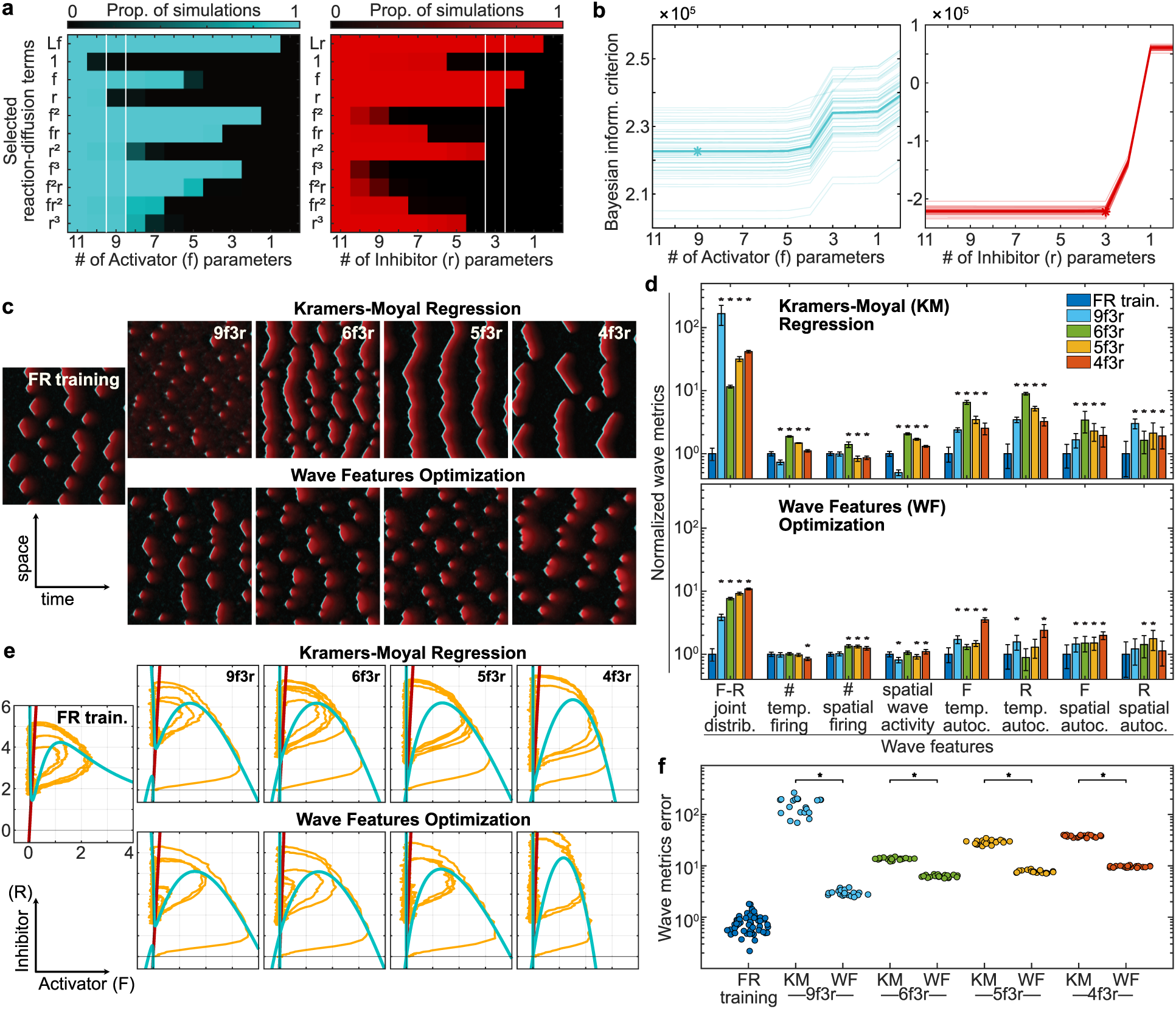
Learning sparse models from FR-1D data by optimizing Kramers-Moyal averages and wave features. **a.**Reverse-greedy search approach estimating parameters using KM regression at each iteration, and sequentially eliminating terms from a library of polynomial reaction-diffusion models. This sparse KM regression was performed separately for the activator equation (left) and inhibitor equation (right), and repeated for each kymograph in the FR-1D training dataset (N=50). Colors indicate the proportion of simulations for which each model term remained selected as model complexity decreased from 9 to zero parameters over the range of iterations. White lines indicate the optimal sparsity as estimated from the next panel. **b.** Bayesian information criterion (BIC) estimated at each iteration for the activator and inhibitor equations and plotted as separate lines for each simulation (average is in bold). The star indicates the most common minimum of the BIC (optimal sparsity) over all simulations. **c.** Reconstructions using KM-based parameters (top row) for four sparse models labeled by the number of terms in the activator (F) and inhibitor (R) equations: 9f3r (optimal sparsity; nine terms for U equation, three terms for V equation), 6f3r, 5f3r, and 4f3r. Bottom row shows representative reconstructions based on parameters estimated for the same sparse model structures and optimizing for wave features. A representative kymograph from the FR-1D training data is plotted on the far left for comparison. **d.** Spatiotemporal wave metrics computed for FR-1D training data (N=50) and both KM-based and wave features (WF) optimization-based reconstruction datasets of the four sparse model structures (N=20 each). Metrics were normalized relative to training data and plotted on a logarithmic scale. Stars indicate statistically significant differences relative to training data for each wave metric (Wilcoxon rank-sum test, *p <* 0.01). **e.** Representative nullclines and phase-plane trajectories for KM-based reconstructions (top row) and wave features based reconstructions (bottom row), as well as the original FR-1D training data (far left). **f.** Scalar wave metrics error computed for each simulation in the FR-1D training data, and both reconstruction datasets (KM-based and WF-based). Each dot represents the square root of the sum of the squared relative error between wave metrics in a given simulation and the mean wave metrics in the FR-1D training data. Stars indicate statistically significant differences (Wilcoxon rank-sum test, *p <* 0.01).

To examine the behavior of these different stochastic models, we generated a set of 20 reconstructions using the estimated parameters from sparse KM regression for the 9f3r model and three sparser models: 6f3r, 5f3r, and 4f3r (Table S2). Fig. 6c shows representative reconstructions with SRDE parameters estimated using KM regression. By visual inspection, the general wave patterns observed in simulations of three of the models deviated from the FR training data: 9f3r had considerably smaller wave firings compared to the training data; 6f3r, 5f3r, and 4f3r showed oscillatory behavior with increasing wave period. Spatiotemporal wave metrics computed from the reconstruction sets quantified the differences in the patterns observed for each model structure (Fig. 6d).

We then performed wave features optimization to identify model parameters to match the patterns observed in FR training data. The coarse-grained approach of sparse KM regression provided candidate sparse model structures with a wide range of spatiotemporal patterns significantly differing from the training data. Fine-tuning as before, we used initial parameters from sparse KM regression, and optimized a wave metrics error objective function over multiple rounds for the reaction, spatial diffusion, and noise parameters (Fig. S5b). Compared to patterns in FR training data, wave features (WF) optimization achieved improved matching of wave metrics for all model structures, with the most improvements visible in the 9f3r model (Fig. 6c). Quantification confirmed these trends across both individual raw wave features (Fig. S6, Fig. S7), normalized wave metrics (Fig. 6d), and overall wave metrics error (Fig. 6f). The improvement in wave metrics for each model tested demonstrated the success of this combined approach in learning sparse polynomial reaction-diffusion models even when the true FR model structure is not fully captured by a polynomial model.

Exploring possible reasons for the improvement, we plotted the spatially homogeneous reaction equation nullclines and phase plane trajectories for each of the models learned from KM regression and wave features optimization (Fig. 6e). In general, for both KM regression and wave features (WF) optimization, all the models had an approximate inverted “N” shape for the F-nullcline. However, for the sparser models, 6f3r, 5f3r, and 4f3r, the 4f3r model, this inverted shape consisted two curves for the F-nullcline: one vertical line at *F* = 0 and a second concave down curve. The intersecting curves resulted in an approximate inverted “N” shape in the region of the phase plane accessed by stochastic trajectories. During WF optimization, the local shape of the curves around the equilibrium point, which relates to system threshold, appeared to become tuned for matching excitability. For example, for the 9f3r model, compared to the reduced wave activity in the KM regression model, the WF optimization model exhibited a decrease in the F-nullcline minimum similar to the training model, resulting in a lower threshold and increased wave activity comparable to training data (Fig. 6e). Interestingly, for the sparsest 4f3r model, WF optimization resulted in more complex changes to the nullclines, but locally around the equilibrium point of the system, the curves moved closer to that of the training model, resulting in improved firing patterns (Fig. S5c).

### Estimating stochastic reaction-diffusion models from limited simulation data

We further challenged our approach by testing scenarios of low-resolution data arising in realworld contexts. Signals acquired from live-cell microscopy are often propagating in 2D space and are measured every few seconds, with only 1-2 species observed at a time. To evaluate estimating stochastic reaction-diffusion models from limited data, in Figure 7 we applied our approach to estimating parameters for the FHN-2D model from datasets with various rates of temporal sampling and observations limited to only one of the species (activator (U) or inhibitor (V)).

**Figure 7:**
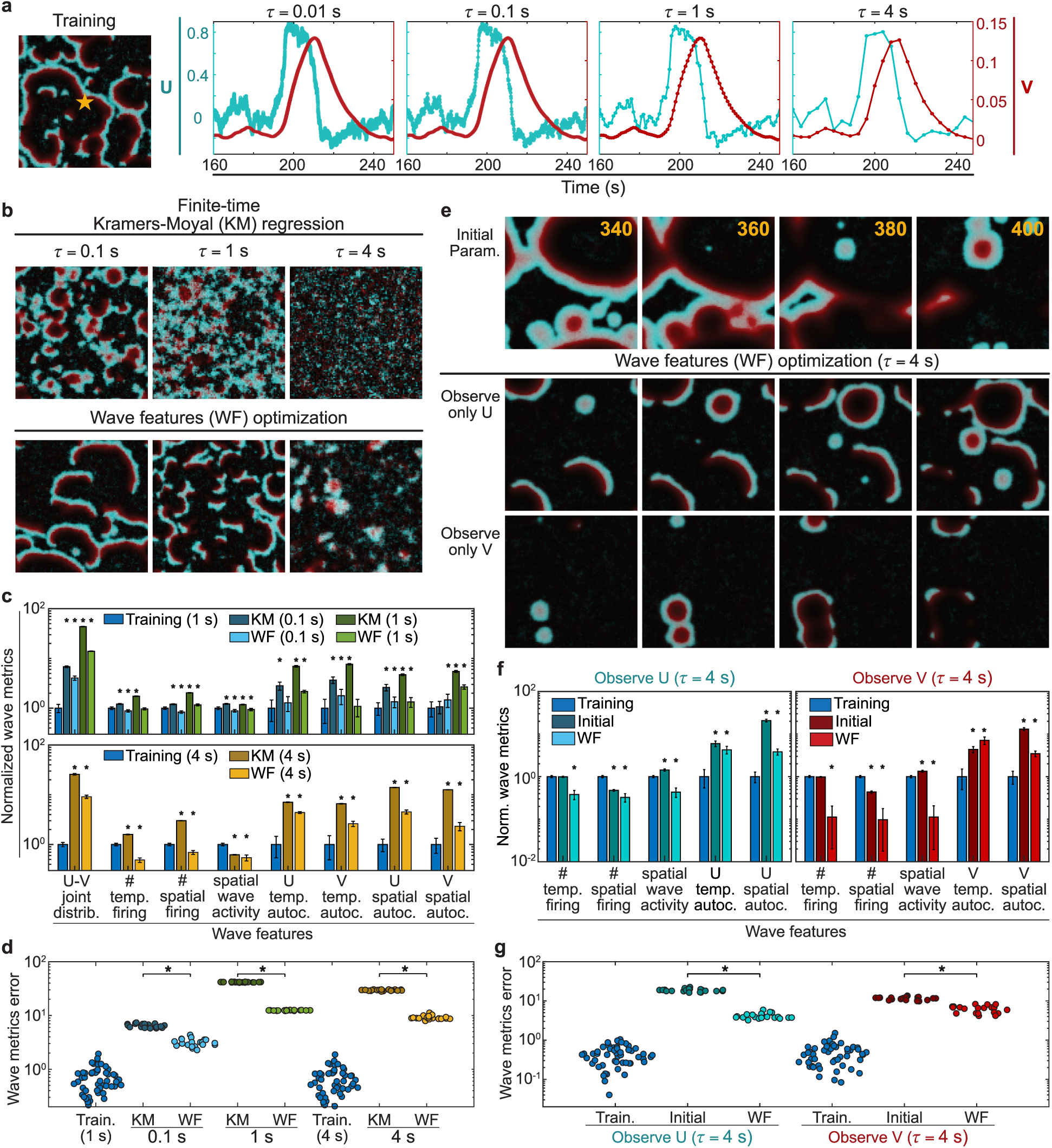
Estimating 2D-spatial stochastic reaction-diffusion models from limited simulation data. **a.** Maximum intensity frame (226 s) from a 2-D spatial simulation in the FHN-2D training dataset. Time series for the activator (U, cyan) and inhibitor (V, red) were extracted at a single spatial grid point (orange star), and plotted with decreasing temporal sampling rates, from *r* = 0.01 s (integration time step) to *r* = 4 s. **b.** Maximum intensity frames from reconstructions based on FHN-2D model parameters estimated using finite-time Kramers-Moyal (KM) regression (top row) or wave features (WF) optimization (bottom row). Finite-time KM regression parameters were estimated from data sampled at *r* = 0.01, 0.1, or 4 s and then used as initial parameters for wave features optimization. **c.** Wave metrics corresponding to 8 wave features computed for reconstruction datasets (N=20 simulations) generated with parameters from finite-time KM regression or WF optimization and compared to that of training data (N=50 simulations). For parameters estimated from data sampled at *r* = 0.01 and 1 s, wave metrics from corresponding reconstructions were computed at *r* = 1 s and compared with metrics from training data sampled at *r* = 1 s (top panel). For parameters estimated from data sampled at *r* = 4 s, wave metrics from the corresponding reconstructions were computed at *r* = 4 s and compared with metrics from training data sampled at *r* = 4 s (bottom panel). **d.** Wave metrics errror based on the wave features objective function used during optimization and computed for FHN-2D training data sampled at 1 s and 4 s, as well as reconstruction datasets from finite-time KM regression and WF optimization. Each dot represents the error between wave metrics in each simulation and the mean wave metrics of the training data. **e.** Frames from simulations generated with an initial parameter set (top row) and after wave features optimization when only observing either activator (middle panel) or the inhibitor (bottom panel) signals sampled at *r* = 4 s. The time stamps (seconds) in the top panel indicate corresponding simulation time points for the frames in each column. **f.** Normalized wave metrics corresponding to 5 singlespecies wave features computed for reconstruction datasets (N=20 simulations) generated with parameters from WF optimization and compared to that of training data (N=50 simulations) when only observing U (left panel), or V (right panel). **g.** As in panel **d.**, wave metrics error based on the single-species wave features objective function used during optimization and computed for FHN-2D training data as well as reconstruction datasets from WF optimization. Stars indicate statistically significant differences between compared groups, or differences relative to training data, as in **c** or **f**, respectively (Wilcoxon rank-sum test, *p <* 0.01).

In Fig. 7a, we show the effects of different rates of temporal subsampling on data quality obtained at a particular spatial grid point. Compared to full signals acquired at the simulation time step of *dt* = 0.01 s, decreasing sampling rates with acquisition every *τ* = 0.1, 1, 4 s resulted in smoothing of high frequency components in the stochastic signals. However, subsampling at even *τ* = 4 s, with a 99.75% reduction in temporal data, still retained much of the general profile of a propagating wave. To study the effects of subsampling on parameter estimation, we first computed finite-time KM averages (Fig. S8) for each of the four sampling rates, while adjusting the number of training simulations used for consistent dataset size. We then applied KM regression to estimate parameters for the deterministic and stochastic components of the SDE that best fit the finite-time KM averages and generated reconstruction datasets of 20 simulations for each case. Maximum intensity frames from representative 2D-spatial simulations based on parameters from finite-time KM regression showed the qualitative deterioration of patterns as sampling rate decreased, with barely any visible patterns at *τ* = 4 s (Fig. 7b, top panel; Movie S3, Movie S4).

Next, we checked whether wave features optimization could yield improved estimates from subsampled data. Similar to before, we first computed wave metrics from FHN-2D training data based on both the activator and inhibitor signals sampled every *τ* = 1 or *τ* = 4 s (Methods). Using the previous objective function with subsampled wave features, we then optimized over the set of reaction, diffusion, and noise parameters of a fixed FHN-2D model structure (Table S3). We tested three cases, each with an initial set of parameters obtained from finite-time KM regression (Fig. 7b, top panel): 1) *τ* = 0.1 s, 2) *τ* = 1 s, and 3) *τ* = 4 s. For cases 1 and 2, wave features were optimized to training data wave features at *τ* = 1 s; for case 3, wave features were optimized to training data wave features at *τ* = 4 s. A set of 20 reconstructions were generated for each case, and maximum intensity frames from representative simulations demonstrated marked improvement in wave patterns for cases 1 and 2, and moderate improvement for case 3 (Fig. 7b, bottom panel; Movie S3, Movie S4). Significant improvements in individual raw wave features (Fig. S9, Fig. S10), normalized wave metrics (Fig. 7c), and wave metrics error (Fig. 7d) were observed over the range of cases, demonstrating the effectiveness of wave features optimization for parameter estimation of 2D-spatial reaction-diffusion models from subsampled datasets with limited dataset size.

To further examine the robustness of our approach, we tested the feasibility of parameter estimation from FHN-2D training datasets that were subsampled and only partially observed. Here, we used subsampled data at *τ* = 4 s and considered two cases: 1) only observing the activator (U), and 2) only observing the inhibitor (V). Since KM averages cannot be estimated from partially-observed systems, we only focused on wave features optimization. For initialization, we used a set of parameters designed to give near-oscillatory behavior with large propagating waves that qualitatively differed from training data as visible in the frames from 2D-simulations (Fig. 7e, top row). Because only one species was observed, we computed a set of 5 wave features from FHN-2D training data for each case, and optimized the objective function based on this subset of wave features to estimate FHN-2D reaction-diffusion model parameters (Table S4). Representative frames from reconstructions based on parameters after wave features optimization demonstrated the improvements in wave patterns (Fig. 7e, Movie S5, Movie S6). Remarkably, in both cases, the wave propagation range and wave profile thicknesses of both species more closely matched that of training data (Fig. 7e). Optimization when only U signal was observed appeared to show more similar wave activity and frequency to that of training data, while optimization when only V signal was observed resulted in markedly reduced wave activity and frequency. Quantitative analysis of each of the 5 raw wave features (Fig. S11), normalized wave metrics (Fig. 7f), and the overall wave metrics objective function each confirmed these trends (Fig. 7g). These results indicate the effectiveness of wave features optimization to generate parameter estimates for 2D-spatial reaction-diffusion models from simulation datasets with limited observations.

### Estimating stochastic reaction-diffusion models from limited cell microscopy data

To further explore learning stochastic reaction-diffusion models from low-resolution data, we tested the algorithm on a cell microscopy dataset to identify optimal parameters for the FHN-2D model. For training, the dataset was limited in three ways: a) only a single species (PIP3) was observed, b) frames were acquired every 7 seconds (low temporal resolution), and c) only a single replicate consisting of 150 frames was used. Figure 8a demonstrates the pre-processing and extraction of spatiotemporal features for training. Because only one species was observed, we used projected finite-time first and second KM averages, two novel KM-based quantities, to estimate stochastic dynamics in this context. We also extracted two additional wave features as measures of spatiotemporal patterns: the probability distribution for a single species (PIP3) and the autocorrelation of the spatial wave activity over time as a measure of global system behavior (Fig. 8a; see Supplementary Methods for details).

**Figure 8:**
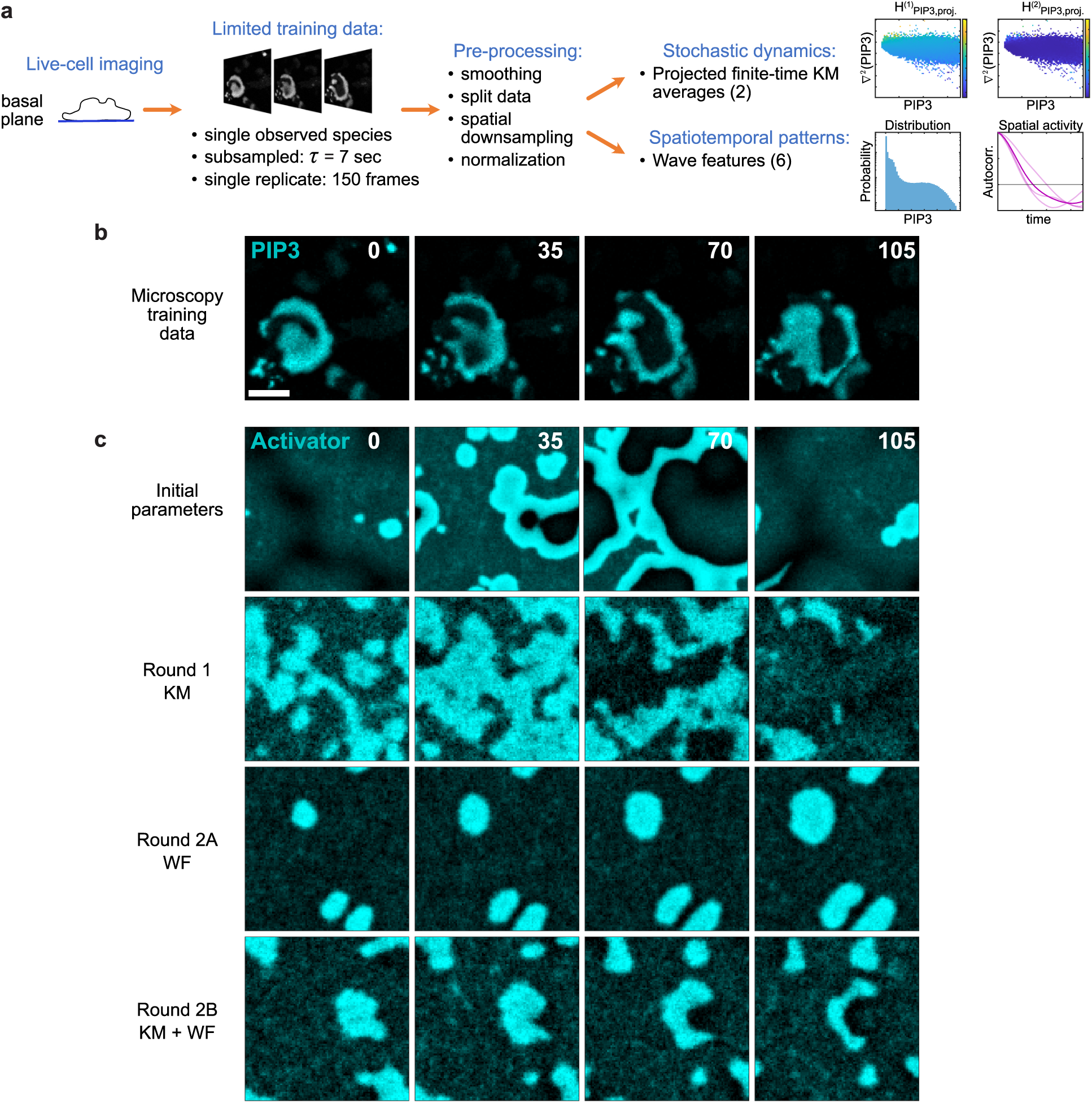
Estimating 2D-spatial stochastic reaction-diffusion models from limited cell microscopy data. **a.** Schematic of feature extraction from live-cell imaging data. Top row of sub-panels on the right show representative plots of projected finite-time, first and second Kramers-Moyal (KM) averages. Bottom row of sub-panels on the right show a representative plot of a single species’ probability distribution and autocorrelation of global spatial activity. **b.** Selected frames from training data of PIP3 signal shown in cyan on the substrate-attached surface of electrofused “giant” *Dictyostelium* cells. Images were acquired every 7 seconds for a total of 150 frames. Time stamps indicate imaging time in seconds. The scale bar indicates 20 *µ*m. **c.** Selected frames from reconstructions generated with different FHN-2D parameters: initial parameters before any optimization (row 1); final parameters after the first round of optimization using only projected finite-time KM averages (row 2); final parameters after a second round of optimization, starting with parameters from round 1, using only wave features (WF) (row 3); final parameters after a second round of optimization, starting with parameters from round 1, using both projected finite-time KM averages and wave features (KM + WF) (row 4). Activator signal is shown in cyan and was optimized to match PIP3 signal from **b**. Time stamps indicate simulation time in seconds relative to the first frame shown in each row.

In Figure 8b, we present sample frames from the training data. These frames exhibit dynamic cortical wave patterns of the signaling marker PIP3 on the substrate-attached surface of electrofused, “giant” *Dictyostelium* cells. We then extracted two KM features (first and second projected finite-time KM averages) and six wave features from the training data to be used during model learning (Fig. S12). The six wave features here included three autocorrelations (temporal, spatial, and global spatial activity) and three probability distributions (temporal firing frequency, spatial firing frequency, and single species signal values). To initiate the optimization, we used the same FHN-2D parameters from figure (Fig. 7e), with representative frames exhibiting oscillations with faster wave speeds and longer propagation distances relative to the PIP3 waves (Fig. 8c, row 1). We then performed one round of KM optimization to match the projected finite-time KM averages of the activator in simulations and the PIP3 signal in giant cells. To do this, we assumed the PIP3 signal to be representative of the FHN activator, U, and optimized a custom objective function that only included error metrics related to the two KM features (Supplementary Methods). The final parameters after round 1 of optimization yielded reconstructions that poorly matched the spatiotemporal features (Fig. 8c, row 2).

Starting from the final parameters after round 1, we then performed a second round of optimization using an objective function with error metrics for only the six wave features (WF), with the three probability distributions weighted less than the three autocorrelations due to the high variability (Supplementary Methods). This optimization resulted in slight improvements with large wave patches, but the dynamics appeared to be significantly slower (Fig. 8c, row 3). Finally, starting from the same final parameters from round 1, we repeated the optimization using an objective function with error metrics for all 8 spatiotemporal features (KM and WF). Surprisingly, this resulted in reconstructions with highly similar dynamics and patterns as observed in the PIP3 training data (Fig. 8c, last row; see also Movie S7). The ability to estimate a coupled stochastic PDE model from a single replicate of noisy microscopy data with low temporal resolution and a single observed species demonstrates the strong robustness of this learning approach.

## Discussion

The development of mathematical models to capture the intricate behaviors of complex systems has a rich history. Traditionally, these models have been knowledge-based, with parameters adjusted through an iterative exploration of parameter space guided by experience and intuition. This method, however, is limited to handcrafted models of low-dimensional systems with few variables and becomes computationally intensive when applied to spatially-distributed systems. Although recent advances in neural networks and data-based learning have offered new possibilities for nonlinear system identification and computational biology, most focus on deterministic PDEs or spatially independent stochastic differential equations, have high computational demands, and often rely on large data with spatiotemporal resolution exceeding system dynamics by orders of magnitude. Additionally, to our knowledge, these approaches have not been applied to excitable reaction-diffusion systems. Such systems are crucial in biological contexts, where signaling networks exhibit noise-driven, dynamic patterns that propagate across cellular boundaries.

Overall, this study introduces a proof-of-concept approach for using limited spatiotemporal data to learn sparse model equations that govern an excitable, stochastic system’s emergent properties and patterns. To this end, we first extracted key features characterizing the datasets: microscale stochastic dynamics and macroscale spatiotemporal patterns. We then estimated model structure and parameters using a two-step machine-learning algorithm that first matches stochastic dynamics using sparse Kramers-Moyal (KM) regression, and second, matches spatiotemporal patterns using wave features (WF) optimization. Using a limited set of wave metrics and a computationally efficient approach for estimating KM averages in a Laplacian-extended state space, our methods demonstrate efficacy over a range of model complexity and data paucity. We validated the method for multiple nonlinear activator-inhibitor models, 1D and 2D spatially-distributed systems, temporally subsampled data, and data with unobserved model species. The effective application of a combined method optimizing for both KM-based and WF-based features using scarce training data from a live-cell imaging experiment conclusively demonstrates the algorithm’s robustness in learning stochastic governing equations from real-world, noisy biological data.

Our approach presents new avenues for understanding not only biochemical waves and pattern formation but also spatially-distributed dynamical systems in general. We learned novel, sparse polynomial reaction-diffusion models for both the FHN and FR excitable systems. From a theoretical perspective, the identification of alternative equations for excitable systems that are simpler than the classical model equations opens the door for further research questions. From an applications perspective, this is essential for understanding real-world systems where the correct model structure is not predetermined, and sparse models can yield more robust predictions and insightful system information. Compared to other methods which focus on learning system trajectories, our approach learns sparse stochastic equations, in which the parameters can provide quantitative, mechanistic insights regarding the excitable system, e.g. reaction model structure, feedback loop strengths, temporal and spatial scales, process noise, and threshold. This presents a significant advantage when modeling dynamic biological systems where learning simplified system structure is just as important as approximating system behavior.

Despite these advances, KM approaches have inherent limitations. Fundamentally, KM averages estimate stochastic dynamics in the limit of infinitesimal time steps and under the assumption of a stationary Markov process. Hence, for systems with rapid dynamics, fitting KM averages works best for datasets that are highly sampled in time. Visualizing the projected finite-time KM averages computed over larger time steps (Fig. S8b–d), we observed decreasing accuracy compared to the standard KM averages, even while keeping data quantity consistent. This effect was less pronounced for the inhibitor due to the absence of process noise in the inhibitor equation here. Optimizing sampling rate for KM-based analysis is also complicated in real-world settings, by the fact that sampling too rapidly can lead to non-Markovian behavior due to physical properties of the system; in these cases, sampling at time steps around the Einstein-Markov time scale is recommended for stochastic modeling (*59, 69*). Additionally, because KM-based methods estimate parameters separately for each species and their SRDE components, the coupling between equations is not accounted for, limiting effectiveness when learning sparse models. There are also often a range of model structures with similar errors in KM quantities that produce dramatically different wave patterns (Fig. 5b, Fig. 6b), highlighting that KM-based stochastic dynamics offer a limited view of system properties. To address these challenges, we developed wave features (WF) optimization.

This Monte Carlo, features-based optimization approach has several advantages. Through the use of a customized objective function, it allows for optimization of a set of key macroscale quantities characterizing system behavior. When KM averages are available as initial parameters for WF optimization, these initial parameters, which approximate stochastic dynamics but deviate significantly in reconstruction patterns, can be effectively tuned to replicate general system features. A key part of the effectiveness lies in the fact that, unlike KM regression where deterministic and stochastic parameters are estimated separately for each species, features-based optimization adjusts the deterministic and stochastic parameters for both activator and inhibitor equations simultaneously.

The strengths of WF optimization are particularly evident when learning sparse models. By jointly tuning parameters of the reaction equation, spatial diffusion coefficients, and noise terms, the algorithm effectively matches wave features (Table S1, Table S2). Even for the sparsest models tested in the FHN and FR models, the 4u2v model and the 5f3r model, respectively, WF optimization recovered wave patterns from initial KM-based reconstructions that were oscillatory (Fig. 5c,e, Fig. 6c). Interestingly, the KM-based and WF-based 4u2v and 5f3r models were all at the Hopf bifurcation point when examining the spatially-homogeneous reaction equation nullclines (insets in Fig. 5c,e, Fig. 6e). This highlights an important distinction from spatially-homogeneous deterministic systems where system behavior can only be tuned by reaction-parameters. WF optimization based models did have slight reaction parameter adjustments for the activator nullcline to approach the training nullclines locally near the system equilibrium (Fig. S2c, Fig. S5c). However, there were also consistent changes in other parameters not visible in phase-plane analysis, such as the spatial diffusion coefficient and state noise parameters, that contributed to the recovery of wave patterns by possibly increasing effective threshold in the system despite being at the Hopf bifurcation (Table S1, Table S2, Fig. S2c, Fig. S5c).

Additionally, using WF optimization to fine-tune parameters for candidate models from sparse KM regression allowed for the identification of novel sparse models for excitable systems. We identified multiple sparse models for both the FHN model and the FR model. For the classical FHN model, which already was a simplified 5u3v model, we found sparser model structures with as few as three activator reaction terms and only one inhibitor term (4u2v, since spatial diffusion adds an extra term for both species). The 4u2v model, after tuning parameters with WF optimization, demonstrated similar stochastic dynamics and wave features to the original model. Similarly, we approximated a biologically-relevant FR model using as few as 4 polynomial terms. Notably, although the FR model contained a rational Hill function, its activator nullcline can be written as a fourth-order polynomial equation. Hence, our library of up to third-order polynomial terms could be used to approximate the nullcline. The sparse models we identified all generally approximated the spatially homogeneous nullclines; however the 4f3r model after WF optimization demonstrated more complicated adjustments (Fig. S5c).

Examining the parameters for these alternative model structures of excitable systems, we observed that the effective positive and negative feedback loops were maintained, indicating that despite the mathematical structure being sparser, the general network structure remained consistent (Table S1, Table S2. Splitting the reaction terms into the signage of the terms that depend on only U, only V, or mixed terms, the reaction network can be inferred. For example, in the 4u2v model structure for the U reaction equation, the V term has a negative coefficient (negative feedback), and the terms that depend on U only (*u*^2^, *u*^3^) yield a positive value in the right quadrant where trajectories lie (Table S1). Our combined approach with approximation of stochastic dynamics and spatiotemporal patterns is thus of high value in modeling biological systems where insights reside in both microscale network structure defined by the stochastic equations as well as the emergent macroscale wave features.

WF optimization also improves estimation from signals with low temporal resolution and unobserved species. In these limited data cases, KM regression yielded poor model estimates (in the case of subsampled data) or was not well-defined (in the case of partially-observed data). Using WF optimization on the other hand, we can tune initial KM-based parameters to obtain approximate models from data with *τ* = 4 s, where temporal resolution is reduced by 99.75% compared to full simulations (Fig. 7b–d). When learning from datasets that resemble experimental data with only one of the two modeled species being observed and a temporal resolution of *τ* = 4 s, WF optimization still provides reasonable parameter estimates, given a set of initial parameters (Fig. 7e). Interestingly, although the activator and inhibitor signals differed significantly in that only the former was modeled with system noise, the optimization procedure converged to similar results. In fact, estimation using the filtered inhibitor signal was slightly poorer based on the wave features in the reconstructions than estimation using the highly noisy activator signal. Additional interesting questions regarding observability of the system arise given that the parameters after optimization using partially-observed data differed significantly from the ground-truth parameters.

The successful application of our approach to a noisy, experimental dataset with high scarcity provides a strong proof-of-concept for the power of this approach. Comparing optimization of projected finite-time KM averages and wave features both separately and together using microscopy data, we validated that the combined optimization of both types of spatiotemporal features yields the best results (Fig. S12, Movie S7). Table S5 shows how the parameters are individually tuned using this dual strategy to learn approximate SRDE models from a noisy, single replicate, low-temporal resolution, dataset of a single observed species in a complex biological network.

The development and validation of these algorithms represent a significant contribution to datadriven learning of spatially-distributed stochastic models of excitable systems. These computational approaches are of interest in our field of cell biology where low resolution, time-lapse imaging of various pattern-forming systems are readily available but often underutilized when constructing models. Future efforts should focus on integrating experimental biology with machine learning and dynamical systems to gain deeper insights into cellular signaling pathways in both health and disease.

## Materials and Methods

### Generating simulations

To generate a training dataset, 50 simulations were obtained for each model in 1D and 2D space (Fig. 1c). Each simulation was carried out by integrating the two-component (activator and inhibitor) stochastic reaction-diffusion equation (SRDE) from 1 over a uniform grid of points in either 1D or 2D space, using the Euler-Maruyama method. The simulations were performed with an integration time step of fi*t* = 0.01 s and ran for a total duration of 400 seconds. Data was saved for all analyses at a “recording” time-step of *τ*_0_ = 0.1 s for FHN simulations and *τ*_0_ = 0.05 s for FR simulations. These simulations were visualized as kymographs for 1D simulations and videos for 2D simulations, as represented in Fig. 1c. For all 1D simulations, 500×1 grid points were used with fi*x* = 1, and for all 2D simulations, a 100×100 grid was used with fi*x* = fi*y* = 1. During training, all spatial points were used for 1D data, and less than 10% of spatial points were used for 2D data. The ground truth SRDE parameters used for all training simulations were obtained from iterative tuning and previous studies and are listed in Table 1 and Table 2. Additional details are described in Supplementary Methods.

### Cell microscopy data

#### Cell culture

The wild-type *Dictyostelium discoideum* cells of axenic AX2 strain (thawed from lab stock; originally obtained from R Kay laboratory, MRC Laboratory of Molecular Biology, UK) were cultured at 22 ^◦^C in HL-5 media that was supplemented with penicillin and streptomycin. Additionally, to ensure stable expression of various constructs, Hygromycin (50 *µ*g/mL) and/or G418 (30 *µ*g/mL) were added to the media according to the resistance conferred by the vectors containing the genes of interest. Cells were kept as adherent culture in petri dishes and subcultured every 2-4 days using appropriate techniques to maintain a healthy confluency of 70-90%. For transfection or electrofusion experiments, cells were transferred to a shaking culture (maintained at 200 rpm) for about 3-4 days. All experiments were conducted within one month of thawing the cells from the frozen stocks.

#### DNA constructs and transfection

The constructs of PHcrac-mCherry (pDM358) and PHcrac-RFP (pDRH) was obtained from the Devreotes lab stock. Axenic AX2 *Dictyostelium* cells were transfected following the standard electroporation protocol. Around 5×10^6^ Ax2 cells were harvested from the shaking culture and pelleted for each transfection. The cells were then washed twice with ice-cold H-50 buffer (20 mM HEPES, 50 mM KCl, 10 mM NaCl, 1 mM MgSO4, 5 mM NaHCO3, 1 mM NaH2PO4, pH adjusted to 7.0), resuspended in 100 *µ*L ice-cold H-50 buffer, and then approximately 1-5 *µ*g of PHcrac plasmid DNA was added to them. The mixture was quickly transferred to an ice-cold 0.1 cm gap cuvette (Bio-Rad, 1652089). The cells in the cuvette were then electroporated twice using a Gene Pulser Xcell Electroporation device, at 0.85 kV voltage and 25 *µ*F capacitance, with a 5-second interval between pulses. Right after electroporation, the cells were incubated on ice inside the cuvette for 5 minutes and then transferred to a 10 cm petri dish containing 10 mL of HL-5 medium, supplemented with heat-killed *Klebsiella aerogenes* bacteria. After 1-2 days of recovery, Hygromycin and/or G418 were added for antibiotic selection.

#### Microscopy

All live-cell experiments were conducted on a stage maintained at ∼22 ^◦^C. Time-lapse images were captured using either a Zeiss LSM 780-FCS Single-point laser scanning confocal microscope (Zeiss Axio Observer with 780-Quasar; 34-channel spectral, high-sensitivity gallium arsenide phosphide detectors) or a Nikon Eclipse Ti-E dSTORM Total Internal Reflection Fluorescence (TIRF) Microscope (equipped with a Photometrics Evolve EMCCD camera). The Zeiss 780 was operated using ZEN Black software, while the Nikon TIRF was controlled with NIS-Elements software. For the Zeiss 780 confocal microscope, a 40X/1.30 Plan-Neofluar oil objective (with appropriate digital zoom) was used, and for the Nikon TIRF, a 100x/1.4 Plan-Apo oil objective was used. In the Zeiss 780 confocal microscope, a 488 nm (Ar laser) was used for GFP and YFP excitation, and a 561 nm (solid-state) laser was used for RFP and mCherry excitation. In the Nikon TIRF, a 488 nm (Ar laser) was used for GFP excitation, and a 561 nm (0.5W fiber laser) was used for mCherry and RFP excitation.

#### Electrofusion

Growth phase *Dictyostelium* cells expressing PHCrac-mCherry or PHCrac-RFP were first harvested from the shaking culture. A total of 1.5 × 10^8^ cells were washed twice and resuspended in 10 mL SB (17 mM Soerensen buffer, 15 mM KH2PO4, and 2 mM Na2HPO4, pH 6.0). The cells were then placed in a conical tube and gently rolled for 30-40 minutes to induce cluster formation. Subsequently, 800 *µ*L of the rolled cells were transferred to a 0.4 cm gap Bio-Rad cuvette using pipette tips with cut-off edges to maintain cluster integrity. Electroporation was performed using a BioRad Gene Pulser (Model 1652098) at 1 kV, 3 *µ*F once, followed by 1 kV, 1 *µ*F twice more to induce membrane hemifusion, with a 3-second interval between pulses. Approximately 35 *µ*L of electrofused cells were then transferred from the cuvette to a Nunc Lab-Tek 8-well chamber. The cells were incubated for 5 minutes in Ca/Mg free media, before adding 450 *µ*L of SB buffer supplemented with 2 mM CaCl2 and 2 mM MgCl2.

#### Live-cell imaging of wave dynamics

To observe the wave dynamics at the substrate-attached surface of the cell membrane, electrofused “giant” *Dictyostelium* cells were used (refer to the “Electrofusion” section for details). After the electrofused cells had recovered, settled, and adhered to the substrate for ∼ 1 hour, live-cell imaging experiments were started. Images were captured at 7-second intervals per frame using either a TIRF microscope or confocal laser scanning microscopes focused on the basal plane at the very bottom surfaces of the cells. Imaging experiments were continued until 150 frames were acquired (about 17 minutes) to monitor the ventral wave dynamics at the membrane-cortex. Microscopy data was then analyzed using a custom MATLAB (MathWorks) script (Supplementary Methods).

### Identifying and learning models using Kramers-Moyal (KM) regression with sparsity enforcement

For model identification, we assume a fixed model structure and estimate the parameters of the deterministic and stochastic components of the stochastic PDE in 1 using data-derived stochastic dynamics (Fig. 1d). For stationary processes with Gaussian white noise, the analytic components of SDEs (the deterministic, or “drift”, term and the stochastic, or “diffusivity”, term) are directly related to data-based conditional moments known as Kramers-Moyal (KM) coefficients, or KM averages (*58, 63*) (Supplementary Methods). There are two KM averages that are computed for each species (activator and inhibitor), and these are linked to the deterministic and stochastic components, respectively, of the SRDE for each species. To estimate the deterministic parameters for the FHN model, we applied linear regression, while for the FR model, we applied nonlinear regression using the Levenberg-Marquardt algorithm. For both the FHN and FR models, we estimated the stochastic parameter of the activator and inhibitor components in the SRDE model directly using the second KM averages.

Model learning involved estimating the reaction model structure as well as all other SRDE parameters using sparse KM regression. A library of up to third-order polynomial reaction terms were used to define an initial set of variable model structures (Fig. 1f), from which a sparse model was learned. Specifically, we considered the following reaction terms for both the activator and inhibitor equations:

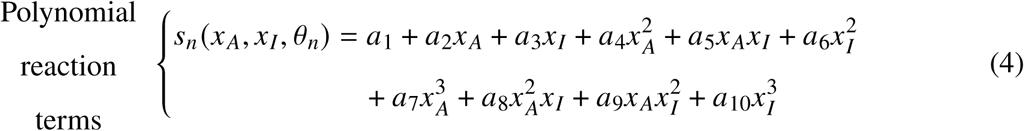

where *0_n_* = {*a*_1_*, a*_2_*, …, a*_10_} denotes the reaction parameters.

In order to learn a sparse polynomial reaction-diffusion model, we used a reverse-greedy search algorithm from (*58*) with modifications (Supplementary Methods). Linear regression was performed iteratively, with elimination of the reaction parameter smallest in absolute value at each step. The spatial diffusion term was retained and only removed at the final iteration. Optimal reaction model sparsity was estimated by minimizing the Bayesian information criterion (BIC), which balances complexity and accuracy (*49,70*). The optimal model structure was determined using the top model parameters retained over all simulations at a given level of sparsity. We also identified candidate sparse model structures by considering models with fewer terms than the estimated optimal sparsity. To assess variability and compare with ground truth parameters, mean and standard deviations of all estimated parameters were calculated over 50 simulations. The estimated parameters were then used to generate an additional 20-50 simulations, constituting a reconstruction dataset (Fig. 1g).

### Quantification and optimization of wave features (WF) for validating and estimating models

A comprehensive, quantitative comparison was performed to evaluate the quality of the reconstructed simulations against the original training dataset. This assessment involved measuring a total of eight wave features: the joint probability distribution of model species, spatial firing frequency, temporal firing frequency, spatial wave activity, and spatial and temporal autocorrelations for both the activator and inhibitor (Supplementary Methods). Together, these metrics of the patterns observed in the simulations provided a rigorous validation of estimated models (Fig. 1e). Wilcoxon rank-sum tests were conducted as a nonparametric test to compare the distribution of wave metrics of reconstruction simulations to that of training simulations.

Using these metrics, we also developed a Monte Carlo optimization scheme to estimate model parameters that generate reconstructions with wave features that best match the wave features in the training data. A Nelder-Mead simplex search was performed to minimize a custom objective function involving the sum of squared relative errors in each wave feature (Supplementary Methods). This WF optimization approach was applied in three cases of model estimation: 1) estimation after sparse KM regression for FHN-1D and FR-1D models; 2) estimation for FHN-2D models from temporally subsampled data; and 3) estimation for FHN-2D models temporally subsampled data only observing the activator or the inhibitor. After optimization, to assess the quality of parameter estimates in each case, 20 reconstructions were generated, and wave features were analyzed.

### Extracting and optimizing spatiotemporal features from live-cell imaging data

Because live-cell imaging data involved observation of only a single species, PIP3, which we modeled as the activator, we had no experimental data for the inhibitor to extract conditional moments used in KM averages. In order to estimate stochastic dynamics, we thus used what we refer to as projected finite-time KM averages, which are computed in a two-dimensional (instead of three-dimensional) binned Laplacian-extended state space consisting of only PIP3 (activator) and its Laplacian. Additionally, we computed six wave features based on the PIP3 signal: three probability distributions (single species probability distribution, temporal firing frequency, and spatial firing frequency) and three autocorrelations (temporal autocorrelation, spatial autocorrelation, and global spatial activity autocorrelation). During optimization, we used the Monte Carlo optimization scheme with the same form of the custom objective function as used with simulation datasets, but with weighting of the autocorrelation metrics more than the probability distribution metrics. To achieve improved learning, we additionally tested cases of including two KM metrics for the stochastic dynamics in the objective function: the root sum of squared errors (RSSE) of the first and second projected finite-time KM averages in the Laplacian-extended state space. Details are provided in Supplementary Methods.

## Acknowledgments

We thank Y. Kevrekidis (School of Engineering, Johns Hopkins University (JHU)) and M.I. Miller (School of Engineering, JHU) for their input and advice as thesis committee members. We thank A. Titi (School of Engineering, JHU) for stimulating discussions. We thank all members of the Devreotes, Iglesias, Kevrekidis, and D.N. Robinson (School of Medicine, JHU) labs for their helpful feedback.

## Funding

This work was supported by the following grants: NIH R35 GM118177 (PND), NIH T32 GM136577 Medical Scientist Training Program (JHU), AFOSR MURI FA95501610052 (PND), NIH R01-GM149073 (PAI), as well as NIH S10 OD016374 (S. Kuo of the JHU Microscope Facility).

## Author contributions

BAS, PND, and PAI conceptualized the overall study. BAS developed the code and conducted the simulations and analysis. BAS and TB conducted microscopy experiments. ChatGPT-4o (OpenAI) was used only to edit some of the initial writing and to vectorize a MAT-LAB script for computing Kramers-Moyal averages. BAS, TB, PAI, and PND wrote the manuscript. PAI and PND supervised the study.

## Competing interests

Authors declare that they have no competing interests.

## Data and materials availability

All data are available in the main text or the supplementary materials. Code is available on Github:

## Supplementary materials

Supplementary Methods Figs. S1 to S12

Tables S1 to S5 References (*71–73*) Movies S1 to S7

## Notes

### Competing Interest Statement

The authors have declared no competing interest.

### Summary of Updates

Figure 1 revised. Figure 8 added. S7 video added. Author added.

## References

1. D. Cottrell, P. S. Swain, P. F. Tupper, Stochastic branching-diffusion models for gene expression. Proc. Natl. Acad. Sci. U. S. A. 109 (25), 9699–9704 (2012), doi:10.1073/pnas.1201103109.

2. K. Takahashi, S. Tanase-Nicola, P. R. ten Wolde, Spatio-temporal correlations can drastically change the response of a MAPK pathway. Proc. Natl. Acad. Sci. U. S. A. 107 (6), 2473–2478 (2010), doi:10.1073/pnas.0906885107.

3. B. A. Bicknell, P. Dayan, G. J. Goodhill, The limits of chemosensation vary across dimensions. Nat. Commun. 6 (1), 7468 (2015), doi:10.1038/ncomms8468.

4. A. N. Landge, B. M. Jordan, X. Diego, P. Müller, Pattern formation mechanisms of self-organizing reaction-diffusion systems. Dev. Biol. 460 (1), 2–11 (2020), doi:10.1016/j.ydbio.2019.10.031.

5. A. De Masi, P. A. Ferrari, J. L. Lebowitz, Reaction-diffusion equations for interacting particle systems. J. Stat. Phys. 44 (3-4), 589–644 (1986), doi:10.1007/bf01011311.

6. M. Doi, Stochastic theory of diffusion-controlled reaction. J. Phys. A Math. Gen. 9 (9), 1479–1495 (1976), doi:10.1088/0305-4470/9/9/009.

7. 7. M. Nagasawa, Stochastic processes in quantum physics, Monographs in Mathematics (Springer, Basel, Switzerland), 2000 ed. (2012).

8. L. Roques, O. Bonnefon, Modelling population dynamics in realistic landscapes with linear elements: A mechanistic-statistical reaction-diffusion approach. PLoS One 11 (3), e0151217 (2016), doi:10.1371/journal.pone.0151217.

9. C. Cosner, Reaction–diffusion equations and ecological modeling, in *Lecture Notes in Mathematics*, Lecture notes in mathematics (Springer Berlin Heidelberg, Berlin, Heidelberg), pp. 77–115 (2008), doi:10.1007/978-3-540-74331-6_3.

10. N. Ahmed, et al., A dynamical study on stochastic reaction diffusion epidemic model with nonlinear incidence rate. Eur. Phys. J. Plus 138 (4), 350 (2023), doi:10.1140/epjp/s13360-023-03936-z.

11. A. D. Corlan, J. Ross, Kinetics methods for clinical epidemiology problems. Proc. Natl. Acad. Sci. U. S. A. 112 (46), 14150–14155 (2015), doi:10.1073/pnas.1510927112.

12. V. Volpert, S. Petrovskii, Reaction-diffusion waves in biology. Phys. Life Rev. 6 (4), 267–310 (2009), doi:10.1016/j.plrev.2009.10.002.

13. V. E. Deneke, S. Di Talia, Chemical waves in cell and developmental biology. J. Cell Biol. (2018), doi:10.1083/jcb.201701158.

14. H. L. Swinney, V. I. Krinsky, eds., *Wave and patterns in chemical and biological media* (MIT Press, Cambridge, Mass.) (1991).

15. P. Gray, S. K. Scott, Sustained oscillations and other exotic patterns of behavior in isothermal reactions. J. Phys. Chem. 89 (1), 22–32 (1985), doi:10.1021/j100247a009.

16. W. Jahnke, W. E. Skaggs, A. T. Winfree, Chemical vortex dynamics in the BelousovZhabotinskii reaction and in the two-variable oregonator model. J. Phys. Chem. 93 (2), 740–749 (1989), doi:10.1021/j100339a047.

17. M. Falcke, Reading the patterns in living cells —the physics of ca2+ signaling. Adv. Phys. 53 (3), 255–440 (2004), doi:10.1080/00018730410001703159.

18. A. T. Winfree, The Geometry of Biological Time (Springer New York) (2001), doi:10.1007/ 978-1-4757-3484-3.

19. R. Kapral, K. Showalter, Chemical Waves and Patterns (Springer Science & Business Media) (2012).

20. E. M. Izhikevich, Dynamical Systems in Neuroscience (MIT Press) (2007).

21. J. Keener, J. Sneyd, eds., Mathematical Physiology, vol. 1: Cellular Physiology of Interdisciplinary Applied Mathematics (Springer New York) (2009), doi:10.1007/978-0-387-75847-3.

22. P. A. Iglesias, Excitable systems in cell motility, in 52nd IEEE Conference on Decision and Control (2013), pp. 757–762, doi:10.1109/CDC.2013.6759973.

23. T. Bretschneider, et al., The three-dimensional dynamics of actin waves, a model of cytoskeletal self-organization. Biophys. J. 96 (7), 2888–2900 (2009), doi:10.1016/j.bpj.2008.12.3942.

24. Y. Arai, et al., Self-organization of the phosphatidylinositol lipids signaling system for random cell migration. Proceedings of the National Academy of Sciences 107 (27), 12399–12404 (2010), doi:10.1073/pnas.0908278107.

25. W. M. Bement, et al., Activator-inhibitor coupling between Rho signalling and actin assembly makes the cell cortex an excitable medium. Nat. Cell Biol. 17 (11), 1471–1483 (2015), doi: 10.1038/ncb3251.

26. O. D. Weiner, W. A. Marganski, L. F. Wu, S. J. Altschuler, M. W. Kirschner, An actin-based wave generator organizes cell motility. PLoS Biol. 5 (9), 2053–2063 (2007), doi: 10.1371/journal.pbio.0050221.

27. L. B. Case, C. M. Waterman, Adhesive F-actin waves: a novel integrin-mediated adhesion complex coupled to ventral actin polymerization. PLoS One 6 (11), e26631 (2011), doi:10.1371/journal.pone.0026631.

28. M. G. Vicker, Eukaryotic cell locomotion depends on the propagation of self-organized reaction-diffusion waves and oscillations of actin filament assembly. Exp. Cell Res. 275 (1), 54–66 (2002), doi:10.1006/excr.2001.5466.

29. A. M. Winans, S. R. Collins, T. Meyer, Waves of actin and microtubule polymerization drive microtubule-based transport and neurite growth before single axon formation. Elife 5, e12387 (2016), doi:10.7554/eLife.12387.

30. V. Marchesin, G. Montagnac, P. Chavrier, ARF6 promotes the formation of Rac1 and WAVE-dependent ventral F-actin rosettes in breast cancer cells in response to epidermal growth factor. PLoS One 10 (3), e0121747 (2015), doi:10.1371/journal.pone.0121747.

31. M. Wu, X. Wu, P. De Camilli, Calcium oscillations-coupled conversion of actin travelling waves to standing oscillations. Proc. Natl. Acad. Sci. U. S. A. 110 (4), 1339–1344 (2013), doi: 10.1073/pnas.1221538110.

32. J.-M. Yang, et al., Integrating chemical and mechanical signals through dynamic coupling between cellular protrusions and pulsed ERK activation. Nat. Commun. 9 (1), 4673 (2018), doi: 10.1038/s41467-018-07150-9.

33. T. Banerjee, et al., Spatiotemporal dynamics of membrane surface charge regulates cell polarity and migration. Nat. Cell Biol. (2022), doi:10.1038/s41556-022-00997-7.

34. Y. Miao, et al., Wave patterns organize cellular protrusions and control cortical dynamics. Mol. Syst. Biol. 15 (3), e8585 (2019), doi:10.15252/msb.20188585.

35. S. Matsuoka, M. Ueda, Mutual inhibition between PTEN and PIP3 generates bistability for polarity in motile cells. Nat. Commun. 9 (1), 4481 (2018), doi:10.1038/s41467-018-06856-0.

36. D. Taniguchi, et al., Phase geometries of two-dimensional excitable waves govern selforganized morphodynamics of amoeboid cells. Proc. Natl. Acad. Sci. U. S. A. 110 (13), 5016–5021 (2013), doi:10.1073/pnas.1218025110.

37. P. N. Devreotes, et al., Excitable Signal Transduction Networks in Directed Cell Migration. Annu. Rev. Cell Dev. Biol. 33 (1), annurev–cellbio–100616–060739 (2017), doi:10.1146/annurev-cellbio-100616-060739.

38. A. J. Ridley, et al., Cell migration: integrating signals from front to back. Science 302 (5651), 1704–1709 (2003), doi:10.1126/science.1092053.

39. Y. Miao, et al., Altering the threshold of an excitable signal transduction network changes cell migratory modes. Nat. Cell Biol. 19 (4), 329–340 (2017), doi:10.1038/ncb3495.

40. H. Zhan, et al., An Excitable Ras/PI3K/ERK Signaling Network Controls Migration and Oncogenic Transformation in Epithelial Cells. Dev. Cell 54 (5), 608–623.e5 (2020), doi: 10.1016/j.devcel.2020.08.001.

41. C.-H. Huang, M. Tang, C. Shi, P. A. Iglesias, P. N. Devreotes, An excitable signal integrator couples to an idling cytoskeletal oscillator to drive cell migration. Nat. Cell Biol. 15 (11), 1307–1316 (2013), doi:10.1038/ncb2859.

42. S. Fukushima, S. Matsuoka, M. Ueda, Excitable dynamics of Ras triggers spontaneous symmetry breaking of PIP3 signaling in motile cells. J. Cell Sci. 132 (5) (2019), doi:10.1242/jcs. 224121.

43. S. Flemming, F. Font, S. Alonso, C. Beta, How cortical waves drive fission of motile cells. Proc. Natl. Acad. Sci. U. S. A. 117 (12), 6330–6338 (2020), doi:10.1073/pnas.1912428117.

44. D. S. Pal, et al., Actuation of single downstream nodes in growth factor network steers immune cell migration. Dev. Cell 58 (13), 1170–1188.e7 (2023), doi:10.1016/j.devcel.2023.04.019.

45. Y. Lin, et al., Ras suppression potentiates rear actomyosin contractility-driven cell polarization and migration. Nat. Cell Biol. 26 (7), 1062–1076 (2024), doi:10.1038/s41556-024-01453-4.

46. C. Shi, C.-H. Huang, P. N. Devreotes, P. A. Iglesias, Interaction of motility, directional sensing, and polarity modules recreates the behaviors of chemotaxing cells. PLoS Comput. Biol. 9 (7), e1003122 (2013), doi:10.1371/journal.pcbi.1003122.

47. T. Banerjee, et al., A dynamic partitioning mechanism polarizes membrane protein distribution. Nat. Commun. 14 (1), 7909 (2023), doi:10.1038/s41467-023-43615-2.

48. G. E. Karniadakis, et al., Physics-informed machine learning. Nature Reviews Physics 3 (6), 422–440 (2021), doi:10.1038/s42254-021-00314-5.

49. S. L. Brunton, J. N. Kutz, *Data-driven science and engineering: Machine learning, dynamical systems, and control* (Cambridge University Press (Virtual Publishing), Cambridge, England) (2019), doi:10.1017/9781108380690.

50. S. L. Brunton, J. L. Proctor, J. N. Kutz, Discovering governing equations from data by sparse identification of nonlinear dynamical systems. Proc. Natl. Acad. Sci. U. S. A. 113 (15), 3932–3937 (2016), doi:10.1073/pnas.1517384113.

51. S. H. Rudy, S. L. Brunton, J. L. Proctor, J. N. Kutz, Data-driven discovery of partial differential equations. Sci Adv 3 (4), e1602614 (2017), doi:10.1126/sciadv.1602614.

52. F. Van Breugel, Y. Liu, B. W. Brunton, J. N. Kutz, PyNumDiff: A Python package for numerical differentiation of noisy time-series data. J. Open Source Softw. 7 (71), 4078 (2022), doi:10. 21105/joss.04078.

53. B. de Silva, et al., PySINDy: A Python package for the sparse identification of nonlinear dynamical systems from data. J. Open Source Softw. 5 (49), 2104 (2020), doi:10.21105/joss.02104.

54. K. Kaheman, J. N. Kutz, S. L. Brunton, SINDy-PI: a robust algorithm for parallel implicit sparse identification of nonlinear dynamics. Proc. Math. Phys. Eng. Sci. 476 (2242), 20200279 (2020), doi:10.1098/rspa.2020.0279.

55. A. Yazdani, L. Lu, M. Raissi, G. E. Karniadakis, Systems biology informed deep learning for inferring parameters and hidden dynamics. PLoS Comput. Biol. 16 (11), e1007575 (2020), doi:10.1371/journal.pcbi.1007575.

56. C. Rao, et al., Encoding physics to learn reaction–diffusion processes. *Nat*. Mach. Intell. 5 (7), 765–779 (2023), doi:10.1038/s42256-023-00685-7.

57. F. Dietrich, et al., Learning effective stochastic differential equations from microscopic simulations: Linking stochastic numerics to deep learning. Chaos 33 (2), 023121 (2023), doi: 10.1063/5.0113632.

58. L. Boninsegna, F. Nüske, C. Clementi, Sparse learning of stochastic dynamical equations. J. Chem. Phys. 148 (24), 241723 (2018), doi:10.1063/1.5018409.

59. J. L. Callaham, J.-C. Loiseau, G. Rigas, S. L. Brunton, Nonlinear stochastic modelling with Langevin regression. Proc. Math. Phys. Eng. Sci. 477 (2250), 20210092 (2021), doi:10.1098/rspa.2021.0092.

60. S. J. Lade, Finite sampling interval effects in Kramers–Moyal analysis. Phys. Lett. A 373 (41), 3705–3709 (2009), doi:10.1016/j.physleta.2009.08.029.

61. R. Friedrich, et al., Extracting model equations from experimental data. Phys. Lett. A 271 (3), 217–222 (2000), doi:10.1016/S0375-9601(00)00334-0.

62. R. Friedrich, C. Renner, M. Siefert, J. Peinke, Comment on “indispensable finite time corrections for Fokker-Planck equations from time series data”. Phys. Rev. Lett. 89 (14) (2002), doi:10.1103/physrevlett.89.149401.

63. S. Siegert, R. Friedrich, J. Peinke, Analysis of data sets of stochastic systems. Phys. Lett. A 243 (5), 275–280 (1998), doi:10.1016/S0375-9601(98)00283-7.

64. G. Amselem, M. Theves, A. Bae, E. Bodenschatz, C. Beta, A stochastic description of Dictyostelium chemotaxis. PLoS One 7 (5), e37213 (2012), doi:10.1371/journal.pone.0037213.

65. R. Fitzhugh, Impulses and Physiological States in Theoretical Models of Nerve Membrane. Biophys. J. 1 (6), 445–466 (1961), doi:10.1016/s0006-3495(61)86902-6.

66. J. Nagumo, S. Arimoto, S. Yoshizawa, An Active Pulse Transmission Line Simulating Nerve Axon. Proceedings of the IRE 50 (10), 2061–2070 (1962), doi:10.1109/JRPROC.1962.288235.

67. S. Bhattacharya, P. A. Iglesias, The Regulation of Cell Motility Through an Excitable Network. IFAC-PapersOnLine 49 (26), 357–363 (2016), doi:10.1016/j.ifacol.2017.03.001.

68. S. Bhattacharya, P. A. Iglesias, The threshold of an excitable system serves as a control mechanism for noise filtering during chemotaxis. PLoS One 13 (7), e0201283 (2018), doi: 10.1371/journal.pone.0201283.

69. R. Friedrich, J. Peinke, M. Sahimi, M. Reza Rahimi Tabar, Approaching complexity by stochastic methods: From biological systems to turbulence. Phys. Rep. 506 (5), 87–162 (2011), doi: 10.1016/j.physrep.2011.05.003.

70. T. Hastie, R. Tibshirani, J. Friedman, The Elements of Statistical Learning: Data Mining, Inference, and Prediction (Springer, New York, NY), 2 ed. (2009).

71. K. Jacobs, Stochastic Processes for Physicists: Understanding Noisy Systems (Cambridge University Press) (2010), doi:10.1017/CBO9780511815980.

72. E. J. Candes, J. Romberg, T. Tao, Robust uncertainty principles: exact signal reconstruction from highly incomplete frequency information. IEEE Trans. Inf. Theory 52 (2), 489–509 (2006), doi:10.1109/TIT.2005.862083.

73. W.-X. Wang, R. Yang, Y.-C. Lai, V. Kovanis, C. Grebogi, Predicting catastrophes in nonlinear dynamical systems by compressive sensing. Phys. Rev. Lett. 106 (15), 154101 (2011), doi: 10.1103/PhysRevLett.106.154101.

