## Supplemental Materials for "Learning stochastic reaction-diffusion models from limited data using spatiotemporal features"

### Supplementary Materials for
### Learning stochastic reaction-diffusion models from limited data using spatiotemporal features

Bedri Abubaker-Sharif, Tatsat Banerjee, Peter N. Devreotes, Pablo A. Iglesias*

**This PDF file includes:**

Supplementary Methods

Figures S1 to S12

Tables S1 to S5

Captions for Movies S1 to S7

**Other Supplementary Materials for this manuscript:**

Movies S1 to S7

#### 1.1 Supplementary Methods

##### 1.1.1 Generating simulations

To generate a training dataset, 50 simulations were obtained for each model in a discretized 1D and 2D space, as described in the main text. An example of the implementation for the activator component of the stochastic reaction-diffusion equation (SRDE) is shown here for the FHN model:

$$\begin{aligned} u_i(t+\Delta t) &= u_i(t) + \Delta t\left(D_u L_u(t) + s_u\big(u_i(t), v_i(t), \theta_u\big)\right) + g_u\sqrt{\Delta t}\mathcal{N}(0,1) \\ v_i(t+\Delta t) &= v_i(t) + \Delta t\left(D_v L_v(t) + s_v\big(u_i(t), v_i(t), \theta_v\big)\right) + g_v\sqrt{\Delta t}\mathcal{N}(0,1) \end{aligned} \tag{E1}$$

where

$$L_u(t) = \frac{u_{i+1}(t) - 2u_i(t) + u_{i-1}(t)}{\Delta x^2} \tag{E2}$$

is the second-order central-difference approximation of the 1D-Laplacian of $u$ with periodic boundary conditions ($L_v$ is defined similarly); $s_u$, $s_v$ define the activator and inhibitor reaction terms; and $\theta_u$ and $\theta_v$ are the reaction parameters. Subscript $i$ refers to the spatial position, and $t$ refers to time. Integration time step is $\Delta t = 0.01$ s and spatial grid points have $\Delta x = \Delta y = 1$, as mentioned in the main text. Note that the additive noise term incorporates an independent and identically distributed standard Gaussian random variable over all spatial and time points.

For reconstruction datasets, simulations were obtained using the same integration scheme with estimated SRDE parameters. Constraints were also placed on the minimum and maximum signal values ($\pm 10$) at any grid point to retain the ability to visualize any reconstruction and limit optimization errors arising from divergent trajectories.

##### 1.1.2 Image processing for live-cell microscopy data

For pre-processing, image sequences of PIP3 signals (150 frames total, acquired every 7 seconds) were regionally cropped and smoothed with 2D median and Gaussian filtering using a custom MATLAB script. Each frame was then spatially downsampled to a $100 \times 100$ spatial grid in order to match the 2D space used for simulations. The dataset was normalized using min-max normalization (minimum and maximum set to 1st and 99th percentile signal values, respectively) and finally split into three equal parts, each consisting of 50 frames (t = 0 to t = 343 seconds), to generate a training set.

##### 1.1.3 Deriving Kramers-Moyal (KM) averages for spatial Langevin equations

As described in the main text, we considered two-component, activator-inhibitor models (activator $x_A$, inhibitor $x_I$) implemented as spatial Langevin equations in the following Itô differential form:

$$\begin{aligned} dx_A &= \left(D_A \nabla^2 x_A + s_A(x_A, x_I, \theta_A)\right) dt + g_A dQ_A \\ dx_I &= \left(D_I \nabla^2 x_I + s_I(x_A, x_I, \theta_I)\right) dt + g_I dQ_I \end{aligned} \tag{E3}$$

To avoid complexities related to deriving Fokker-Planck (FP) equations (which are the basis of KM terms) for stochastic partial differential equations in continuous time and space, we considered the finite-difference approximation of the spatial Langevin equations used in the integration scheme. Here, over a discretized spatial domain, the trajectories at each grid point evolve according to SDEs and corresponding FP equations with identical structure and parameters (*71*). By defining a state vector for each spatial point, $\mathbf{x} = \left[\begin{smallmatrix} x_A \\ x_I \end{smallmatrix}\right] = \left[\begin{smallmatrix} x_1 \\ x_2 \end{smallmatrix}\right]$, with a corresponding Laplacian vector, $L\mathbf{x} = \left[\begin{smallmatrix} Lx_A \\ Lx_I \end{smallmatrix}\right] = \left[\begin{smallmatrix} \nabla^2 x_A \\ \nabla^2 x_I \end{smallmatrix}\right]$, we have the following SDE in vector form at each spatial grid point:

$$\begin{aligned} &\mathbf{dx} = \mathbf{J}(\mathbf{x}, L\mathbf{x}, \Theta_{\text{det.}})dt + \mathbf{G}(\Theta_{\text{stoch.}})\mathbf{dQ} \\ &\mathbf{J} = \begin{bmatrix} D_A Lx_A + s_A(\mathbf{x}, \theta_A) \\ D_I Lx_I + s_I(\mathbf{x}, \theta_I) \end{bmatrix}, \quad \mathbf{G} = \text{diag}(g_A, g_I), \quad \mathbf{dQ} = \begin{bmatrix} dQ_A \\ dQ_I \end{bmatrix} \end{aligned} \tag{E4}$$

where the deterministic and stochastic model parameters are given by $\Theta_{\text{det.}} = \{D_A, \theta_{s_A}, D_I, \theta_{s_I}\}$ and $\Theta_{\text{stoch.}} = \{g_A, g_I\}$, respectively. In general, $\mathbf{G}$ is an $N \times M$ matrix, where $N$ is the number of species and $M$ is the number of mutually independent noise terms; and $\mathbf{dQ}$ is $M \times 1$. We approximate $\nabla^2$ using second order central differences resulting in dependencies on the two neighboring grid points in 1D, and 4 neighboring grid points in 2D.

Based on the finite-difference approximation, the spatial diffusion term at a particular grid point can be viewed as a known external input upon which we can condition. For the SDE at each grid point in equation E4, the corresponding approximate FP equation at each grid point, with conditioning on the approximate Laplacian, is:

$$\frac{\partial P(\mathbf{x}, t | L\mathbf{x})}{\partial t} = -\sum_{i=1}^{2} \frac{\partial}{\partial x_i} \left[\mathbf{H}_i^{(1)} P\right] + \sum_{i=1}^{2} \sum_{j=1}^{2} \frac{\partial^2}{\partial_{x_i} \partial_{x_j}} \left[\mathbf{H}_{ij}^{(2)} P\right] \tag{E5}$$

where $P(\mathbf{x}, t | L\mathbf{x})$ is a time evolution of the scalar conditional probability distribution for the state vector at a particular grid point, and $\mathbf{H}^{(1)} = \begin{bmatrix} H_1^{(1)} \\ H_2^{(1)} \end{bmatrix} = \begin{bmatrix} H_A^{(1)} \\ H_I^{(1)} \end{bmatrix}$ and matrix $\mathbf{H}^{(2)} = \text{diag}\left(H_A^{(2)}, H_I^{(2)}\right)$

are the empirically estimated Kramers-Moyal coefficients, or KM averages, of the Fokker-Planck equation.

Using this extension to discretized, spatially-distributed systems, the first and second KM averages in the FP equation relate directly to the analytic drift (deterministic component) and diffusivity (stochastic component), respectively, for the SDE in equation E4 as follows:

$$\mathbf{H}^{(1)}(\mathbf{x}, L\mathbf{x}) = \widehat{\mathbf{H}}^{(1)}(\mathbf{x}, L\mathbf{x}, \Theta_{\text{det.}}) = \mathbf{J}(\mathbf{x}, L\mathbf{x}, \Theta_{\text{det.}}) \tag{E6}$$

$$\mathbf{H}^{(2)} = \widehat{\mathbf{H}}^{(2)}(\Theta_{\text{stoch.}}) = \tfrac{1}{2}\mathbf{G}\mathbf{G}^{T} \tag{E7}$$

where $\widehat{\mathbf{H}}^{(1)}$ and $\widehat{\mathbf{H}}^{(2)}$ denote the parameterized, theoretical KM averages. Because we assume constant noise parameters, $\widehat{\mathbf{H}}^{(2)}$ is a diagonal $2\times2$ matrix. Computationally, the data-derived, empirical KM averages ($H_n^{(1)}$ and $H_n^{(2)}$) approximate the theoretical KM averages ($\widehat{H}_n^{(1)}$ and $\widehat{H}_n^{(2)}$) for the $n$-th species as follows:

$$H_n^{(1)}(\mathbf{x}, Lx_n) \approx \widehat{H}_n^{(1)}(\mathbf{x}, Lx_n, \Theta_{n,\text{det.}}) = D_n L x_n + s_n(\mathbf{x}, \theta_n) \tag{E8}$$

$$H_n^{(2)} \approx \widehat{H}_n^{(2)}(\Theta_{n,\text{stoch.}}) = \tfrac{1}{2} g_n^2 \tag{E9}$$

where $\Theta_{n,\text{det.}} = \{D_n, \theta_n\}$ and $\Theta_{n,\text{stoch.}} = \{g_n\}$ correspond to parameters of the $n$-th component in E3; and $\widehat{H}_n^{(2)}$ denotes the $n$-th diagonal entry of $\widehat{\mathbf{H}}^{(2)}$. The optimization problem (described later) then becomes finding the set of parameters for which the right-hand side (RHS) theoretical KM averages best approximate the left-hand side (LHS) empirical KM averages.

In the limit of infinitesimal time, the empirical KM averages converge to the theoretical KM averages in equations E8 and E9. The first empirical KM average for the $n$-th species, $H_n^{(1)}$, is here defined as:

$$H_n^{(1)}(\mathbf{x}_0, L_{n0}) = \lim_{\tau \to 0} \frac{1}{\tau} m_{n,\tau}^{(1)}(\mathbf{x}_0, L_{n0}), \tag{E10}$$

$$m_{n,\tau}^{(1)}(\mathbf{x}_0, L_{n0}) = \mathbf{E}\left[ x_n(t+\tau) - x_n(t) \middle| \mathbf{x}(t) = \mathbf{x}_0, Lx_n(t) = L_{n0} \right] \tag{E11}$$

where $\mathbf{x} = \begin{bmatrix} x_A \\ x_I \end{bmatrix}$ and $Lx_n$ is the Laplacian approximation in E2 for the $n$-th species (with initial values $\mathbf{x}_0$ and $Ln_0$, respectively, over a time step $\tau$); and $m_{n,\tau}^{(1)}$ is the finite-time first moment. Similarly, the second KM average for the $n$-th species, $H_n^{(2)}$, is here defined as:

$$H_n^{(2)} = \lim_{\tau\to 0} \frac{1}{2\tau} m_{n,\tau}^{(2)} \quad \text{(E12)}$$

$$m_{n,\tau}^{(2)} = \mathbf{E}\left[(x_n(t+\tau) - x_n(t))^2\right] \quad \text{(E13)}$$

where $m_{n,\tau}^{(2)}$ is the finite-time second moment computed over all bins. Note that for the second KM average, conditioning is not necessary since the stochastic component of the SRDE is a state-independent constant; thus, $m_{n,\tau}^{(2)}$ and $H_n^{(2)}$ are scalar constants.

##### 1.1.4 Computing and visualizing stochastic dynamics with Kramers-Moyal (KM) averages

Overall, the empirical KM averages, $H_n^{(1)}$ and $H_n^{(2)}$ in E10 and E12, estimate the average change and average squared change, respectively, in the $n$-th species over an infinitesimal time step. We compute the first KM average through conditioning on initial values of three key dependent variables in the deterministic component of E3: the two species and the Laplacian of the $n$-th species, which form a region in 3D space which we refer to as the Laplacian-extended state space. In practice, the conditioning in E11 was performed using binned initial values $\mathbf{x}_0$ and $Ln_0$ (for $\mathbf{x}_n$ and $Lx_n$, respectively). Note that although conditioning can be implemented based on the neighboring points used in the Laplacian approximation, we instead directly condition on the Laplacian values for computational efficiency. Additionally, although the KM averages here are derived based on FP equations at individual grid points, the conditional expectations can be estimated over all spatiotemporal points since the data at all points follow the same SDE/FP model.

Kramers-Moyal analysis began with extracting a few additional quantities from the raw simulation data sampled at the recording time step of $\tau = \tau_0$. Given the time-series signal for the $n$-th species at a given spatial point, $x_n(t)$, we computed 1) the approximate Laplacian, $Lx_n(t)$, using second-order central differences, and 2) the forward finite-difference, $dx_n(t) = x_n(t+\tau) - x_n(t)$, over all spatiotemporal points, excluding the first and last time points and the edge spatial points to ignore boundary conditions. For 1D spatial simulations, we used all 498 interior spatial points (500 total grid points – 2); for 2D spatial simulations, to reduce computational time, we used data from every 3$^{\text{rd}}$ row and column in the interior spatial grid, corresponding to 1089 spatial points ($\sim \frac{1}{9} \times 10,000$ total grid points).

Next, we vectorized the three signals for each species—$x_n(t)$, $Lx_n(t)$, $dx_n(t)$—by stacking the time-series from all selected spatial points, ensuring that the rows (time points) remained consistent across all signals. To facilitate the estimation of conditional moments, we discretized the state variable and Laplacian signals, $x_n(t)$ and $Lx_n(t)$, using a binning scheme that uniformly divided their range into a fixed number of intervals (bins). Each bin corresponded to a specific range of signal values, and we applied this scheme to both the FHN and FR datasets. For the binning process, we used 50 bins for the FHN data and 150 bins for the FR data, chosen to balance accuracy and computational efficiency. The bin ranges were defined to extend 10–15% beyond the minimum and maximum signal values obtained from the initial simulation, ensuring that all possible values across the 50 stochastic simulations were covered.

To compute the conditional expectations $m^{(1)}_{n,\tau}$ and $m^{(2)}_{n,\tau}$ for the $n$-th species, we treated the three signals $x_A(t), x_I(t), Lx_n(t)$ as specifying a point in $\mathbb{R}^3$, a Laplacian-extended state space, and discretized this space using a 3D grid of bins as defined above. Each axis of the grid corresponded to one of the signals, and the total number of bins was the product of the bin counts along each axis. For the FHN data, this resulted in a grid of $50 \times 50 \times 50$ bins, while for the FR data, we used a finer grid of $150 \times 150 \times 150$ bins. For each time point (row), we assigned the values of $x_A(t)$, $x_I(t)$, and $Lx_n(t)$ to a specific bin in the 3D grid based on which intervals they fell into for each signal. Rows corresponding to the same bin were grouped together, and the values of $dx_n(t)$ for a given bin were averaged to estimate the finite-time first moments for each species, as described in equation E11. Similarly, the finite-time second moments were estimated by squaring the values of $dx_n(t)$ across all rows (irrespective of binning) and averaging them, as shown in equation E13. These finite-time first and second moments, $m^{(1)}_{n,\tau}$ and $m^{(2)}_{n,\tau}$, were computed separately for each simulation and for each species, at four multiples of $\tau_0$: $\tau = \tau_0, 2\tau_0, 3\tau_0, 4\tau_0$

Finally, to compute the empirical KM averages $H^{(1)}_n$ and $H^{(2)}_n$, we estimated the finite-time KM averages for this set of four finite-time conditional moments. These finite-time KM averages correspond to E10 and E12, without taking the limits: the finite-time moments, $m^{(1)}_{n,\tau}$ and $m^{(2)}_{n,\tau}$, multiplied by $\frac{1}{\tau}$ or $\frac{1}{2\tau}$, respectively. The finite-time KM averages were defined on the same binned Laplacian-extended state space, giving us up to four values of the finite-time KM averages at each bin (stochastic trajectories accessed only a subset of bins in the 3D space). We applied 1D spline extrapolation using MATLAB's `interp1` function to estimate the limiting values at bins with at least three available finite-time KM averages. Rather than computing limits as $\tau \longrightarrow 0$, we used $\tau \longrightarrow 0.01$ (the integration time step) to obtain the final empirical KM averages.

For visualization of the empirical KM averages defined on the discretized 3D Laplacian-extended state space, we averaged values along the Laplacian dimension and plotted the "projected" empirical KM averages as a heatmap in the 2D activator-inhibitor phase plane.

##### 1.1.5 Parameter estimation given a fixed model structure using Kramers-Moyal (KM) regression

To estimate parameters given a fixed SDE model structure with KM regression, we performed the following general steps for each species in a single simulation:

1. Compute finite-time conditional moments, $m_{n,\tau}^{(1)}$ and $m_{n,\tau}^{(2)}$ in E11 and E13, from stochastic trajectories of spatiotemporal data sampled at a series of time steps $\tau = \{\tau_0, 2\tau_0, 3\tau_0, 4\tau_0\}$, where $\tau_0$ is the recording time-step, mentioned previously.

2. Estimate empirical KM averages, $H_n^{(1)}$ and $H_n^{(2)}$ in E10 and E12, in the limit of infinitesimal time steps using spline extrapolation.

3. Apply least-squares regression to find the set of SRDE parameters, $\Theta_{n,\text{det.}}$ and $\Theta_{n,\text{stoch.}}$, for which the RHS theoretical KM averages in E8 and E9 best approximate the LHS empirical KM averages, using a fixed model structure.

Step 3 involved model identification of the optimal parameters for a given structure of the deterministic and stochastic components of the SRDE. This parameter set was obtained by minimizing the mean squared error (MSE) between the empirical KM averages in equations E10 and E12 and the parameterized theoretical KM averages in equations E8 and E9, as follows:

$$J_n^{(1)}(\Theta_{n,\text{det.}}) = \frac{1}{N} \sum_{\mathbf{x}_0, L_{n0}} \left( H_n^{(1)}(\mathbf{x}_0, L_{n0}) - \widehat{H}_n^{(1)}(\mathbf{x}_0, L_{n0}, \Theta_{n,\text{det.}}) \right)^2 \tag{E14}$$

$$J_n^{(2)}(\Theta_{n,\text{stoch.}}) = \left( H_n^{(2)} - \widehat{H}_n^{(2)}(\Theta_{n,\text{stoch.}}) \right)^2 \tag{E15}$$

where $J_n^{(1)}$ and $J_n^{(2)}$ are the cost functions (MSE) for the first and second KM averages of the $n$-th species; the sum in E14 is computed over the binned values of $\mathbf{x}_0$ and $Ln_0$ in the Laplacian-extended state space; $N$ is the total number of bins over which the sum is computed; the MSE in E15 is simply a squared error since the second KM averages are scalars; and $\Theta_{n,\text{det.}}$ and $\Theta_{n,\text{stoch.}}$ correspond to parameters in the deterministic and stochastic terms, respectively, for the $n$-th species in equation E3. The full procedure described above was repeated for each simulation in a training dataset in order to obtain statistics on parameter sets.

When applying KM regression on temporally subsampled data ($\tau = 0.1, 1, 4$ s), we combined data from multiple simulations to generate a single KM average and single parameter estimate, while maintaining a consistent data volume. Specifically, finite-time moments in step 1 were computed from one simulation for $\tau = 0.1$ s, from 10 simulations for $\tau = 1$ s, and from 40 simulations for $\tau = 4$ s. Since extrapolation procedures fail with large time steps, finite-time empirical KM averages were calculated using the standard KM average formulas in E10 and E12, without taking the limit. Using these finite-time empirical KM averages, computed separately for each time step $\tau$, we applied the same regression procedures to generate parameter estimates for each sampling rate.

###### 1.1.6 Optimizing stochastic dynamics using KM regression

Model identification was achieved by finding the optimal set of parameters for the deterministic and stochastic components of the SRDE given a model structure. This parameter set was obtained by minimizing the MSE in E14 and E15. We estimated the optimal deterministic parameters for each component of the SRDE using least squares regression. Two different regression approaches were employed: 1) for the FHN model, linear regression with basis functions derived from the exact terms in the model, or 2) for the FR model, nonlinear regression with the exact nonlinear terms in the FR model.

For both linear and nonlinear regression, we vectorized $\Theta_{n,\text{det.}}$, $H_n^{(1)}$, and $\widehat{H}_n^{(1)}$ (denoted as $\boldsymbol{\beta}_n$, $\mathbf{h}_n^{(1)}$, and $\widehat{\mathbf{h}}_n^{(1)}$, respectively for clarity) to obtain equation E16 below. For linear regression, we additionally converted E16 to matrix form as follows:

$$J_n^{(1)}(\boldsymbol{\beta}_n) = \frac{1}{N}\left(\mathbf{h}_n^{(1)} - \widehat{\mathbf{h}}_n^{(1)}(\boldsymbol{\beta}_n)\right)^T \left(\mathbf{h}_n^{(1)} - \widehat{\mathbf{h}}_n^{(1)}(\boldsymbol{\beta}_n)\right) \tag{E16}$$

$$= \frac{1}{N}\left(\mathbf{h}_n^{(1)} - \mathbf{Z}_n\boldsymbol{\beta}_n\right)^T \left(\mathbf{h}_n^{(1)} - \mathbf{Z}_n\boldsymbol{\beta}_n\right) \tag{E17}$$

Here, $\mathbf{Z}_n$ is the design matrix for the $n$-th species, with columns containing the exact terms in the deterministic model structure for the $n$-th component of the SRDE. Each row of $\mathbf{Z}_n$ matches the rows of the vector $\mathbf{h}_n^{(1)}$, identifying a unique point in the binned 3D Laplacian-extended state space (e.g. for the FHN model, $\{u_0, v_0, Ln_0\}$); specifically, the rows of the design matrix correspond to values of the terms in the model structure evaluated at a particular point in this discretized space. For example, $\mathbf{Z}_n$ and $\boldsymbol{\beta}_n$ for the activator species in the FHN model were constructed as:

$$\mathbf{Z}_u = \begin{bmatrix} | & | & | & | & | \\ \mathbf{Lu}_0 & \mathbf{u}_0 & \mathbf{v}_0 & \mathbf{u}_0^2 & \mathbf{u}_0^3 \\ | & | & | & | & | \end{bmatrix} \quad \text{and} \quad \boldsymbol{\beta}_u = \left[D_u, a_1, a_2, a_3, a_4\right]^T \tag{E18}$$

where the FHN model parameters in $\boldsymbol{\beta}_u$ correspond to the deterministic SRDE parameters in E3. Similar design matrices and parameter vectors were constructed for the inhibitor species.

To guarantee the restricted isometry property before solving the optimization problem (*72*), we rewrote E17 with normalized columns of $\mathbf{Z}'_n$ and an adjusted parameter vector $\boldsymbol{\beta}'_n$, as done in (*73*):

$$J_n^{(1)}(\boldsymbol{\beta}_n) = J_n^{(1)}(\boldsymbol{\beta}'_n) = \frac{1}{N}\left(\mathbf{h}_n^{(1)} - \mathbf{Z}'_n\boldsymbol{\beta}'_n\right)^T\left(\mathbf{h}_n^{(1)} - \mathbf{Z}'{}_n\boldsymbol{\beta}'_n\right) \ , \tag{E19}$$

$$\text{where } \mathbf{Z}'_n = \mathbf{Z}_n\mathbf{M}_n^{-1}, \quad \boldsymbol{\beta}'_n = \mathbf{M}_n\boldsymbol{\beta}_n \ ,$$

$$\text{and } \mathbf{M}_n = \text{diag}\{m_{n,i}\}, \quad m_{n,i} = \text{L2-norm of i-th column of } \mathbf{Z}_n$$

For the $n$-th species, the operations involving the diagonal matrix $\mathbf{M}_n$ produced the following effects without changing the overall cost function in E14: the $i$-th column of the adjusted $\mathbf{Z}'_n$ is calculated as the $i$-th column of the original $\mathbf{Z}_n$ divided by its corresponding L2-norm ($m_{n,i}$); and the $i$-th component of $\boldsymbol{\beta}'_n$ is calculated as the $i$-th component of the original $\boldsymbol{\beta}_n$ multiplied by $m_{n,i}$.

Finally, in the linear regression case, deterministic parameters of the SRDE for the $n$-th species were obtained by minimizing the MSE in equation E19. The solution of the normal equation provided the optimal parameters $\boldsymbol{\beta}_n^{'*}$ in terms of the adjusted design matrix ($\mathbf{Z}'_n$); the final deterministic SRDE parameter estimates, $\boldsymbol{\beta}_n^*$, based on the original design matrix ($\mathbf{Z}_n$) were then estimated as follows:

$$\boldsymbol{\beta}_n^{'*} = \left(\mathbf{Z}_n'^T\mathbf{Z}'_n\right)^{-1}\mathbf{Z}'_n\mathbf{h}_n^{(1)} \tag{E20}$$

$$\boldsymbol{\beta}_n^* = \mathbf{M}_n\boldsymbol{\beta}_n^{'*} \tag{E21}$$

where $\mathbf{h}_n^{(1)}$ represents the vectorized empirical KM averages, and $\mathbf{M}_n$ is defined as above based on the original design matrix for the $n$-th species.

In the case of nonlinear regression with FR data, we used MATLAB's built-in `nlinfit` function based on the Levenberg-Marquardt nonlinear least squares algorithm to find optimal parameters for the deterministic component of the SRDE for each species. This minimized the MSE in equation E16, which involved terms that were nonlinear in the parameters as seen in the FR model structure.

For both the FHN and FR datasets, we estimated the stochastic parameters for both the activator and inhibitor components in the SRDE model by setting E15 equal to 0 and directly solving for $g_n$, as follows:

$$\Theta^*_{n,\text{stoch.}} = g^*_n = \underset{g_n}{\operatorname{argmin}} \left( H^{(2)}_n - \frac{1}{2} g^2_n \right)^2 = \sqrt{2H^{(2)}_n} \tag{E22}$$

##### 1.1.7 Learning model structure and parameters from a library of models using sparse KM regression

A library of up to third order polynomial reaction terms were used to define an initial set of model structures, from which a sparse model was learned. This library of reaction terms together with a fixed Laplacian term for spatial diffusion defined the deterministic component of the SRDE and the theoretical first KM averages for both species in E8. Specifically, we considered the following reaction terms for both the activator and inhibitor equations:

$$\text{Polynomial reaction terms} \begin{cases} s_n(x_A, x_I, \theta_n) = a_1 + a_2 x_A + a_3 x_I + a_4 x_A^2 + a_5 x_A x_I + a_6 x_I^2 \\ \qquad + a_7 x_A^3 + a_8 x_A^2 x_I + a_9 x_A x_I^2 + a_{10} x_I^3 \\ \text{where } \theta_n = \{a_1, a_2, ..., a_{10}\} \end{cases} \tag{E23}$$

In order to learn a sparse reaction-diffusion model, we used a reverse-greedy search algorithm from (*58*) with some modifications. After completing steps 1 and 2 described previously, in step 3, instead of using a fixed model structure, we applied linear regression iteratively using the expanded polynomial reaction-diffusion library as basis functions. These basis functions, defining the deterministic component of the SRDE and forming the theoretical first KM averages for each species, were then fit the empirical first KM averages.

We solved the normal equation with 11 columns in the design matrix (one Laplacian term and 10 polynomial terms) at iteration 0. At each subsequent iteration, we eliminated the reaction parameter smallest in absolute value along with its corresponding column in the design matrix and solved the normal equation. The spatial diffusion parameter and its corresponding column in the design matrix were maintained regardless of parameter magnitude to enforce a reaction-diffusion model structure, and thus were the last to be removed at iteration 11. Sparse KM regression was applied separately to each of the 50 training simulations. Given the variability across simulations, the model parameters retained at each iteration also varied. This distribution in estimated model structures was measured by calculating the proportion of simulations in which a specific model term in the library remained selected at each iteration.

Sparse model learning involved two components: 1) learning the optimal sparsity (number of parameters, $k^*$) and 2) learning the exact terms in the reaction model to use at optimal sparsity (specific parameters, $\theta^*$). The optimal reaction model sparsity for the $n$-th species' equation was determined by minimizing the Bayesian information criterion (BIC), which balances model sparsity and model accuracy (*49, 70*). Under a Gaussian model with likelihood $\mathcal{L}$ and sum of squared errors (SSE), BIC is defined as: $\text{BIC} = \log(N)k - 2\log(\mathcal{L})$, where $\log(\mathcal{L}) = -(N/2)\log(2\pi) - (N/2)\log(\sigma^2) - \text{SSE}/(2\sigma^2)$. Approximating the error variance, $\sigma^2$, with the MSE, we estimated the BIC at each iteration of sparse KM regression as follows:

$$\text{BIC} = \log(N)k + N\left(\log(2\pi) + \log(\text{MSE}) + 1\right) \tag{E24}$$

where $k$ is the number of remaining parameters, $N$ is the total number of bins in the MSE, and $\text{MSE} = \frac{1}{N}\text{SSE} = J_n^{(1)}$ from E14.

The optimal sparsity for each simulation was estimated as the number of parameters at the BIC minimum. A distribution of the optimal sparsity from all simulations was computed, and the mode was selected as the final estimate of optimal sparsity, $k^*$. Next, to identify the specific parameters (or model terms) to use at this optimal sparsity, we used the distribution in estimated model structures previously computed. Specifically, the optimal model structure was determined using the top $k^*$ model parameters retained in the distribution at iteration $j^* = 11 - k^*$.

This approach was used to estimated reaction-diffusion model structure together with model parameters for the deterministic component of both the activator and inhibitor SRDEs. The stochastic component of the SRDEs was estimated as a constant parameter, as before. We also identified candidate sparse model structures by considering model structures with $k < k^*$ for both the activator and inhibitor equations.

##### 1.1.8 Computing wave features to quantify spatiotemporal patterns

For any given 1D or 2D spatial simulation, the following eight wave features were extracted to quantify spatiotemporal patterns:

1. State probability distribution: joint probability distribution of the activator and inhibitor.
2. Temporal firing frequency: probability distribution of the no. of firings observed over time.
3. Spatial firing frequency: probability distribution of the no. of firings observed over space.
4. Spatial wave activity: proportion of spatial pixels above threshold over time.
5. Activator temporal autocorrelation: average activator patterns over time.
6. Inhibitor temporal autocorrelation: average inhibitor patterns over time.
7. Activator spatial autocorrelation: average activator patterns over space.
8. Inhibitor spatial autocorrelation: average inhibitor patterns over space.

To assess the quality of reconstructions, these quantities were estimated for each of the simulations in both training and reconstruction datasets. Quantification was done using all spatial pixels from the last 300 s of every simulation. All simulations were sampled at $\tau = 1$ s for wave feature analysis. Once each wave feature was computed for a given simulation, the quantities were averaged to provide a high-dimensional, mean wave feature characterizing the dataset. Standard deviations were usually computed as well to measure variability. Quantification details can be found below.

Joint probability distributions (feature 1) were computed for each simulation from the activator and inhibitor signals over all available spatiotemporal points. The probability distributions were computed over a discretized 2D state space, with 70 uniformly-spaced bins for the activator and inhibitor signals. The range of signals in the first training simulation, with 15% additional extension beyond the minimum and maximum, was used to construct a consistent binning scheme for all simulations while capturing variability in the system.

Wave features 2-4 involved counts based on a pre-defined threshold of the inhibitor signal. Temporal firing frequency (feature 2) at a given spatial pixel was estimated based on the number of times the inhibitor signal over time surpassed a fixed threshold level (defined as signal more than

15% of the range above baseline from simulation 1 in the training data). Counts were measured from time-series at each spatial pixel to generate a distribution of temporal firings for a given simulation. Spatial firing frequency (feature 3) at a given time point was estimated based on the number of times the inhibitor signal over space surpassed the threshold. Counts were measured from signal along the spatial dimension at each time point to generate a distribution of spatial firings for a given simulation. For 1D simulations, there was only 1 spatial dimension; for 2D simulations, signals were extracted only along rows and columns (no signals along other directions). Spatial wave activity (feature 4) was estimated as the proportion of spatial pixels in which the inhibitor signal surpassed threshold and was measured at each time point for a given simulation.

Wave features 5-8 involved evaluating the autocorrelation function on signals over time or space and averaging. To evaluate temporal patterns, the autocorrelation function was computed for activator/inhibitor time-series signals from a given spatial pixel. The values were then averaged over all spatial pixels to give the temporal autocorrelation of the activator/inhibitor for a given simulation (features 5 and 6). To evaluate spatial patterns, the autocorrelation function was computed for activator/inhibitor signals along the spatial dimension at a given time point. As before, for autocorrelation analysis on 2D simulations, signals were extracted only along rows and columns. The autocorrelation values were then averaged over all time points to give the spatial autocorrelation of the activator/inhibitor for a given simulation (features 7 and 8).

Next, to generate wave metrics for each simulation, the 8 raw wave features (WF) above were converted from high-dimensional quantities to 8 scalar quantities grouped in a wave metric vector, $\omega = [\omega_{\mathrm{wf}}^{(1)}, ..., \omega_{\mathrm{wf}}^{(8)}]^T$, as follows: feature 1) Kullback-Leibler (KL) divergence between the joint probability distribution of a given simulation and the average joint probability distribution of the full training dataset; features 2, 3) mean of the distribution of temporal/spatial firing frequency for a given simulation; feature 4) spatial wave activity averaged over time points for a given simulation; features 5-8) root mean squared error between a given simulation's average temporal/spatial autocorrelation and the average temporal/spatial autocorrelation of the full training dataset.

Using these wave metrics, we computed summary metrics for each simulation. Two types of summary metrics were used to compare wave features in reconstructions vs. training data: 1) normalized wave metrics and 2) wave metrics error. For the $i$-th simulation in a dataset (training or reconstruction) with wave metric $\omega_i$, the normalized wave metric $\hat{\omega}_i$ was computed by elementwise division, as follows: $\hat{\omega}_i^{(j)} = \omega_i^{(j)} / \overline{\omega}_{\text{train}}^{(j)}$, where $\overline{\omega}_{\text{train}}$ is the average 8-component wave metric for the training dataset, and $j$ refers to a component of the wave metric vector. For the $i$-th simulation in a dataset, the wave metrics error, $e(\omega_i)$, was defined as the root of the sum of squared relative errors, as follows:

$$e(\omega_i) = \sqrt{\sum_j \left( \frac{\overline{\omega}_{\text{train}}^{(j)} - \omega_i^{(j)}}{\overline{\omega}_{\text{train}}^{(j)} + c} \right)^2} \tag{E25}$$

where $j$ refers to a component of the wave metric vector and $c = 0.005$ for avoiding division by zero. Wilcoxon rank-sum tests were conducted as a nonparametric test to compare the distribution of summary wave metrics of reconstruction simulations to that of training simulations.

##### 1.1.9 Parameter estimation from simulation data using wave features optimization

We developed a Monte Carlo optimization scheme to estimate model parameters that generated reconstructions that best matched the wave features in the training data. This approach was applied in three parameter estimation cases: 1) estimation after sparse KM regression for FHN-1D and FR-1D models; 2) estimation from temporally subsampled FHN-2D data; 3) estimation from FHN-2D data when only observing either the activator or the inhibitor. After optimization, to assess the quality of parameter estimates in each case, 20 reconstructions were generated and wave features analyzed as described in section 1.1.8.

For case 1, we initialized the algorithm by using the optimal SRDE parameters identified from sparse KM regression for a given candidate 1D spatial reaction-diffusion model structure. Using MATLAB's native fminsearch to perform a Nelder-Mead simplex search, we optimized a custom objective function involving the sum of squared relative errors in the average wave metrics of a reconstruction dataset ($\overline{\omega}_{\text{recon}}$) relative to the average wave metrics of the training dataset ($\overline{\omega}_{\text{train}}$). To evaluate the objective function at each iteration, 20 simulations with identical spatial and temporal length scales as that of training data (400 s duration, over a $100 \times 100$ grid) were generated via parallel computation to form a reconstruction dataset, and 8 wave features with corresponding 8-component wave metric vector were computed for each simulation over a 300 s window, as described previously. The reconstruction dataset wave metric is itself a function of the SRDE parameters that generated it, and so we defined a custom objective function using the sum of squared relative errors from E25 as follows:

$$J(\Theta_A, \Theta_I) = \left(e\left(\overline{\omega}_{\text{recon}}(\Theta_A, \Theta_I)\right)\right)^2 = \sum_j \left(\frac{\overline{\omega}_{\text{train}}^{(j)} - \overline{\omega}_{\text{recon}}^{(j)}}{\overline{\omega}_{\text{train}}^{(j)} + c}\right)^2 \tag{E26}$$

where $j$ refers to a component of the wave metric, $c = 0.005$ to avoid division by zero, and $\Theta_A$ and $\Theta_I$ refer to the full SRDE parameters (deterministic and stochastic components) for the activator and inhibitor in E3. Optimization of this objective function was performed over the full set of deterministic (reaction-diffusion) and stochastic model parameters. We conducted a first round of optimization using initial parameters from sparse KM regression, with multiple runs of 100 iterations conducted in parallel (~3-5 computers). The parameter set that minimized E26 after one round were then used to initialize two to three additional rounds of optimization. During these rounds, to facilitate the search process, we also incorporated runs with initial parameters modified with 1% Gaussian noise. The final estimated parameters were the ones that minimized the custom objective function after all rounds.

For case 2, we performed wave features optimization using initial parameter estimates from finite-time KM regression. These initial parameters were computed as described previously for a fixed FHN model structure using 2D-spatial data sampled at three time steps ($\tau = 0.1, 1, 4$ s). Similar to before, we computed 8 wave features and a wave metric for each simulation in the training data; however, we computed wave metrics separately for data sampled every $\tau = 1$ s and 4 s. Additionally, given the volume of 2D spatial data, we limited wave feature analysis to a 200 s time window, and only used spatial pixels from every third row and column. Using the same objective function, we then optimized over the set of reaction, diffusion, and noise parameters of a fixed FHN-2D model structure. Unlike case 1, however, given the computational cost of 2D-spatial simulations, we generated only 1-2 reconstructions (duration 200 - 300 s) at each iteration of optimization. Wave metrics for the reconstructions were also computed over a limited set of spatiotemporal points, as done with training data. For initialization, parameters from three cases of finite-time KM regression (0.1, 1, or 4 s sampled data) were used, and wave features optimization results compared. For initial parameters based on 0.1 or 1 s data, wave metrics were computed on 1 s sampled reconstructions at each iteration and were optimized to training data wave features at $\tau = 1$ s using E26; similarly, for initial parameters based on 4 s sampled data, wave metrics were computed from and optimized to 4 s sampled data.

For case 3, we used wave features optimization to estimate FHN-2D model parameters when only one of the modeled species was observed every 4 s. The same wave features optimization approach developed for 2D-spatial data in case 2 above was applied, with the following adjustments. Since KM parameters were not available, we used initial knowledge-based FHN-2D activator/inhibitor equation parameters for an oscillatory system that produced larger waves, distinct from that of the training data. Additionally, wave features and wave metrics for training data were computed based only on a single observed species, sampled every 4 s. The joint probability distribution and the average temporal and spatial autocorrelation of the unobserved species were no longer available, resulting in a 5-component single-species wave metric. During optimization, the single-species wave metric was computed for reconstructions sampled at 4 s and matched to the single-species wave metric computed for training data sampled at 4 s by minimizing E26.

###### 1.1.10 Computing spatiotemporal features from live-cell imaging data

In order to improve learning when using limited live-cell imaging data, we extracted two types of spatiotemporal features, stochastic dynamics and wave features, as follows:

Stochastic dynamics:

1. First projected finite-time KM averages: indirect measure of deterministic “drift” in SRDE
2. Second projected finite-time KM averages: indirect measure of stochastic “diffusivity” in SRDE

Wave features:

1. State probability distribution: probability distribution of the activator.
2. Temporal firing frequency: probability distribution of the no. of firings observed over time.
3. Spatial firing frequency: probability distribution of the no. of firings observed over space.
4. Activator temporal autocorrelation: average activator patterns over time.
5. Activator spatial autocorrelation: average activator patterns over space.
6. Global spatial activity autocorrelation: average patterns of global spatial activity over time.

These were computed for each of the time-series data in the PIP3 training dataset as well as any reconstructions during optimization. As a measure of stochastic dynamics for live-cell imaging data with only a single observed species (PIP3, modeled as the activator), we estimated empirical "projected" finite-time KM averages, which are computed in a two-dimensional (instead of three-dimensional) binned Laplacian-extended state space consisting of only the observed species (activator, $x_A$) and its Laplacian ($Lx_A$). These empirical quantities were computed for both the first and second finite-time KM averages. Similar to E10, but conditioning on only the observed species and its Laplacian and without taking limits, the first projected finite-time KM average of the activator, $H^{(1)}_{A,\tau,\text{proj}}$, was defined as follows:

$$H^{(1)}_{A,\tau,\text{proj}}(x_{A0}, L_{A0}) = \frac{1}{\tau} m^{(1)}_{A,\tau}(x_{A0}, L_{A0}) \tag{E27}$$

$$m^{(1)}_{A,\tau,\text{proj}}(x_{A0}, L_{A0}) = \mathbf{E}\left[x_A(t+\tau) - x_A(t) \middle| x_A(t) = x_{A0}, Lx_A(t) = L_{A0}\right] \tag{E28}$$

The second projected finite-time KM average of the activator, $H^{(2)}_{A,\tau,\text{proj}}$, was similarly defined using E12 with conditioning to relax the assumption of state-independent noise and without taking limits:

$$H^{(2)}_{A,\tau,\text{proj}}(x_{A0}, L_{A0}) = \frac{1}{2\tau} m^{(2)}_{A,\tau}(x_{A0}, L_{A0}) \tag{E29}$$

$$m^{(2)}_{A,\tau,\text{proj}}(x_{A0}, L_{A0}) = \mathbf{E}\left[(x_A(t+\tau) - x_A(t))^2 \middle| x_A(t) = x_{A0}, Lx_A(t) = L_{A0}\right] \tag{E30}$$

We note that in both sets of the above equations, $\tau = 7$ s, based on the acquisition time step of the PIP3 training data.

In order to facilitate optimization, we converted these two KM averages to KM metrics, similar to how raw wave features were converted to wave metrics. This was important since an analytical form for theoretical projected finite-time KM averages was not available in order to define a cost function as before in E14 and E15. The KM metrics were defined as the root sum of squared errors (RSSE) of the KM values from a given spatiotemporal time-series in a dataset to the mean KM values from the PIP3 training data.

$$\omega^{(1)}_{\text{km}} = \sqrt{\sum_{x_{A0}, L_{A0}} \left(\overline{H}^{(1)}_{A,\tau,\text{proj}}(x_{A0}, L_{A0}) - \widehat{\overline{H}}^{(1)}_{A,\tau,\text{proj}}(x_{A0}, L_{A0})\right)^2} \tag{E31}$$

$$\omega^{(2)}_{\text{km}} = \sqrt{\sum_{x_{A0}, L_{A0}} \left(\overline{H}^{(2)}_{A,\tau,\text{proj}}(x_{A0}, L_{A0}) - \widehat{\overline{H}}^{(2)}_{A,\tau,\text{proj}}(x_{A0}, L_{A0})\right)^2} \tag{E32}$$

where $\overline{H}^{(1)}_{A,\tau,\mathrm{proj}}$ and $\overline{H}^{(2)}_{A,\tau,\mathrm{proj}}$ are the average of the empirical KM values computed using E27 and E29 for each spatiotemporal time-series in the PIP3 training dataset, and similarly $\overline{\widehat{H}}^{(1)}_{A,\tau,\mathrm{proj}}$ and $\overline{\widehat{H}}^{(2)}_{A,\tau,\mathrm{proj}}$ are the average of the empirical KM values computed for each spatiotemporal simulation in a reconstruction dataset.

These summations in E31 and E32 were computed over the union of the set of points $\{x_{A0}, L_{A0}\}$ in the binned space accessed by the KM values from a given dataset (set 1) and the mean KM values of the training data (set 2). The KM values at points that were only accessed by one of the two sets were defined to be 0 in the opposing set (e.g. KM values at a point accessed only in set 1 would be defined as 0 in set 2). Importantly, because the activator (PIP3) values from training data were based on normalized signals (see 1.1.2), the activator values from FHN-2D models were also computed based on similarly normalized signals. This resulted in KM values occupying a similar region in the Laplacian-extended state space. Finally, the two scalar error quantities for the first and second projected finite-time KM averages were grouped in a two component KM metric vector, $\omega_{\mathrm{km}} = [\omega^{(1)}_{\mathrm{km}}, \omega^{(2)}_{\mathrm{km}}]^T$.

Next, we computed six wave features based on the PIP3/activator signal. Three involved probability distributions similar to those listed previously in section 1.1.8: 1) single species probability distribution, 2) distribution of temporal firing frequency, and 3) distribution of spatial firing frequency. Feature 1 was computed similar to feature 1 in section 1.1.8 but using a single species instead of the joint distribution. Features 2 and 3 were computed as described previously. The next three wave features involved autocorrelations: 4) temporal autocorrelation, 5) spatial autocorrelation, and 6) global spatial activity autocorrelation. Features 4 and 5 were computed as described previously. Feature 6, defined as the autocorrelation of the spatial wave activity over time described in section 1.1.8, was a new wave feature providing a measure of global system behavior.

As before, to generate wave metrics, the 6 raw wave features were converted from high-dimensional quantities to scalar quantities grouped in a 6-component wave metric vector, $\omega_{\mathrm{wf}} = [\omega^{(1)}_{\mathrm{wf}}, ..., \omega^{(6)}_{\mathrm{wf}}]^T$, as follows: feature 1) Kullback-Leibler (KL) divergence between the activator probability distribution of a given simulation and the average PIP3 probability distribution of the training dataset; features 2, 3) KL divergence between the distribution of temporal/spatial firing frequency for a given simulation and the average temporal/spatial firing frequency distribution of the PIP3 training dataset; feature 4-6) root sum of squared errors between a given simulation's average autocorrelation and the average autocorrelation of the PIP3 training dataset.

##### 1.1.11 Parameter estimation from live-cell imaging data using spatiotemporal features optimization

We used a Monte Carlo optimization scheme to estimate FHN-2D model parameters from live-cell imaging data. Here, we identified parameters that generated SRDE reconstructions that best matched the eight spatiotemporal metrics of the PIP3 training data described in section 1.1.10 (two KM metrics and six WF metrics). This approach was applied in three estimation rounds based on optimizing a custom objective function involving different features: round 1) only projected finite-time KM averages (KM), round 2A) only wave features (WF), and round 2B) both (KM+WF). After optimization, to assess the quality of parameter estimates over the full range of spatiotemporal features (not just the features being optimized), 20 reconstructions were generated using the optimal parameter estimates from each round of optimization and all 8 spatiotemporal features were analyzed as described in section 1.1.10.

Using MATLAB's native fminsearch to perform a Nelder-Mead simplex search, we optimized a custom objective function involving the sum of squared relative errors in the average spatiotemporal metrics of a reconstruction dataset ($\overline{\omega}_{\text{recon}}$) relative to the average spatiotemporal metrics of the training dataset ($\overline{\omega}_{\text{train}}$). To evaluate the objective function at each iteration, 3 to 6 simulations were generated via parallel computation to form a reconstruction dataset with identical spatial length scales as that of training data (100×100 grid) and a temporal duration at least that of training data (343 s). For feature extraction, various spatiotemporal features (depending on the estimation case) were computed with a corresponding spatiotemporal metric vector, $\omega_i$, for the $i$-th simulation as described in section 1.1.10. We note that the features were computed using all spatial points and over a 343 s time window to match that of the PIP3 training data. The overall spatiotemporal metric vector of the reconstruction dataset, $\overline{\omega}_{\text{recon}}$, was then computed as the average of these $\omega_i$.

As previously, this spatiotemporal metric is itself a function of the SRDE parameters that generated the reconstructions. However, here, we defined a weighted variation of the previously used objective function based on the sum of squared relative errors as follows:

$$J(\Theta_A, \Theta_I) = \sum_j \left( \alpha_j \frac{\overline{\omega}^{(j)}_{\text{train}} - \overline{\omega}^{(j)}_{\text{recon}}}{\overline{\omega}^{(j)}_{\text{train}} + c} \right)^2 \tag{E33}$$

where $j$ refers to a component of the spatiotemporal metric, $\alpha_j$ refers to a weighting factor for each component, $c = 0.0001$ to avoid division by zero, and $\Theta_A$ and $\Theta_I$ refer to the full SRDE parameters (deterministic and stochastic components) for the activator and inhibitor in E3. Optimization of this objective function was performed over the full set of deterministic (reaction-diffusion) and stochastic model parameters.

For round 1 of optimization, only stochastic dynamics were used for optimization, and so feature extraction involved only the first and second projected finite-time KM averages. Hence, for the $i$-th simulation in a reconstruction dataset, $\omega_i = \omega_{\text{km}}$, a two-component vector as defined in section 1.1.10. Additionally, we set $\alpha_j = 1$ to have equal weighting for both KM values. During this first round of optimization, multiple (at least 3) runs of 100 iterations were conducted in parallel to minimize the objective function in E33. At each objective function evaluation, 3 to 6 simulations were generated, and spatiotemporal metrics were computed over a 343 s time window. The starting point for the time window ranged from 7 s to 100 s, in order to skip initial frames during which system activity was biased by the lack of inhibitor values. We initialized the algorithm by using the same initial knowledge-based FHN-2D model parameters for an oscillatory system used in section 1.1.9. For initial conditions of the activator, we used the normalized signals from the initial frames in the PIP3 training dataset. For initial conditions of the inhibitor at each spatial point at time 0, we set values to either 0, or the values of the inhibitor from solving the spatially-homogeneous FHN-2D inhibitor nullcline equation based on the available parameter set.

The parameter set that minimized E33 after round 1 was then used to initialize both round 2A and round 2B optimization. During round 2A, only wave features were used for optimization; hence, for the $i$-th simulation in a reconstruction dataset generated during optimization, $\omega_i = \omega_{\text{wf}}$, a 6-component vector as defined in section 1.1.10. Because of the variability of the probability distributions in the PIP3 training data and the sensitivity of the KL divergence, we weighted the probability-based wave metrics (components 1-3) less than the autocorrelation-based wave metrics (components 4-6). To do this, we set $\alpha_j = 0.1$ for $j = 1, 2, 3$ and $\alpha_j = 1$ for $j = 4, 5, 6$. Similar to round 1, multiple (at least 3) runs of 100 iterations were conducted in parallel to minimize the objective function in E33.

During round 2B, both KM values and wave features were used for optimization. This meant that for the $i$-th simulation in a reconstruction dataset generated during optimization, $\omega_i =$

$[\omega_{\text{km}}^{(1)}, \omega_{\text{km}}^{(2)}, \omega_{\text{wf}}^{(1)}, ..., \omega_{\text{wf}}^{(6)}]^T$, an 8-component vector concatenating KM and WF metrics defined in section 1.1.10. To compare this optimization to round 2A in which only WF metrics were used for optimization, we initialized round 2B similarly with the optimal parameters identified from round 1. Likewise, we weighted the probability-based wave metrics less than the other metrics (KM metrics and autocorrelation-based WF metrics). To do this, we set $\alpha_j = 0.1$ for $j = 3, 4, 5$ and $\alpha_j = 1$ for $j = 1, 2, 6, 7, 8$. Similar to round 2A, multiple (at least 3) runs of 100 iterations were conducted in parallel to minimize the objective function in E33.

### Supplementary Figures

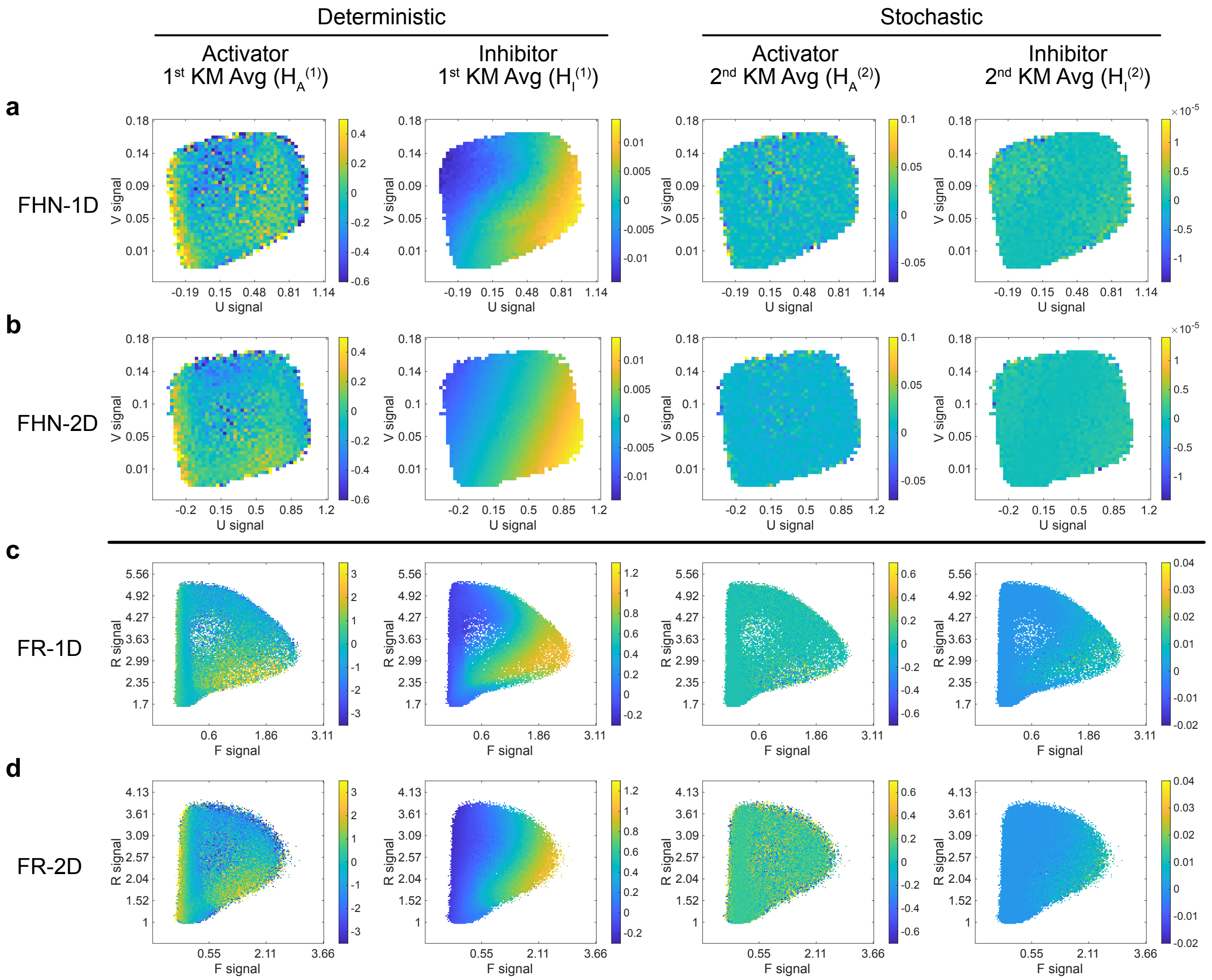


**Figure S1**: **Visualizing KM averages for FHN and FR datasets.** KM averages computed over a Laplacian-extended state space from FHN-1D/2D and FR-1D/2D training data, and then averaged over Laplacian values and plotted as a heatmap in the activator-inhibitor phase plane. Computations from a single simulation in each of the four training sets are shown.

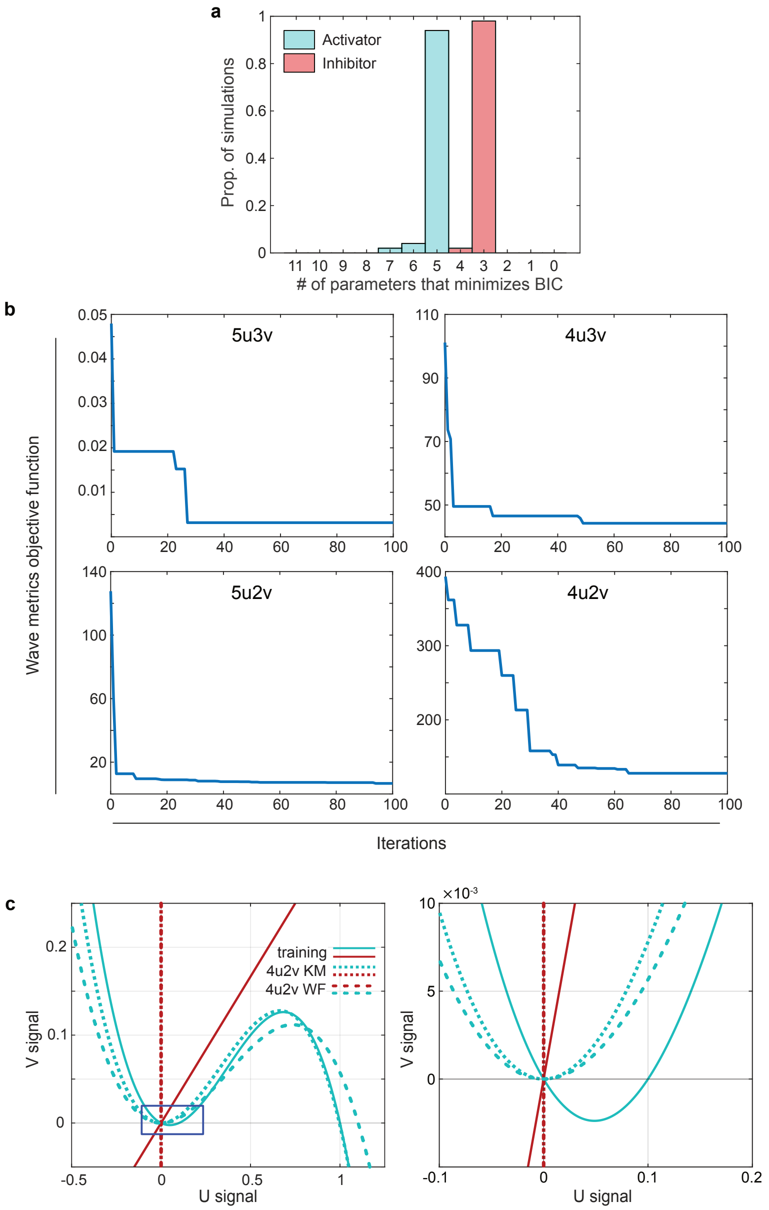


**Figure S2**: **FHN system identification using sparse KM regression and wave features optimization. a**. Distribution of the minimum of the Bayesian information criterion (BIC) computed for each simulation. **b**. Wave metrics objective function values tracked over the first 100 iterations of wave features optimization, with model structure labeled. **c**. Spatially-homogeneous nullclines plotted in the phase-plane for three models: FHN training model parameters (solid lines); 4u2v sparse-KM regression based parameters (short dashes); and 4u2v WF optimization based parameters (long dashes). The right panel is a zoomed in plot of the region depicted by the blue box in the left panel.

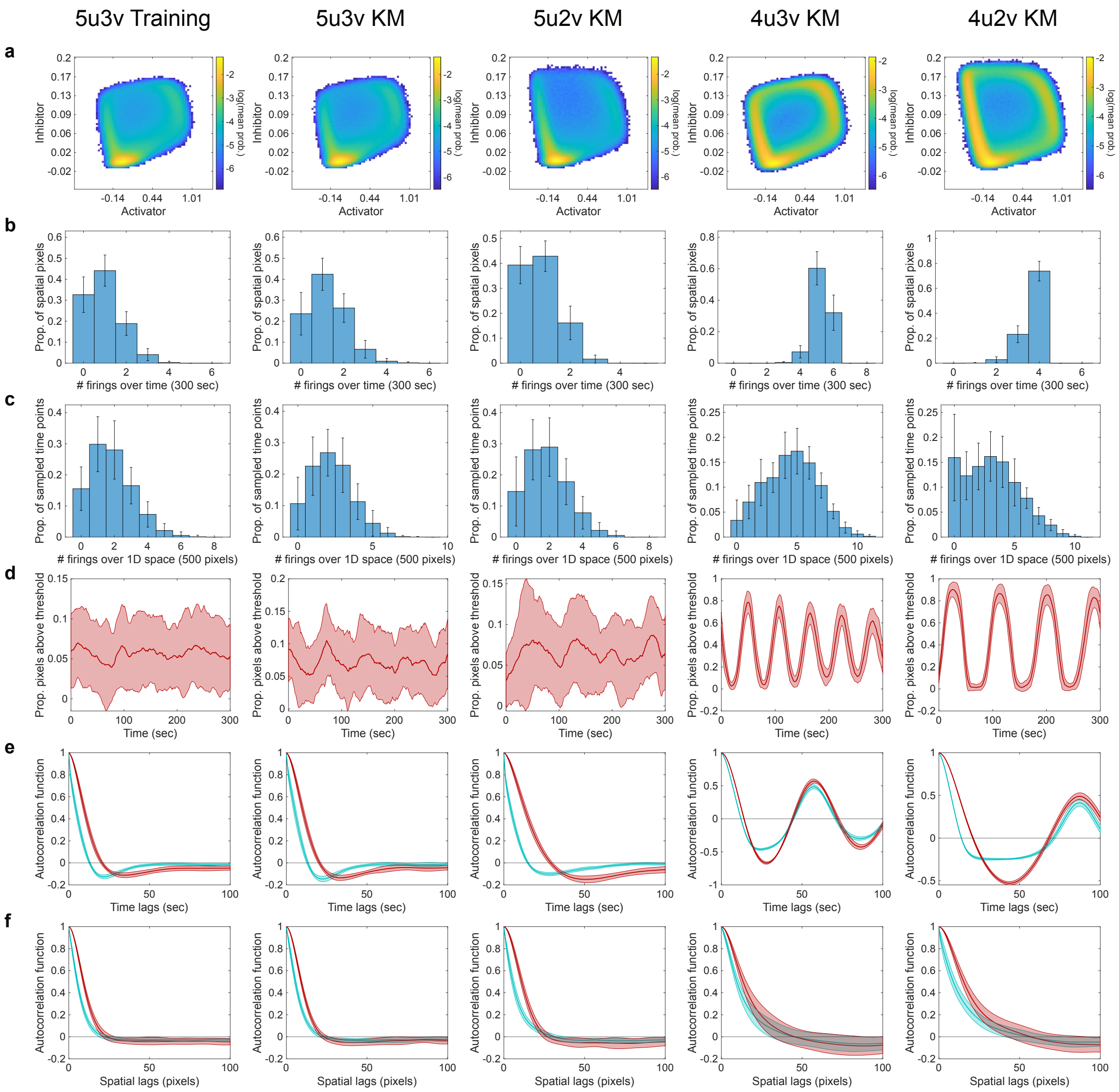


**Figure S3**: **Full spatiotemporal analysis of reconstructions of sparse FHN-1D models estimated by KM regression.** Eight wave features computed for the FHN training set (5u3v, N=50 simulations), and four polynomial reaction-diffusion model reconstruction sets using KM-based parameters (5u3v, 5u2v, 4u3v, and 4u2v; N=20 each). **a**. Joint probability distribution of the activator and inhibitor. **b**. Temporal firing frequency. **c**. Spatial firing frequency. **d**. Spatial wave activity. **e**. Temporal autocorrelation of the activator (cyan) and inhibitor (red). **f**. Spatial autocorrelation of the activator (cyan) and inhibitor (red). Shaded regions and error bars indicate mean ± standard deviation.

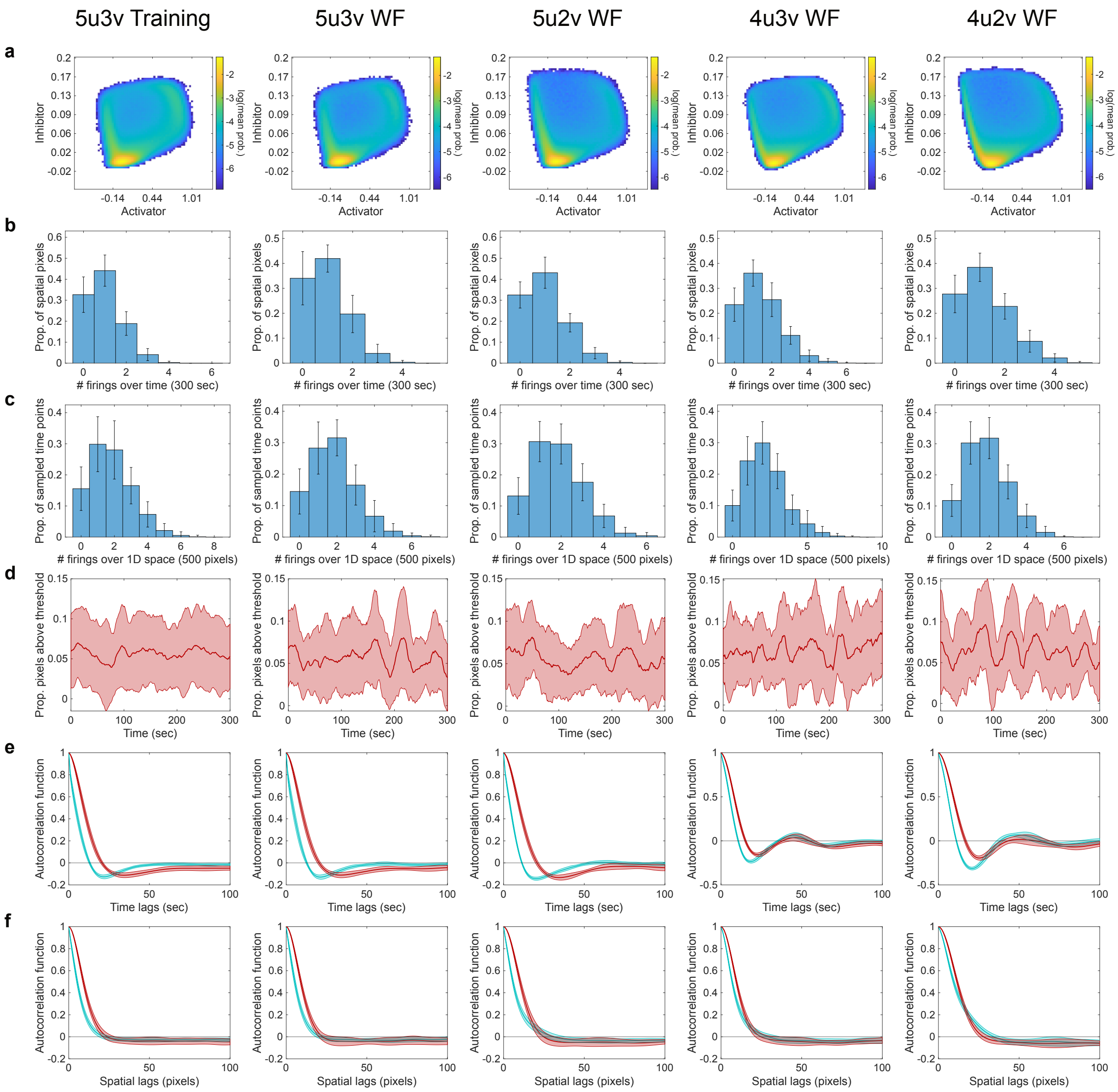


**Figure S4**: **Full spatiotemporal analysis of reconstructions of sparse FHN-1D models estimated by wave features (WF) optimization.** Eight wave features computed for the FHN training set (5u3v, N=50 simulations), and four polynomial reaction-diffusion model reconstruction sets using WF optimization based parameters (5u3v, 5u2v, 4u3v, and 4u2v; N=20 each). **a**. Joint probability distribution of the activator and inhibitor. **b**. Temporal firing frequency. **c**. Spatial firing frequency. **d**. Spatial wave activity. **e**. Temporal autocorrelation of the activator (cyan) and inhibitor (red). **f**. Spatial autocorrelation of the activator (cyan) and inhibitor (red). Shaded regions and error bars indicate mean ± standard deviation.

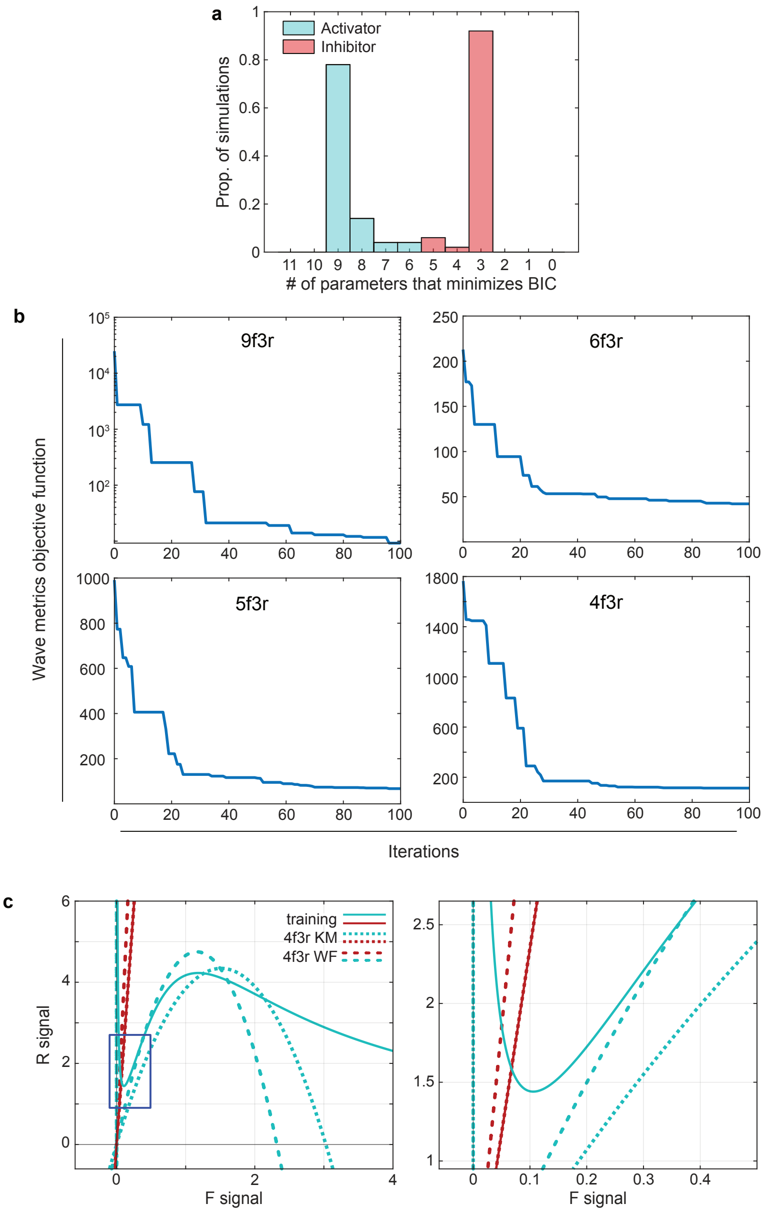


**Figure S5**: **FR system identification using sparse KM regression and wave features optimization. a**. Distribution of the minimum of the Bayesian information criterion (BIC) computed for each simulation. **b**. Wave metrics objective function values tracked over the first 100 iterations of wave features optimization, with model structure labeled. **c**. Spatially-homogeneous nullclines plotted in the phase-plane for three models: FR training model parameters (solid lines); 4f3r sparse-KM regression based parameters (short dashes); and 4f3r WF optimization based parameters (long dashes). The right panel is a zoomed in plot of the region depicted by the blue box in the left panel.

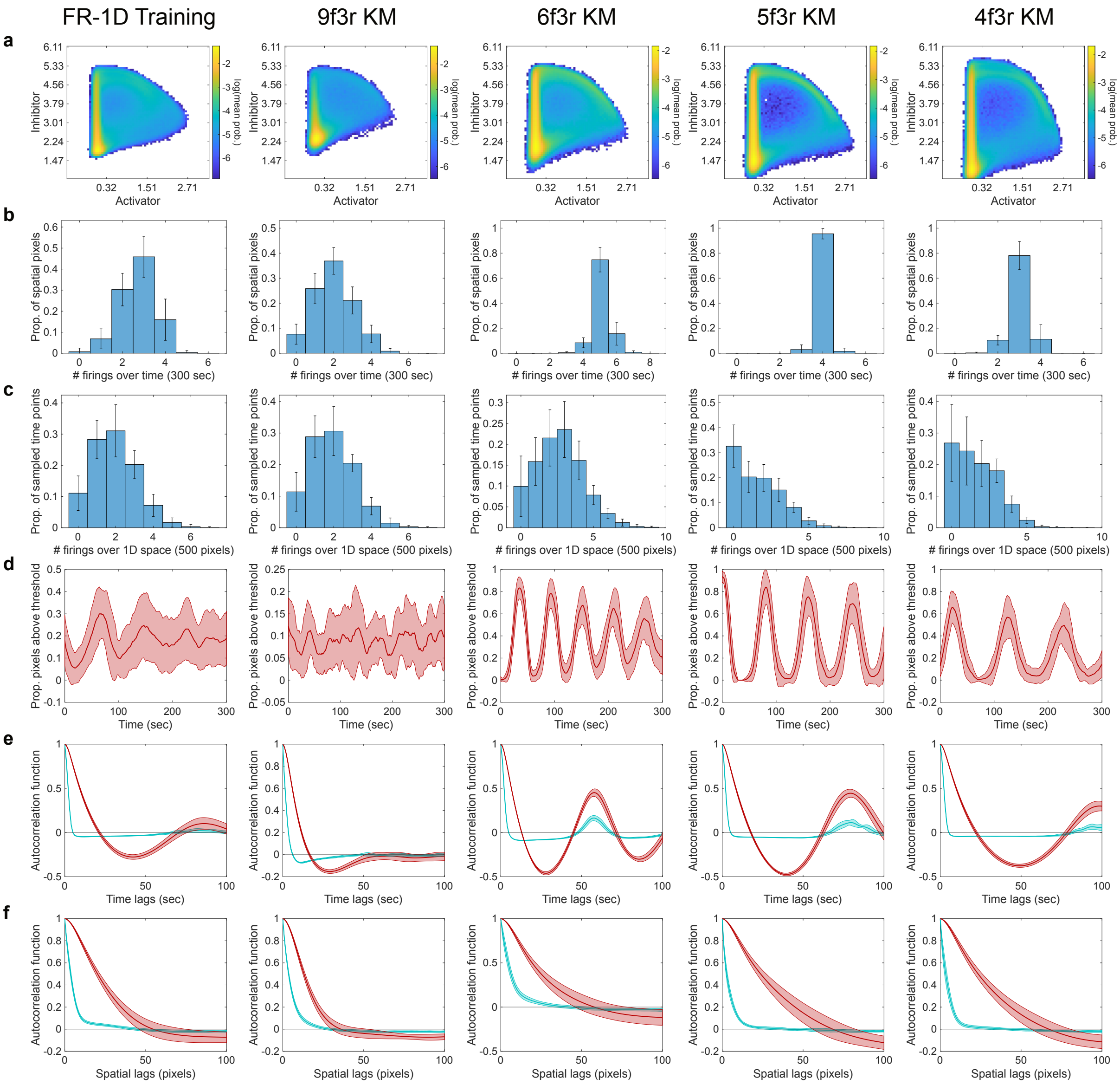


**Figure S6**: **Spatiotemporal analysis of reconstructions of sparse FR-1D models estimated by KM regression.** Eight wave features computed for the FR training set (N=50 simulations), and four polynomial reaction-diffusion model reconstruction sets using KM-based parameters (9f3r, 6f3r, 5f3r, and 4f3r; N=20 each). **a**. Joint probability distribution of the activator and inhibitor. **b**. Temporal firing frequency. **c**. Spatial firing frequency. **d**. Spatial wave activity. **e**. Temporal autocorrelation of the activator (cyan) and inhibitor (red). **f**. Spatial autocorrelation of the activator (cyan) and inhibitor (red). Shaded regions and error bars indicate mean ± standard deviation.

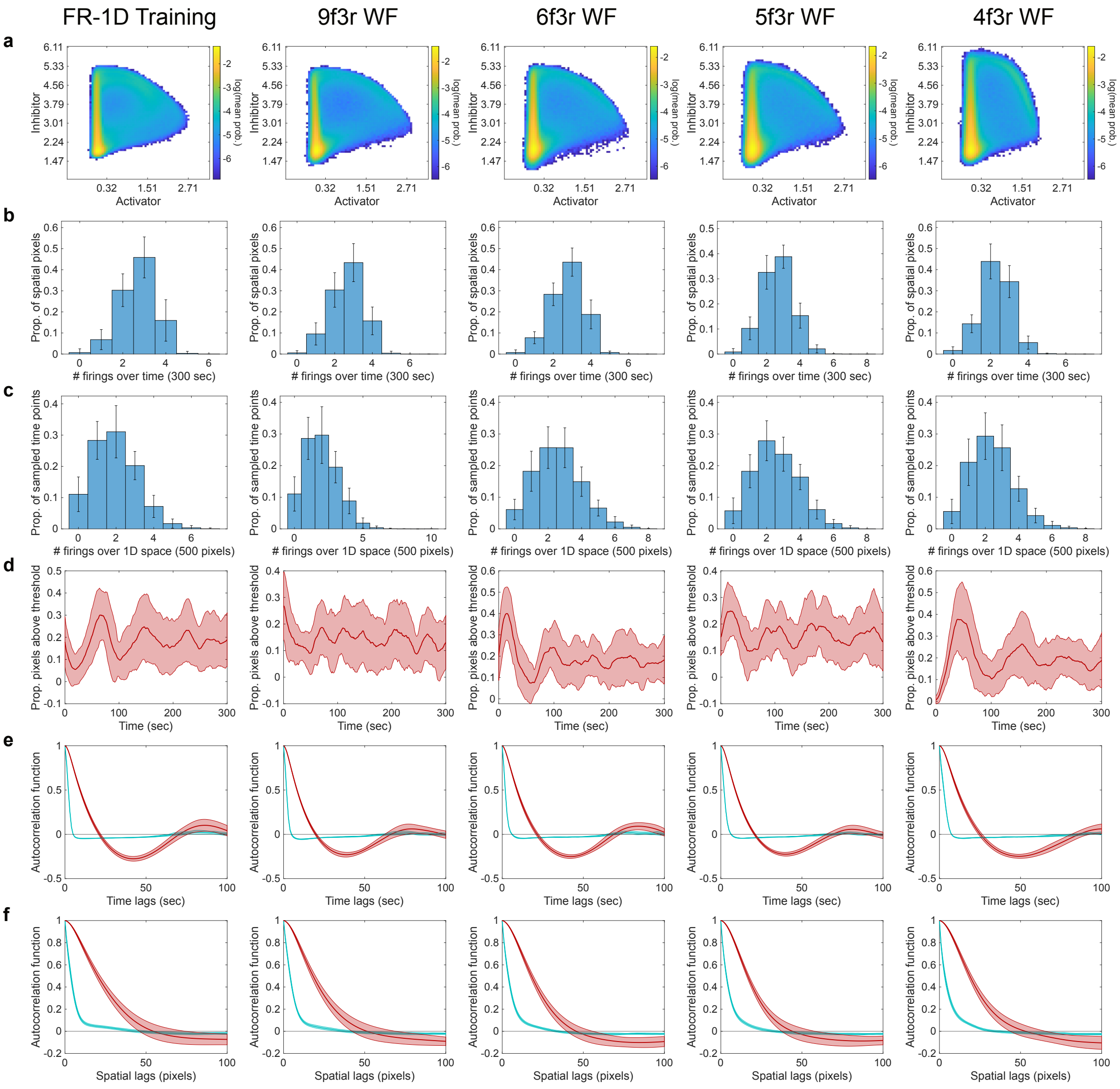


**Figure S7**: **Spatiotemporal analysis of reconstructions of sparse FR-1D models estimated by wave features (WF) optimization.** Eight wave features computed for the FR training set (N=50 simulations), and four polynomial reaction-diffusion model reconstruction sets using WF optimization based parameters (9f3r, 6f3r, 5f3r, and 4f3r; N=20 each). **a**. Joint probability distribution of the activator and inhibitor. **b**. Temporal firing frequency. **c**. Spatial firing frequency. **d**. Spatial wave activity. **e**. Temporal autocorrelation of the activator (cyan) and inhibitor (red). **f**. Spatial autocorrelation of the activator (cyan) and inhibitor (red). Shaded regions and error bars indicate mean ± standard deviation.

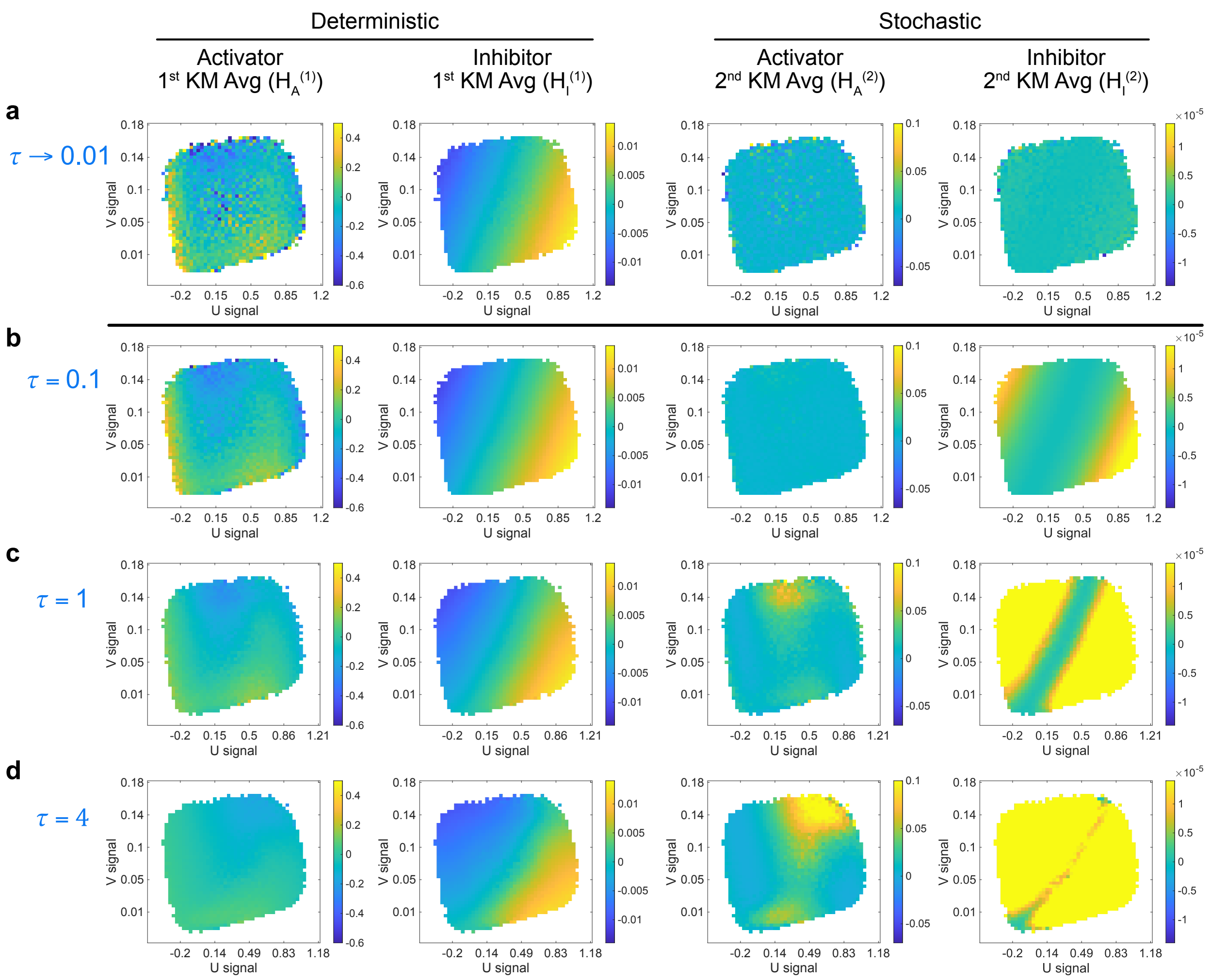


**Figure S8**: **Visualizing finite-time KM averages computed from FHN-2D data with various rates of temporal subsampling ($\tau$ = 0.1, 1, 4 s).** First and second KM averages for the activator and inhibitor computed over a Laplacian-extended state space, averaged over Laplacian values, and plotted as a heatmap in the activator-inhibitor phase plane. **a**. KM averages computed in the limit of $\tau \to 0.01$ from one FHN-2D training simulation recorded at $\tau = 0.1$ s. **b**. Finite-time KM averages computed for $\tau = 0.1$ s from one FHN-2D training simulation. **c**. Finite-time KM averages computed for $\tau = 1$ s from 10 FHN-2D training simulations. **d**. Finite-time KM averages computed for $\tau = 4$ s from 40 FHN-2D training simulations.

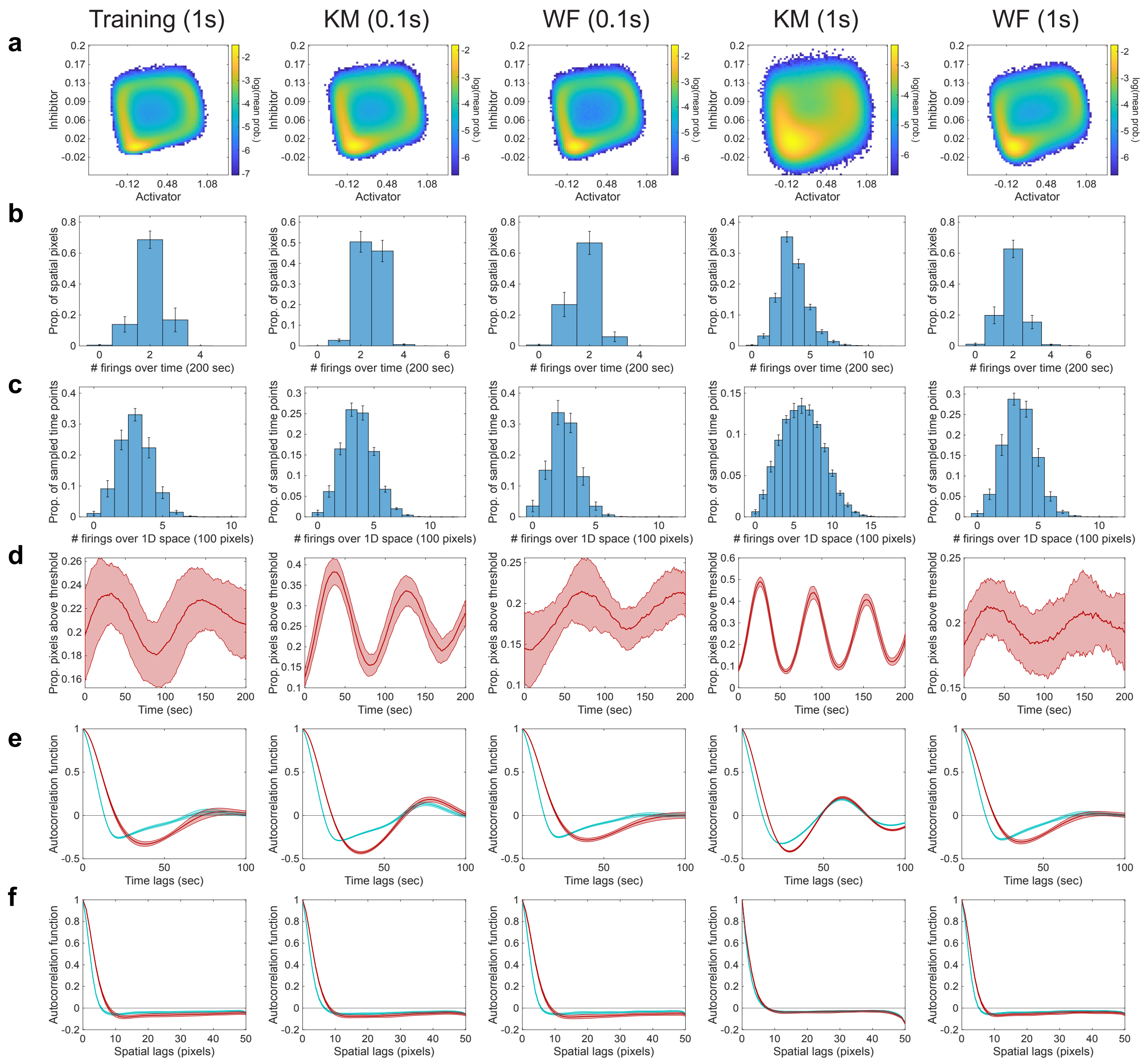


**Figure S9**: **Spatiotemporal analysis of FHN-2D reconstructions based on parameters from finite-time KM regression and wave features (WF) optimization on subsampled data (**$\tau = 0.1, 1$ **s).** Eight wave features computed for the FHN-2D training model (N=50 simulations) and reconstruction sets using finite-time KM and WF based parameters (N=20 each), all subsampled at the indicated time step, $\tau$. **a**. Joint probability distribution of the activator and inhibitor. **b**. Temporal firing frequency. **c**. Spatial firing frequency. **d**. Spatial wave activity. **e**. Temporal autocorrelation of the activator (cyan) and inhibitor (red). **f**. Spatial autocorrelation of the activator (cyan) and inhibitor (red). Shaded regions and error bars indicate mean ± standard deviation.

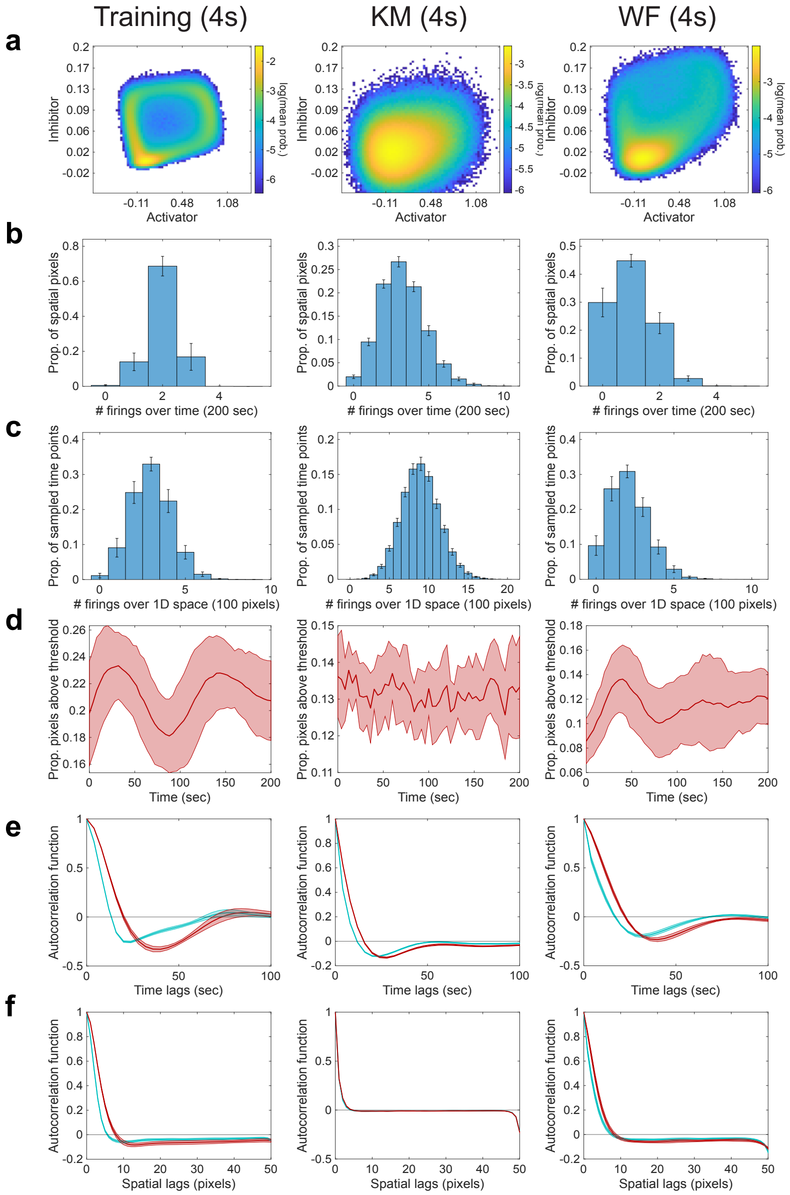


**Figure S10**: **Spatiotemporal analysis of FHN-2D reconstructions based on parameters from finite-time KM regression and wave features (WF) optimization on subsampled data ($\tau$ = 4 s).** Eight wave features computed for the FHN-2D training model (N=50 simulations) and reconstruction sets using finite-time KM and WF based parameters (N=20 each), all subsampled at the indicated time step, $\tau$. **a**. Joint probability distribution of the activator and inhibitor. **b**. Temporal firing frequency. **c**. Spatial firing frequency. **d**. Spatial wave activity. **e**. Temporal autocorrelation of the activator (cyan) and inhibitor (red). **f**. Spatial autocorrelation of the activator (cyan) and inhibitor (red). Shaded regions and error bars indicate mean ± standard deviation.

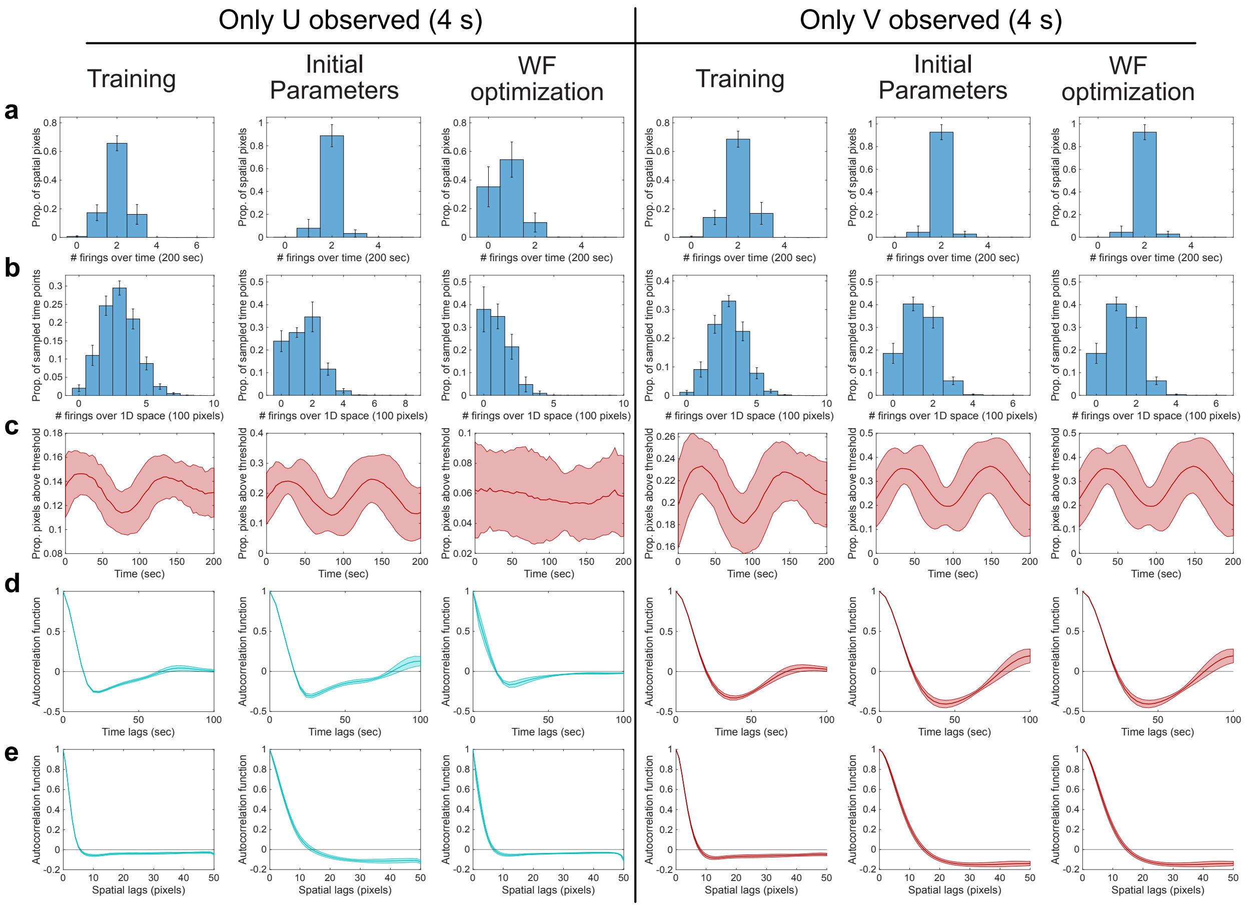


**Figure S11**: **Spatiotemporal analysis of reconstructions based on parameters from wave features (WF) optimization on subsampled** ($\tau$ = 4 s)**, partially-observed data.** Five wave features computed for the FHN-2D training model (N=50 simulations) and reconstruction sets using initial parameters and WF optimization based parameters when only observing U or only observing V (N=20 each). **a**. Temporal firing frequency. **b**. Spatial firing frequency. **c**. Spatial wave activity. **d**. Temporal autocorrelation of either only the activator (cyan), or only the inhibitor (red). **e**. Spatial autocorrelation of either only the activator (cyan) and or only the inhibitor (red). Shaded regions indicate mean ± standard deviation. Shaded regions and error bars indicate mean ± standard deviation.

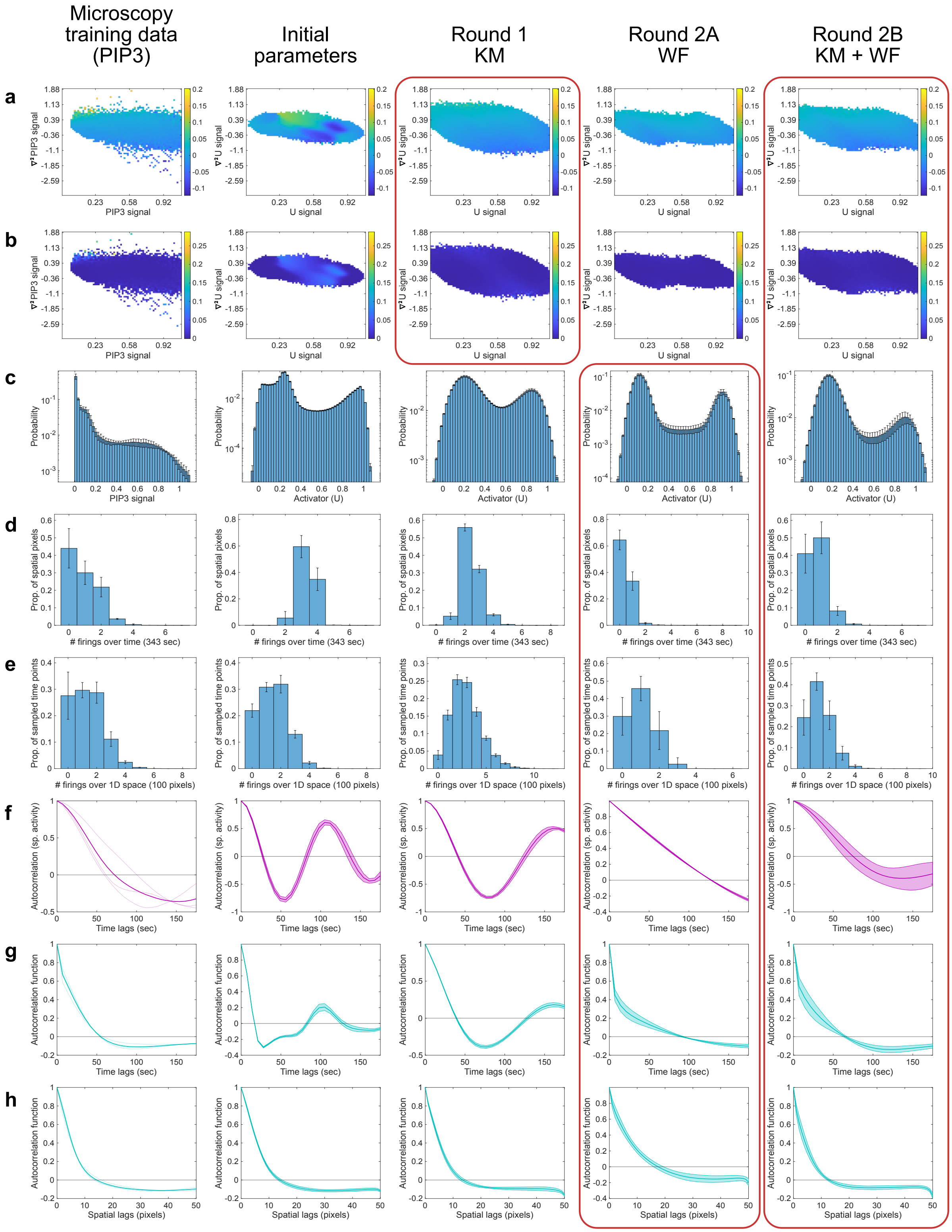

Microscopy training data (PIP3)
Initial parameters
Round 1 KM
Round 2A WF
Round 2B KM + WF
a
b
c
d
e
f
g
h
PIP3 signal
U signal
Activator (U)
### firings over time (343 sec)
### firings over 1D space (100 pixels)
Time lags (sec)
Spatial lags (pixels)
Probability
Prop. of spatial pixels
Prop. of sampled time points
Autocorrelation (sp. activity)
Autocorrelation function

**Figure S12**: **Spatiotemporal features of FHN-2D reconstructions based on parameters after optimization on low-temporal resolution ($\tau$ = 7 s) live-cell imaging data with only one observed species (PIP3).** Eight spatiotemporal features computed for PIP3 microscopy training data (single replicate, split into N=3 parts) and four reconstruction sets (N=20 simulations). The four reconstruction sets are based on: 1) initial FHN-2D parameters, 2) final parameters after round 1 optimization using projected finite-time KM averages only, 3) final parameters after round 2A optimization using only wave features (WF), and 4) final parameters after round 2B optimization using projected finite-time KM averages and wave features (KM + WF). N=20 for all reconstructions. **a**. Projected finite-time first KM averages for PIP3/activator. **b**. Projected finite-time second KM averages for PIP3/activator. **c**. Probability distribution of PIP3/activator. **d**. Temporal firing frequency distribution. **e**. Spatial firing frequency distribution. **f**. Autocorrelation of global spatial activity. **g**. Temporal autocorrelation of PIP3/activator. **h**. Spatial autocorrelation of PIP3/activator. Shaded regions and error bars indicate mean ± standard deviation. Red boxes indicate features that were optimized for at each round.

#### Supplementary Tables

**Table S1**: **Estimated FHN-1D model structures and parameters from sparse KM regression and wave features optimization.**

Estimated polynomial stochastic reaction-diffusion equation (SRDE) parameters for FHN-1D data using sparse Kramers-Moyal (KM) regression and wave features (WF) optimization. Standard deviation (SD) with N=50 for KM-based parameters. Note: $x_n$ refers to the $n$-th species in the set $\{u, v\}$.

General SRDE: $dx_n = D_n\nabla^2 x_n + c_1 + c_2u + c_3v + c_4u^2 + c_5uv + c_6v^2 + c_7u^3 + c_8u^2v + c_9uv^2 + c_{10}v^3 + g_n dQ_n$

| | Activator SRDE terms and parameters | | | | | | Inhibitor SRDE terms and parameters | | | |
|---|---|---|---|---|---|---|---|---|---|---|
| | $\nabla^2 u$ | $u$ | $v$ | $u^2$ | $u^3$ | $dQ_u$ | $\nabla^2 v$ | $u$ | $v$ | $dQ_v$ |
| **Models** | $D_u$ | $a_2$ | $a_3$ | $a_4$ | $a_7$ | $g_u$ | $D_v$ | $b_2$ | $b_3$ | $g_v$ |
| **Ground truth** | 1.4 | -0.2 | -2 | 2.2 | -2 | 0.1 | 1.2 | 0.02 | -0.06 | 0 |
| **5u3v** | | | | | | | | | | |
| KM | 1.3823 | -0.1934 | -1.9839 | 2.1634 | -1.9689 | 0.1026 | 1.1943 | 0.0200 | -0.0599 | $6.29\times10^{-4}$ |
| (SD) | 0.0353 | 0.0251 | 0.0698 | 0.0894 | 0.0910 | 0.0015 | 0.0054 | $2.22\times10^{-5}$ | $1.31\times10^{-4}$ | $2.49\times10^{-5}$ |
| WF | 1.4017 | -0.1933 | -1.9938 | 2.1664 | -1.9661 | 0.0996 | 1.2103 | 0.0201 | -0.0590 | $7.04\times10^{-5}$ |
| **5u2v** | | | | | | | | | | |
| KM | 1.3823 | -0.1934 | -1.9839 | 2.1634 | -1.9689 | 0.1026 | 1.8601 | 0.0142 | – | $6.29\times10^{-4}$ |
| (SD) | 0.0353 | 0.0251 | 0.0698 | 0.0894 | 0.0910 | 0.0015 | 0.0649 | $2.68\times10^{-4}$ | – | $2.49\times10^{-5}$ |
| WF | 1.4338 | -0.1354 | -2.3847 | 2.1085 | -1.9126 | 0.0938 | 2.0688 | 0.0180 | – | $4.53\times10^{-4}$ |
| **4u3v** | | | | | | | | | | |
| KM | 1.4198 | – | -1.9318 | 1.6749 | -1.6798 | 0.1026 | 1.1943 | 0.0200 | -0.0599 | $6.29\times10^{-4}$ |
| (SD) | 0.0359 | – | 0.0713 | 0.0799 | 0.0938 | 0.0015 | 0.0054 | $2.22\times10^{-5}$ | $1.31\times10^{-4}$ | $2.49\times10^{-5}$ |
| WF | 1.3349 | – | -2.3356 | 1.6936 | -1.5933 | 0.0568 | 1.8138 | 0.0225 | -0.0510 | $4.83\times10^{-4}$ |
| **4u2v** | | | | | | | | | | |
| KM | 1.4198 | – | -1.9318 | 1.6749 | -1.6798 | 0.1026 | 1.8601 | 0.0142 | – | $6.29\times10^{-4}$ |
| (SD) | 0.0359 | – | 0.0713 | 0.0799 | 0.0938 | 0.0015 | 0.0649 | $2.68\times10^{-4}$ | – | $2.49\times10^{-5}$ |
| WF | 1.5507 | – | -2.2421 | 1.3954 | -1.2667 | 0.0521 | 2.2159 | 0.0175 | – | $6.89\times10^{-5}$ |

**Table S2**: **Estimated FR-1D model structures and parameters from sparse KM regression and wave features optimization.**

Estimated polynomial reaction-diffusion SRDE parameters for FR-1D data using sparse Kramers-Moyal (KM) regression and wave features (WF) optimization. Standard deviation (SD) with N=50 for KM-based parameters. Note: $x_n$ refers to the $n$-th species in the set $\{f, r\}$.

General SRDE: $dx_n = D_n \nabla^2 x_n + c_1 + c_2 f + c_3 r + c_4 f^2 + c_5 fr + c_6 r^2 + c_7 f^3 + c_8 f^2 r + c_9 fr^2 + c_{10} r^3 + g_n dQ_n$

| | Activator SRDE terms and parameters | | | | | | | | | | Inhibitor SRDE terms and parameters | | | |
|---|---|---|---|---|---|---|---|---|---|---|---|---|---|---|
| | $\nabla^2 f$ | $f$ | $f^2$ | $fr$ | $r^2$ | $f^3$ | $f^2 r$ | $fr^2$ | $r^3$ | $dQ_f$ | $\nabla^2 r$ | $f$ | $r$ | $dQ_r$ |
| **Models** | $D_f$ | $a_2$ | $a_4$ | $a_5$ | $a_6$ | $a_7$ | $a_8$ | $a_9$ | $a_{10}$ | $g_f$ | $D_r$ | $b_2$ | $b_3$ | $g_r$ |
| **9f3r** | | | | | | | | | | | | | | |
| KM | 2.7461 | 0.7979 | 3.5613 | -0.7928 | 0.0160 | -0.9996 | -0.1868 | 0.0327 | -0.0037 | 0.1610 | 5.1973 | 0.5872 | -0.0250 | 0.0360 |
| (SD) | 0.0255 | 0.1612 | 0.0995 | 0.0977 | 0.0015 | 0.0250 | 0.0152 | 0.0132 | $3.45 \times 10^{-4}$ | 0.0012 | 0.0044 | $2.10 \times 10^{-4}$ | $3.80 \times 10^{-5}$ | $7.19 \times 10^{-4}$ |
| WF | 3.1791 | 0.6792 | 3.7542 | -0.8726 | 0.0154 | -0.9454 | -0.1884 | 0.0327 | -0.0039 | 0.1618 | 4.6107 | 0.5826 | -0.0251 | 0.0372 |
| **6f3r** | | | | | | | | | | | | | | |
| KM | 2.7651 | 0.7683 | 3.3639 | -0.6429 | – | -0.9902 | -0.1456 | – | – | 0.1610 | 5.1973 | 0.5872 | -0.0250 | 0.0360 |
| (SD) | 0.0255 | 0.0792 | 0.0880 | 0.0187 | – | 0.0230 | 0.0142 | – | – | 0.0012 | 0.0044 | $2.10 \times 10^{-4}$ | $3.80 \times 10^{-5}$ | $7.19 \times 10^{-4}$ |
| WF | 3.0371 | 0.7367 | 3.4607 | -0.6944 | – | -0.9675 | -0.1498 | – | – | 0.1681 | 5.6612 | 0.6083 | -0.0161 | 0.0369 |
| **5f3r** | | | | | | | | | | | | | | |
| KM | 2.7687 | – | 4.0124 | -0.4680 | – | -1.0536 | -0.2720 | – | – | 0.1610 | 5.1973 | 0.5872 | -0.0250 | 0.0360 |
| (SD) | 0.0257 | – | 0.0495 | 0.0088 | – | 0.0201 | 0.0076 | – | – | 0.0012 | 0.0044 | $2.10 \times 10^{-4}$ | $3.80 \times 10^{-5}$ | $7.19 \times 10^{-4}$ |
| WF | 2.6033 | – | 3.8783 | -0.4006 | – | -0.9802 | -0.3105 | – | – | 0.1990 | 6.3614 | 0.6363 | -0.0183 | 0.0352 |
| **4f3r** | | | | | | | | | | | | | | |
| KM | 2.6751 | – | 3.8016 | -0.6631 | – | -1.2554 | – | – | – | 0.1610 | 5.1973 | 0.5872 | -0.0250 | 0.0360 |
| (SD) | 0.0253 | – | 0.0483 | 0.0069 | – | 0.0204 | – | – | – | 0.0012 | 0.0044 | $2.10 \times 10^{-4}$ | $3.80 \times 10^{-5}$ | $7.19 \times 10^{-4}$ |
| WF | 2.2441 | – | 4.3097 | -0.5260 | – | -1.8584 | – | – | – | 0.1961 | 7.3452 | 0.5568 | -0.0151 | 0.0281 |

**Table S3**: **Estimated FHN-2D model parameters from finite-time KM regression and wave features optimization on subsampled, fully-observed data** ($\tau = 0.1, 1, 4$ **s).**

Estimated stochastic reaction-diffusion equation (SRDE) parameters for the FHN-2D model using finite-time Kramers-Moyal (FT-KM) regression and wave features (WF) optimization with varying sampling time steps ($\tau$).

Activator SRDE: $du = D_u \nabla^2 u + a_1 u + a_2 v + a_3 u^2 + a_4 u^3 + g_u dQ_u$

Inhibitor SRDE: $dv = D_v \nabla^2 v + b_1 u + b_2 v + g_v dQ_v$

| | Activator SRDE parameters | | | | | | Inhibitor SRDE parameters | | | |
|---|---|---|---|---|---|---|---|---|---|---|
| **Models** | $D_u$ | $a_1$ | $a_2$ | $a_3$ | $a_4$ | $g_u$ | $D_v$ | $b_1$ | $b_2$ | $g_v$ |
| Ground truth | 0.13 | -0.2 | -2 | 2.2 | -2 | 0.0725 | 0.00625 | 0.015 | -0.045 | 0 |
| $\tau = 0.1$ s | | | | | | | | | | |
| FT-KM | 0.1343 | -0.1833 | -2.0303 | 2.1201 | -1.9158 | 0.0831 | 0.0068 | 0.0150 | -0.0452 | 0.0020 |
| WF | 0.1479 | -0.1931 | -1.9242 | 2.1012 | -1.8905 | 0.0727 | 0.0068 | 0.0151 | -0.0464 | 0.0019 |
| $\tau = 1$ s | | | | | | | | | | |
| FT-KM | 0.1091 | -0.1170 | -1.6788 | 1.4589 | -1.2793 | 0.1274 | 0.0104 | 0.0149 | -0.0484 | 0.0060 |
| WF | 0.1093 | -0.1279 | -1.9406 | 1.5803 | -1.2971 | 0.0611 | 0.0121 | 0.0149 | -0.0638 | 0.0038 |
| $\tau = 4$ s | | | | | | | | | | |
| FT-KM | 0.0664 | -0.0619 | -0.9560 | 0.5687 | -0.4633 | 0.1667 | 0.0169 | 0.0139 | -0.0558 | 0.0115 |
| WF | 0.1521 | -0.0456 | -1.0370 | 0.7663 | -0.5732 | 0.1153 | 0.0214 | 0.0136 | -0.0467 | 0.0033 |

**Table S4**: **Estimated FHN-2D model parameters from wave features optimization on subsampled, partially-observed data** ($\tau = 4$ **s).**

Estimated stochastic reaction-diffusion equation (SRDE) parameters for the FHN-2D model using wave features (WF) optimization with only one of the two species observed at a sampling time step of $\tau = 4$ s. Initial parameters indicate the parameters at the start of optimization.

Activator SRDE: $du = D_u \nabla^2 u + a_1 u + a_2 v + a_3 u^2 + a_4 u^3 + g_u dQ_u$

Inhibitor SRDE: $dv = D_v \nabla^2 v + b_1 u + b_2 v + g_v dQ_v$

| | Activator SRDE parameters | | | | | | Inhibitor SRDE parameters | | | |
|---|---|---|---|---|---|---|---|---|---|---|
| **Models** | $D_u$ | $a_1$ | $a_2$ | $a_3$ | $a_4$ | $g_u$ | $D_v$ | $b_1$ | $b_2$ | $g_v$ |
| Ground truth | 0.13 | -0.2 | -2 | 2.2 | -2 | 0.0725 | 0.00625 | 0.015 | -0.045 | 0 |
| Initial parameters | 0.26 | -0.02 | -2 | 2.02 | -2 | 0.0435 | 0.25 | 0.015 | -0.045 | 0 |
| WF: observe only U | 0.2552 | -0.0211 | -2.0679 | 1.9604 | -2.0303 | 0.0434 | 0.2482 | 0.0155 | -0.0451 | $2.24 \times 10^{-5}$ |
| WF: observe only V | 0.2625 | -0.0209 | -2.0542 | 1.9221 | -2.0310 | 0.0432 | 0.2529 | 0.0152 | -0.0457 | $1.39 \times 10^{-5}$ |

**Table S5**: **Estimated FHN-2D model parameters from projected finite-time KM averages and wave features optimization on live-cell imaging data of a single observed species (PIP3) measured every $\tau = 7$ s.**

Estimated stochastic reaction-diffusion equation (SRDE) parameters for the FHN-2D model using projected finite-time Kramers-Moyal (KM) and wave features (WF) optimization on live-cell imaging data of PIP3 (activator). Data was acquired at a sampling rate of $\tau = 7$ s for 150 frames. Initial parameters indicate the parameters at the start of optimization.

Activator SRDE: $du = D_u \nabla^2 u + a_1 u + a_2 v + a_3 u^2 + a_4 u^3 + g_u dQ_u$

Inhibitor SRDE: $dv = D_v \nabla^2 v + b_1 u + b_2 v + g_v dQ_v$

| | Activator SRDE parameters | | | | | | Inhibitor SRDE parameters | | | |
|---|---|---|---|---|---|---|---|---|---|---|
| **Models** | $D_u$ | $a_1$ | $a_2$ | $a_3$ | $a_4$ | $g_u$ | $D_v$ | $b_1$ | $b_2$ | $g_v$ |
| Initial parameters | 0.26 | -0.02 | -2 | 2.02 | -2 | 0.0435 | 0.25 | 0.015 | -0.045 | 0 |
| Round 1 (KM) | 0.2735 | -0.0206 | -1.8755 | 1.4200 | -2.7579 | 0.0529 | 0.2787 | 0.0046 | -0.0558 | $8.72 \times 10^{-4}$ |
| Round 2A (WF) | 0.2484 | -0.0196 | -2.1540 | 1.2917 | -2.4113 | 0.0283 | 0.4495 | 0.0042 | -0.0702 | $8.25 \times 10^{-4}$ |
| Round 2B (KM + WF) | 0.2891 | -0.0218 | -1.9978 | 1.4383 | -2.7277 | 0.0406 | 0.3007 | 0.0046 | -0.0586 | $9.23 \times 10^{-4}$ |

#### Other Supplementary Files

**Movie S1. Representative FHN-2D model simulations of training and reconstruction data from KM regression.** Animated version of Fig. 3e.

**Movie S2. Representative FR-2D model simulations of training and reconstruction data from KM regression.** Animated version of Fig. 4e.

**Movie S3. Representative FHN-2D model reconstructions based on parameters from finite-time KM regression and wave features optimization on subsampled, fully-observed training data ($\tau$ = 1 s).** Animated version of Fig. 7b, top panel.

**Movie S4. Representative FHN-2D model reconstructions based on parameters from finite-time KM regression and wave features optimization on subsampled, fully-observed training data ($\tau$ = 4 s).** Animated version of Fig. 7b, top panel.

**Movie S5. Representative FHN-2D model reconstructions based on parameters from wave features optimization on subsampled training data with only the activator (U) observed ($\tau$ = 4 s).** Animated version of Fig. 7e.

**Movie S6. Representative FHN-2D model reconstructions based on parameters from wave features optimization on subsampled training data with only the inhibitor (V) observed ($\tau$ = 4 s).** Animated version of Fig. 7e.

**Movie S7. Representative FHN-2D model reconstructions based on parameters from spatiotemporal features optimization on live-cell imaging data of a single observed species (PIP3) measured every $\tau$ = 7 s.** Animated version of Fig. 8c. Original PIP3 training data with 150 frames are shown together with the activator signal from four reconstructions over the same time interval using 1) initial FHN-2D parameters, 2) final parameters after round 1 optimization using projected finite-time KM averages only, 3) final parameters after round 2A optimization using only wave features (WF), and 4) final parameters after round 2B optimization using projected finite-time KM averages and wave features (KM + WF).
